# Neutrophils orchestrate pre-metastatic immunosuppressive bone marrow remodeling during prostate cancer metastasis

**DOI:** 10.64898/2026.10.08.757800

**Authors:** Yingkui Tang, Xuan Zhou, Duocai Li, Yadan Liu, Guanghui Zhu, Jian Wang, Xiumei Wu, Da Zhong, Ziyi Li, Haojie Huang, Bin-Zhi Qian, Yuanyuan Zhang, Jianru Xiao, Zhiying Yue, Sujun Chen, Shancheng Ren

**Affiliations:** West China School of Public Health and West China Fourth Hospital, and State Key Laboratory of Biotherapy, Sichuan University; Chengdu, 610041, P.R. China; Department of Urology, Shanghai Changzheng Hospital, Naval Medical University; Shanghai, 200003, P.R. China; Department of Urology, The Affiliated Hospital of Jiangnan University; Wuxi, 214122, P.R. China; Jiangsu Cancer Hospital, Jiangsu Institute of Cancer Research, The Affiliated Cancer Hospital of Nanjing Medical University; Nanjing, 211166, P.R. China; National Key Laboratory of Immunity and Inflammation, Suzhou Institute of Systems Medicine, Chinese Academy of Medical Sciences & Peking Union Medical College; Suzhou, 215123, P.R. China; Department of Urology, Institute of Urologic Science and Technology, The First Affiliated Hospital, Zhejiang University School of Medicine; Hangzhou, 310003, P.R. China; Fudan University Shanghai Cancer Center, Shanghai Medical College, Center for Integrative Spatial-Omics Research, The Human Phenome Institute, Zhangjiang-Fudan International Innovation Center, Fudan University; Shanghai, 200433, P.R. China; State Key Laboratory of Molecular Oncology, National Cancer Center/National Clinical Research Center for Cancer/Cancer Hospital, Chinese Academy of Medical Sciences and Peking Union Medical College; Beijing, 100021, P.R. China; Department of Orthopedic Oncology, The Second Affiliated Hospital of Naval Medical University; Shanghai, 200003, P.R. China; Precision Research Center for Refractory Diseases, Shanghai General Hospital, Shanghai Jiaotong University, School of Medicine; Shanghai, 200240, P.R. China

**Author notes:** Corresponding author. (S.R.), (S.C.), (Z.Y.), (J.X.).

**Keywords:** Prostate cancer, Bone metastasis, Pre-metastatic niche, Tumor-associated neutrophils, IGF1, Immunosuppression

## Abstract

The dynamic reprogramming of the bone marrow (BM) microenvironment during metastatic progression is poorly defined. Through single-cell profiling of BM samples spanning benign conditions, high-risk bone metastatic potential localized prostate cancer (HMP) and established bone metastatic prostate cancer (PCa), we uncovered a primary tumor-to-BM signaling axis that orchestrates niche formation through neutrophil-mediated remodeling. Tumor-associated neutrophils (TANs) emerge during the HMP stage, persist throughout metastatic progression, correlate with poor patient prognosis, and exhibit direct immunosuppressive activity that contributes to a BM environment permissive to metastatic colonization. Functional and genetic studies showed that tumor-associated systemic IGF1 reprogrammed BM neutrophils through IGF1R-MAPK/ERK-KLF4 signaling, with IL-1β, reactive oxygen species and nitric oxide contributing to TAN-mediated T cell suppression. Tumor-cell Igf1 perturbation, IGF1R blockade and neutrophil-specific *Igf1r* deletion attenuated TAN-associated BM remodeling, and reduced subsequent bone metastatic progression. In bone-metastasis models, combined IGF1R and PD-1 blockade suppressed bone-metastatic progression. Translationally, our investigator-initiated clinical trial (NCT06866548) of teprotumumab plus tislelizumab in patients with established metastatic castration-resistant prostate cancer (mCRPC) provided initial safety data and exploratory clinical observations. Together, these findings identify IGF1R-dependent neutrophil reprogramming as a functionally important component of the evolving PCa bone metastatic microenvironment and provide a rationale for therapeutic targeting of IGF1R signaling together with immune checkpoint blockade.

## Main Text

Distant metastasis to bone is the primary driver of mortality in prostate cancer (PCa), the second most common cancer in men worldwide ^1–3^. This lethal progression, characterized by the dissemination of tumor cells to and colonization of the skeleton, occurs in up to 80% of men with advanced disease ^4,5^. The clinical consequences are stark: the five-year survival rate plummets from over 97% for localized disease to approximately 32% for patients with bone metastases ^5^. Beyond its devastating impact on survival, bone metastatic disease is accompanied by debilitating skeletal-related events, including severe pain and pathological fractures, which profoundly impair quality of life ^6^.

Characteristics of bone metastases have been partly resolved, which is largely framed by the classic “vicious cycle” paradigm—a reciprocal dialogue among tumor cells, osteoblasts and osteoclasts ^7–12^. Recent studies revealed the involvement of other cell types, such as neutrophils and macrophages, in metastatic PCa (mPCa) progression ^13–15^, which highlights the complex reprogramming in the metastatic BM niche. Thus, more comprehensive studies are needed to delineate the dynamic reprogramming of BM during bone metastasis. Beside the metastatic niche itself, the pre-metastatic niche (PMN)—a microenvironment systemically reprogrammed to support future colonization—has emerged as a critical paradigm in metastasis to organs such as the lung and liver ^16,17^. Previous studies have demonstrated the PMN formation in breast cancer bone metastasis using mouse models ^18,19^, mediated by osteoclasts, and pro-osteolytic T and B cells. However, whether analogous BM alterations are already present in patients with PCa who are at high metastatic risk but lack pathologically detectable bone metastasis, which cellular states characterize this setting, and how these alterations relate to the established metastatic niche remain poorly defined. A multi-stage, high-resolution cellular atlas is therefore required to delineate the metastatic cascade, from the earliest niche preparations to the complex ecosystem of the established lesion.

Here, through single-cell transcriptional profiling of human BM across non-malignant benign (benign), clinically high-risk, without pathologically confirmed bone metastasis (HMP), and metastatic stages, we characterize the evolution of BM remodeling during PCa progression. We identify a previously unrecognized population of CCR3^+^ tumor-associated neutrophils (TANs) that emerges during the HMP stage and persists in metastatic BM, and is associated with tumor-related systemic insulin-like growth factor 1 (IGF1), which correlates with poor patient prognosis. In preclinical models, systemic IGF1 induces key features of this BM state and reprograms neutrophils toward a TAN state via the MAPK/ERK-KLF4 axis that contributes to T cell dysfunction through IL-1β-, ROS-and NO-associated mechanisms. Importantly, IGF1R blockade or neutrophil-specific *Igf1r* deletion attenuated TAN-associated BM remodeling, and combined IGF1R blockade with anti-PD-1 therapy reduced bone metastatic progression in vivo. An exploratory investigator-initiated clinical trial further evaluated the safety and preliminary clinical activity of combined IGF1R and PD-1 blockade in patients with metastatic castration-resistant prostate cancer (mCRPC). Collectively, these findings support a model in which tumor-associated systemic IGF1 contributes to TAN-associated immunosuppressive BM remodeling and provide a rationale for further evaluation of IGF1R blockade together with immune checkpoint inhibition in advanced PCa.

### A single-cell atlas reveals dynamic bone marrow microenvironment remodeling during PCa progression

To delineate the dynamic remodeling of the BM microenvironment during PCa bone metastasis, we performed single-cell RNA-sequencing (scRNA-seq) on BM samples from distinct stages of disease progression (main cohort): non-malignant benign (Benign; n=6), High-risk bone metastatic potential localized PCa (HMP; n=7), and metastatic PCa (Mets; n=7) (Figure 1a; Table S1). Benign BM samples were obtained from the lumbar vertebrae of patients with non-malignant conditions, such as lumbar disc herniation and lumbar spinal stenosis. The HMP group comprised patients with highly aggressive, locally advanced PCa (unfavorable intermediate-risk, n=1; very high-risk, n=6) who were initially suspected of having bone metastases based on imaging findings ^20^. Although targeted biopsies of these suspicious lesions were pathologically negative for tumor cells at the time of collection, updated clinical follow-up revealed that three of these patients subsequently developed radiologically confirmed bone metastases (Table S1). For the remaining 4 patients who have not yet presented with overt metastasis, their current follow-up duration is approximately 2 years, and longer follow-up will be required to determine their subsequent metastatic status. Therefore, these samples provide a transcriptomic snapshot of the BM microenvironment in patients at high risk for metastasis but without confirmed tumor cell seeding. In the Mets group, BM samples were collected via needle biopsy or surgery from patients diagnosed with PCa bone metastasis. Samples were subjected to scRNA-seq using the BD Rhapsody platform (Methods).

**Fig. 1.**
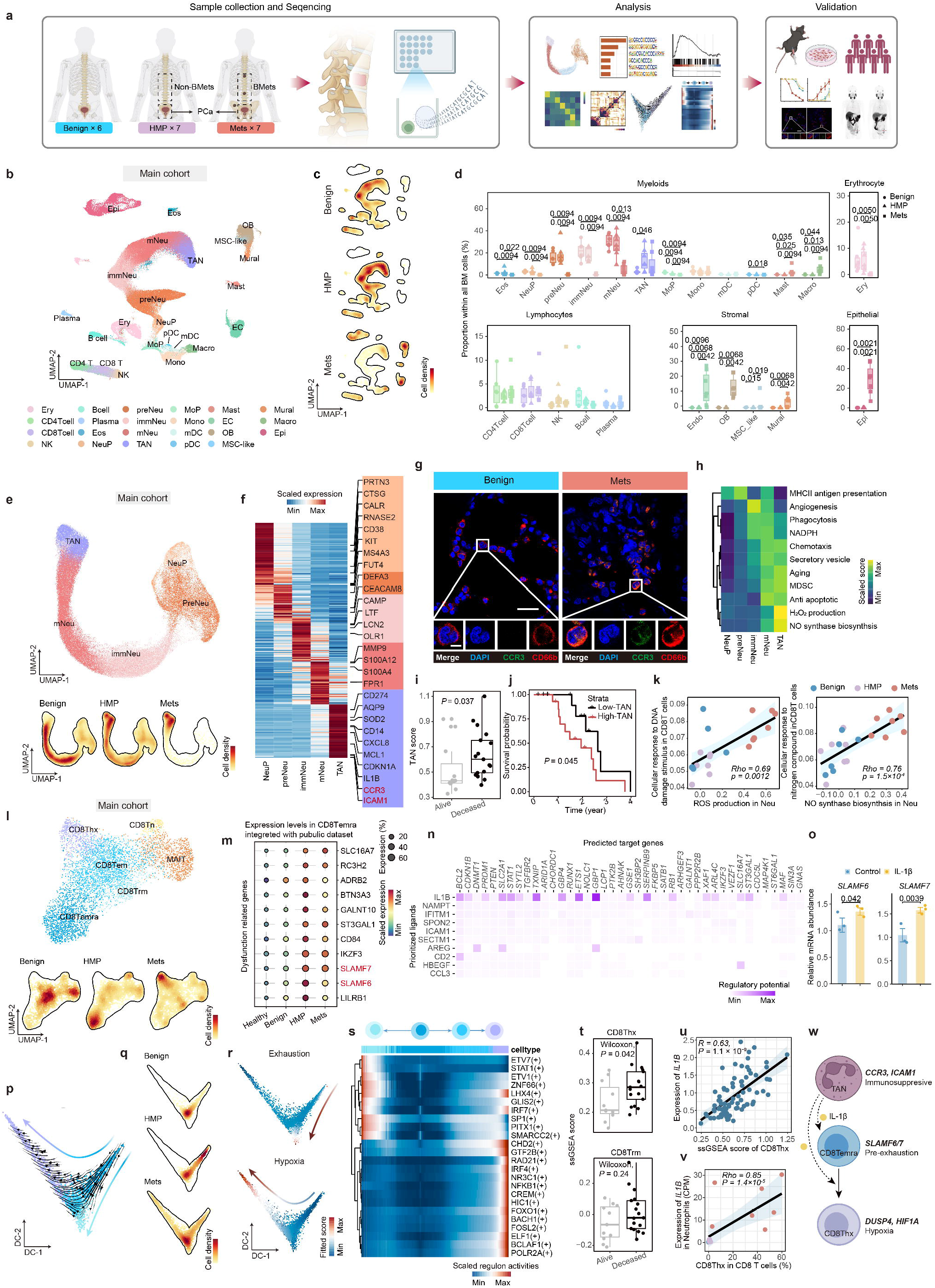
Single-cell profiling identifies TAN-associated immune remodeling in HMP bone marrow. (a) Schematic of bone marrow (BM) collection, single-cell RNA sequencing and downstream validation in the Benign (n = 6), HMP (n = 7) and Mets (n = 7) cohorts. HMP denotes patients without detectable bone metastasis but at high risk of metastatic progression; Mets denotes prostate cancer bone metastasis. (b) UMAP of all cells in the main cohort, colored by major cell type. (c) Cell-density maps on the integrated UMAP by disease group. (d) Proportions of major myeloid, lymphoid, stromal, erythroid and epithelial populations across groups. Statistical comparisons were performed using patient-level cell proportions. Values above comparisons show adjusted *P* values. (e) UMAP of neutrophil subpopulations and corresponding density maps by disease group. (f) Heatmap of differentially expressed genes across neutrophil states; values are row-scaled average expression. (g) Representative immunofluorescence images of benign and metastatic BM stained for DAPI, CCR3 and CD66b. Scale bars, 25 µm (main) and 5 µm (insets). (h) Neutrophil functional-module scores across subpopulations. (i) TAN signature scores in metastatic prostate cancer samples from the Stand Up To Cancer (SU2C) cohort, stratified by survival status. (j) Kaplan-Meier analysis of overall survival in SU2C samples stratified by the median TAN score. (k) Correlations between neutrophil ROS- or nitric-oxide-related programs and DNA-damage or nitrogen-response programs in CD8 T cells across the main cohort. Each point represents one patient sample. (l) UMAP of CD8 T cell subtypes and density maps by disease group. (m) Expression of dysfunction-related genes in CD8Temra cells across integrated healthy, Benign, HMP and Mets datasets; dot size indicates the expressing-cell fraction and color indicates scaled mean expression. (n) NicheNet-predicted ligands and target genes associated with CD8Temra alteration in HMP. (o) Relative *SLAMF6* and *SLAMF7* mRNA abundance in healthy male peripheral blood CD8 T cells after IL-1β stimulation. Data are presented as mean ± s.e.m.; n = 3 donors. (p, q) Diffusion-map and RNA-velocity analysis of CD8Temra-state transitions, with cell-density maps by disease group. (r) Fitted exhaustion and hypoxia scores along the inferred trajectories. (s) Regulon activity across the CD8Temra-to- CD8Trm and CD8Temra-to-CD8Thx trajectories. (t) CD8Thx and CD8Trm ssGSEA scores in SU2C samples stratified by survival status. (u) Correlation between the CD8Thx ssGSEA score and IL1B expression in SU2C samples. (v) Correlation between the proportion of CD8Thx cells and neutrophil *IL1B* expression in the main cohort. (w) Proposed model linking TANs with CD8 T cell dysfunction across HMP and metastatic BM. *Unless otherwise indicated, bar plots with error bars show mean ± s.d. Statistical significance was assessed using two-sided unpaired Student’s t-tests (o), Wilcoxon rank-sum tests (d, i, t), a log-rank test (j), Pearson correlation (u) and Spearman correlation (k, v).*

Unbiased, graph-based clustering of **174,662** high-quality cells identified major hematopoietic and stromal lineages, as well as metastatic prostate epithelial cells (Figures 1b, S1a). Among the stromal populations, osteoblasts were prominently expanded in the Mets group (Figures 1b-d). We further annotated these cells into several subtypes (Figure S1b-d). By comparing them with published normal BM osteoblasts (Figure S1e-i), we found that the immature osteoblast subset OB1 displayed an *ALPL*^hi^ *BGLAP*^low^ woven-bone signature, a feature commonly associated with osteoblastic PCa bone metastases ^21–24^. OB1 also showed increased expression of cancer-associated fibroblast markers (Figure S1h) ^25–28^. Cell-cell communication analysis further suggested a potential association between tumor-derived TGFB3 signaling and OB1 expansion ^29^ (Figure S1j-n).

Endothelial cells and macrophages were also increased in metastatic BM. After integrating our endothelial cells with published prostate and normal BM endothelial cell datasets (Figure S2a-f), we found that endothelial cells from bone metastases more closely resembled prostate endothelial cells than normal bone marrow (BM) endothelial cells ^30^ (Figure S2f). This similarity was particularly evident in the *ESM1*^+^ CapEC2 subset, which showed features of tip endothelial cells (Figure S2e, f) ^31–35^. Consistently, cell-cell communication analysis predicted VEGFA signaling from metastatic PCa cells to the pro-angiogenic CapEC2 population (Figure S2h-j). In parallel, we identified a *SIGLEC8*^+^ macrophage subset, termed Macro1, that showed high similarity to prostate-associated macrophages (Figures S2k-q) ^36^.

Together, these findings indicate that established bone metastases are accompanied by broad tumor-associated remodeling of the stromal, vascular, and myeloid compartments. However, stage-wise comparison suggested that these architectural changes were mainly enriched in the Mets stage and therefore may represent relatively late events during metastatic niche formation. By contrast, immune alterations were already evident at the HMP stage, before detectable tumor-cell colonization of the BM (Figures 1b-d, S1a). Notably, a distinct neutrophil subset, annotated as TANs, was already expanded in the HMP group and remained enriched in the Mets group (Figures 1c-d). Because our study aimed to identify early microenvironmental changes associated with bone metastatic potential, we next focused on the origin and function of this early-emerging TAN population.

### A unique neutrophil subtype emerges in HMP BM and exhibits multifaceted immunosuppressive capabilities

Besides the established metastatic niche, HMP BM already exhibited significant microenvironmental reprogramming compared with benign BM. Most notably, a tumor associated neutrophil (TAN) subtype was significantly enriched in BM from HMP and Mets groups (Figures 1b-d, S3a). Five distinct neutrophil populations were identified: progenitors (NeuP), pre-neutrophils (preNeu), immature neutrophils (immNeu), mature neutrophils (mNeu), alongside TANs, defined by high expression of *CCR3* and *ICAM1* (Figures 1e, f). Integrated analysis using a public BM dataset did not identify a comparable TAN state in normal BM ^37–39^ (Figures S3b-e). Despite inter-patient variation in abundance, the TAN state and its associated transcriptional features were reproducibly detected across multiple independent HMP and Mets samples (Figures S3f-k). Additionally, TAN enrichment was also observed irrespective of androgen-deprivation therapy (ADT) status (Figures S4a-d). TANs were detected in both HMP and Mets BM, with limited transcriptomic differences (Figures S4e-i). Immunofluorescence staining further corroborated the presence of CCR3^+^ICAM1^+^CD66b^+^ TANs within metastatic bone lesions (Figures 1g, S4j), whereas flow cytometry did not identify a comparable TAN phenotype in peripheral blood (PB) (Figures S4k, l).

Compared with TANs associated with primary PCa, BM-resident TANs lacked expression of *F10* and *TREM2*, representing a previously unreported subtype ^40,41^ (Figure S4m). Transcriptomically, TANs displayed an immunosuppressive-like profile: they showed reduced expression of genes associated with antigen presentation pathways while upregulating signatures of myeloid-derived suppressor cells (MDSC) (Figure 1h). TANs also displayed elevated levels of immune checkpoint molecules, including *CD274* (PD-L1) and members of the butyrophilin (BTN) family (Figures 1f, S4m) ^42,43^, alongside upregulation of immunosuppressive factors *PTGS2* and *IL1B* ^44^. This was further accompanied by enrichment of gene signatures associated with nitric oxide (NO) biogenesis ^45^ and reactive oxygen species (hydrogen peroxide) metabolic pathways (Figures 1h, S5a, b). Specifically, TANs exhibited elevated expression of genes encoding hydrogen peroxide production enzymes and the exporting channel *AQP9*, alongside decreased expression of ROS clearance-related genes (Figures S5b, c), which raised the possibility that TANs might contribute to a microenvironment conducive to T cell hyporesponsiveness and DNA damage ^46,47^. Consistently, TAN-derived ROS production and NO biosynthesis pathways exhibited a strong correlation with the CD8 T cell DNA damage response and nitrogen response scores, respectively (Figures 1k and S5d-g). Furthermore, aligning with potential roles in cellular survival and tumor interaction, TANs also showed high expression of longevity-associated genes (*MCL-1*, *CDKN1A*) and the pro-tumor factor *OSM* ^48,49^ (Figure S5h). As expected, survival analysis in the Stand Up To Cancer (SU2C) mPCa bone metastasis samples revealed that higher TAN signature score was significantly associated with worse overall survival in mPCa patients ^29^ (Figures 1i, j, S5i).

### A dysfunctional T cell and monocyte landscape correlates with TAN enrichment

Given their immunosuppressive transcriptomic signature, we next investigated the potential correlation between TANs and the altered immune landscape in the BM niche from the HMP group. Coinciding with the emergence of TANs, we observed a significant expansion of terminally differentiated effector memory CD8 T cells (CD8Temra) (Figures 1l, S5j, k). Considering the low abundance of CD8Temra in our benign stage, we integrated CD8 T cells in healthy and benign BM from published datasets for validation and confirmed this observation ^15,50^ (Figures S5l-o). Compared with CD8Temra in healthy BM, CD8Temra in HMP were dysfunctional, exhibiting molecular signatures of pre-exhaustion, including upregulation of *SLAMF6* ^51^, *SLAMF7* ^52^, and the transcription factors (TFs) *IKZF1* and *IKZF3* ^53,54^ (Figures 1m, S5p, q). NicheNet analysis predicted IL-1β as a candidate ligand associated with this pre-exhausted state (Figure 1n). Consistently, IL-1β stimulated human PB CD8 T cells upregulated expression of *SLAMF6* and *SLAMF7* (Figure 1o). Notably, TANs constituted a major source of *IL1B* expression in HMP (Figure S5r, s), suggesting they are potential contributors to the observed CD8Temra pre-exhaustion. Moreover, *CCL5* was highly expressed in T cells (Figure S5t), potentially mediating chemotaxis of the *CCR3* expressing TANs toward T cells via CCL5-CCR3 signaling, followed by ICAM1-ITGAL mediated adhesion between these two cell populations. This proximity could further facilitate the suppression of T cells by TANs ^55^.

Concurrently, we observed disrupted monocyte differentiation in the HMP group, which coincided with the expansion of TANs. After reclustering, we annotated monocytes as classical monocytes (cMono1, cMono2), non-classical monocytes (ncMono) and intermediate tumor associated macrophages (intTAM) ^56,57^ (Figure S6a-f). Trajectory analysis suggested cMono2 represents a transitional state differentiating into ncMono (Figure S6e) ^58^. While the HMP bone niche exhibited an accumulation of cMono2 (Figure S6f), their subsequent differentiation into protective, anti-metastasis ncMono was altered ^59^ (Figure S6f). This pattern was further validated using integrated healthy and benign BM data from published dataset (Figures S6g-j). After comparing the expression of ncMono differentiation related genes between healthy and HMP in cMono2, we noticed downregulated *NR4A1*, a master regulator of ncMono differentiation ^60,61^, in cMono2 from HMP (Figure S6k). Cell-cell communication analysis suggested a potential interaction with TANs: the upstream inhibitor of NR4A1 signaling, *HAVCR2* (*TIM-3*) ^62^ (Figure S6l), was globally expressed on these monocytes (Figure S6m), while its ligand, *LGALS9* (Galectin-9), was predominantly upregulated in TANs in the HMP bone niche (Figure S6n, o). Consistent with putative HAVCR2 activation and NR4A1 suppression, downstream target genes were significantly altered in cMono2 cells from both HMP and Mets groups ^63,64^, providing transcriptomic support for a hypothesized TAN-associated LGALS9-HAVCR2-NR4A1 axis (Figure S6p). Moreover, among the cell populations examined, TAN abundance showed the strongest positive correlation with cMono2 abundance (Figure S6q). Aside from monocytes and CD8 T cells, minimal alterations were observed in other immune compartments in the HMP group, including CD4 T cells, Natural killer (NK) cells and B cells (Figures S7a-k). Taken together, our transcriptomic data suggest that the accumulation of TANs closely correlates with an immunosuppressive BM environment in HMP group, potentially linked to the observed dysregulation of CD8Temra and the stalled differentiation of anti-metastatic ncMono.

### TAN-associated IL-1β correlates with a profoundly immunosuppressive T cell landscape in established metastases

We next sought to explore whether the immunosuppressive role of TAN persists into Mets stage. Analysis of CD8 T cells in Mets suggested that our previously identified dysfunctional HMP CD8Temra cells may represent a transitional state towards the terminally exhausted resident memory (CD8Trm) and hypoxic T cells (CD8Thx) populations, both of which are known to be dysregulated in tumor progression ^65–67^ (Figures 1p-r, S7l-n). This transcriptional shift was associated with a unique set of TFs (Figure 1s), including regulators implicated in CD8 T cell dysfunction ^68–71^. *PDCD1* emerged as the commonly and most highly expressed inhibitory immune checkpoint within these cells (Figure S7o). After comparing their signature in SU2C mPCa bone-metastasis samples, we revealed that CD8Thx was particularly associated with poor prognosis (Figure 1t). Notably, computational analysis predicted IL1B as a candidate ligand associated with the dysregulated signature in CD8Thx (Figure S7p). Correspondingly, TANs showed relatively high mean *IL1B* expression and constituted a substantial fraction of *IL1B*-expressing cells in metastatic BM (Figure S5r, s). Consistently, we also observed significant positive correlation between *IL1B* and CD8Thx in bulk and scRNA-seq data (Figures 1u, v).

On the other hand, markedly expanded in metastatic BM (Figure S7k, q-s). Among the ligands that interacted with CD4Treg, *IL1B* showed the most prominent upregulation (Figure S7t). IL-1β was reported to be associated with Treg expansion ^72,73^. Consistently, in SU2C mPCa bone-metastasis samples, *IL1B* expression was significantly correlated with *FOXP3* (Figure S7u). Moreover, *IL1B* expression in monocytes, macrophages and TAN was significantly correlated with Treg abundance (Figure S7v). Together, these data suggest an association between IL-1β-expressing TANs and an immunosuppressive BM microenvironment in Mets, potentially linking to the observed emergence of CD8Thx and the expansion of CD4Tregs (Figure 1w).

### Distal signals from primary tumor induce BM-specific TANs

Having observed the pre-metastatic enrichment of TANs, we sought to identify the potential upstream signals associated with TANs. Initial analysis of the HMP BM using NicheNet failed to identify a locally-produced ligand that was significantly upregulated. Moreover, the absence of TAN in circulation argues against their migration from peripheral tissues into BM (Figures S4k, l). Therefore, we hypothesized that a distal signal originating from the primary tumor might contribute to this phenomenon. To explore this hypothesis in vivo, we utilized a surrogate model via intraperitoneal (i.p.) injection of conditioned medium (CM) from TRAMP-C1 cells or matched cell-free serum-free medium processed in parallel as the control (Figure 2a). TRAMP-C1 CM induced several BM alterations observed in the HMP cohort, including the expansion of CCR3^+^ neutrophils, CD8Temra cells, and intermediate monocytes, alongside a decrease in non-classical monocytes (Figures 2b, c, S8a-c). The absence of these phenotypes in the PB and spleen suggested the relative specificity of this alteration in BM (Figures S8d-f). Histological analysis showed spatial proximity between CCR3^+^ neutrophils and CCL5^+^ ITGAL^+^ T cells (Figure S8g), consistent with the cellular interaction pattern described above.

**Fig. 2.**
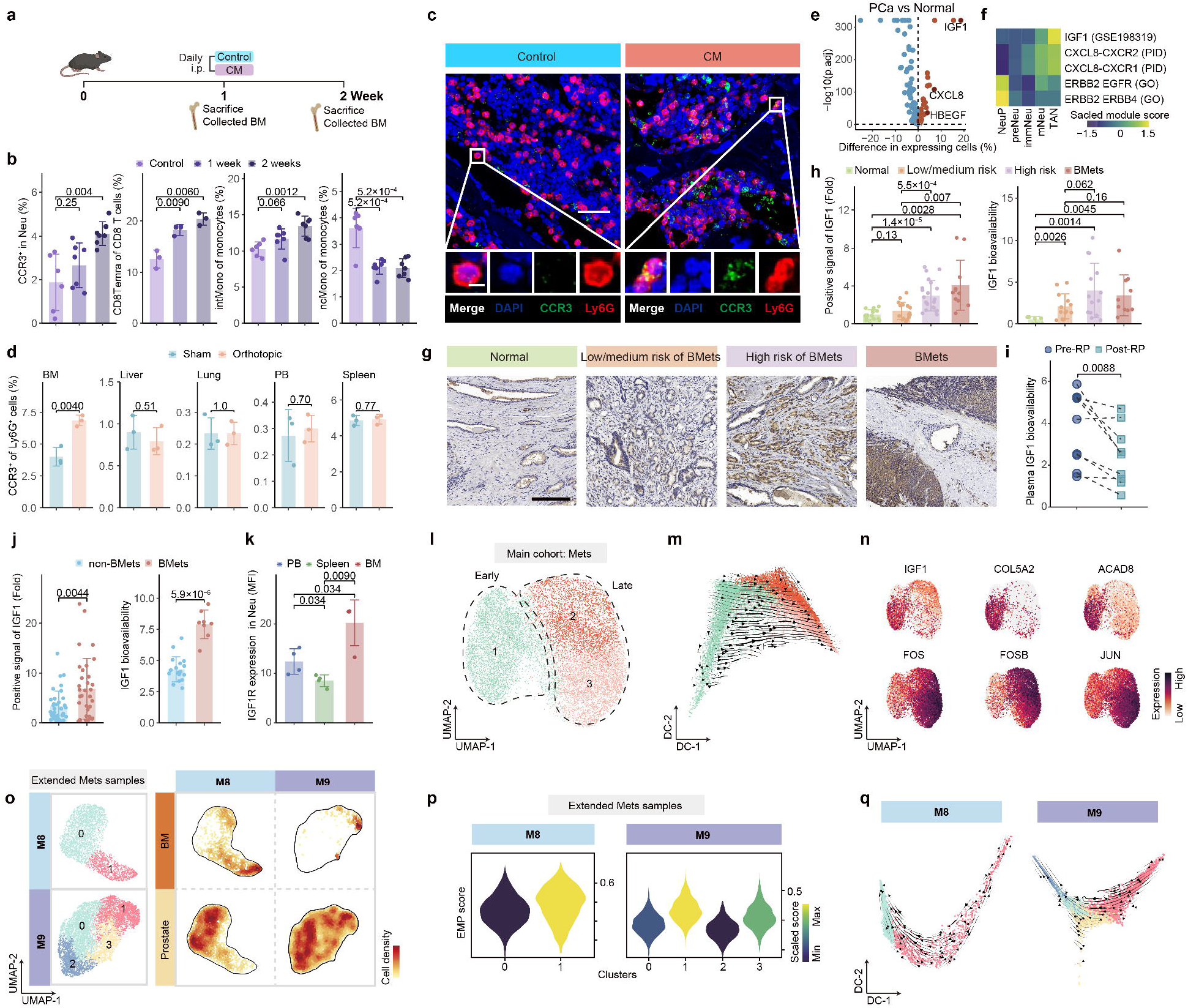
Tumour-associated systemic IGF1 is linked to TAN emergence in the bone marrow. (a) Experimental design of the conditioned-medium (CM) model. Mice received daily intraperitoneal injections of TRAMP-C1 CM or matched blank medium (Control) for 2 weeks; BM was collected after 1 or 2 weeks. (b) Flow-cytometric quantification of CCR3-positive neutrophils, CD8Temra cells, intermediate monocytes (intMono) and non-classical monocytes (ncMono) during niche establishment. Each dot represents one mouse; for CCR3 neutrophils, intMono and ncMono measurements, n = 6, 7 and 7 mice for Control, 1 week and 2 weeks groups, respectively. For CD8 T cells, n = 3 mice per group. (c) Representative multiplex immunofluorescence images of BM stained for DAPI, CCR3 and Ly6G. Scale bars, 50 µm in the main images and 5 µm in the insets. (d) Frequency of CCR3-positive Ly6G-positive neutrophils in BM, liver, lung, peripheral blood (PB) and spleen from sham-operated and orthotopic prostate-tumor-bearing mice. Each dot represents one mouse; n = 3 mice per group. (e) Differential expression in epithelial cells from primary prostate cancer versus normal prostate datasets; selected tumor-secreted factors are labeled. (f) Activity of IGF1-, CXCL8-and ERBB-family downstream gene modules across neutrophil states. (g) Representative immunohistochemical staining of IGF1 in normal prostate and prostate cancer tissues from patients with low/medium risk, high risk or established bone metastasis. Scale bar, 200 µm. (h) Quantification of prostate-tissue IGF1 staining and plasma IGF1 bioavailability across normal, low/medium-risk, high-risk and bone-metastatic prostate cancer groups. For tissue IGF1 staining, normal prostate, n = 15 patients; low/medium-risk PCa, n = 16; high-risk PCa, n = 21; bone-metastatic PCa, n = 11. For plasma IGF1 bioavailability, normal prostate, n = 6 patients; low/medium-risk PCa, n = 13; high-risk PCa, n = 14; bone-metastatic PCa, n = 10. (i) Paired plasma IGF1 bioavailability before and after radical prostatectomy (RP); n = 9 matched patient pairs. (j) IGF1 staining in primary prostate cancer tissues and plasma IGF1 bioavailability in patients with advanced localized PCa, stratified according to subsequent bone metastasis development during follow-up (non-BMets, remained free of bone metastasis; BMets, subsequently developed bone metastases). For tissue IGF1 staining, non-BMets, n = 36 patients; BMets, n = 33. For plasma IGF1 bioavailability, non-BMets, n = 18 patients; BMets, n = 8. (k) Flow-cytometric quantification of IGF1R mean fluorescence intensity (MFI) on neutrophils from PB, spleen and BM. Each dot represents one mouse; n = 4, 4 and 3 mice for PB, spleen and BM, respectively. (l) UMAP of epithelial cells in metastatic BM samples. Early and late subtypes were labeled. (m) Diffusion-map and RNA-velocity analysis of epithelial-subtype progression. (n) UMAP feature plots of *IGF1*, *COL5A2*, *ACAD8*, *FOS*, *FOSB* and *JUN*. (o) Malignant epithelial cells from paired primary-prostate cancer and BM samples from patients M8 and M9, shown by cluster identity and tissue-specific density. (p) EMP scores across paired-sample epithelial clusters. (q) Patient-specific diffusion maps with RNA-velocity vectors, colored by cluster identity. *Unless otherwise indicated, bar plots with error bars show mean ± s.d. Statistical significance was assessed using two-sided unpaired Student’s t-tests (b,d,h,j,k), a paired Student’s t-test (i), Fisher’s exact tests with multiple-testing correction (e). Student’s t-test P values were adjusted using the Benjamini-Hochberg method where multiple comparisons were performed.*

To further corroborate these findings, we established an orthotopic PCa mouse model (Figure S8h). Orthotopic tumor growth increased IGF1 abundance and bioavailability in both PB and BM (Figure S8i). Notably, CCR3^+^Ly6G^+^ neutrophils increased in the BM, whereas no comparable increase was observed in peripheral blood, spleen, liver or lung (Figure 2d and Figure S8j). Moreover, after analyzing mouse samples from previous study by Guo et al ^74^, we didn’t observe appreciable accumulation of CCR3^+^Ly6G^+^ within spontaneous PCa liver and lung metastatic lesions (Figures S8k), supporting preferential enrichment of this TAN population in the BM among the tissues examined.

### Primary tumor-associated IGF1 correlates with the formation of TANs

Next, to investigate potential signals associated with PMN-like alterations, we collected prostate samples from three patients of HMP group and performed scRNA-seq, which resulted in three paired prostate-BM data (Extended HMP samples). We applied an adapted ligand-receptor prioritization workflow across these paired datasets to nominate secreted factors that could potentially act on bone marrow neutrophils (Figures S9a-c) (Methods). Based on this computational prediction, we nominated three candidate ligands, among which insulin-like growth factor 1 (*IGF1*) exhibited the most significant upregulation in PCa compared to normal prostate (Figure 2e). We independently analyzed candidate ligands associated with TAN differentiation within the metastatic BM microenvironment, where IGF1 again ranked among the leading predicted ligands (Figure S9d). Moreover, *IGF1* was expressed predominantly by epithelial cells (Figure S9e). Analysis of pan-cancer clinical samples revealed that *IGF1* was mainly expressed by fibroblasts ^75^. Surprisingly, the upregulation of *IGF1* in epithelial cells was a specific phenomenon in PCa (Figure S9f). Moreover, inferred IGF1 pathway activity was highest in TANs, among the major BM cell populations examined (Figures 2f and S9g).

IGF1 is a critical growth factor, which is primarily synthesized and secreted by the liver and reported to function in an endocrine manner ^76^. Several studies have reported its correlations with bone metastasis ^77,78^. However, the specific role of IGF1 in PCa bone metastasis remains unclear. In the BM microenvironment, *IGF1* expression is largely restricted to osteoprogenitors and megakaryocytes ^79^, while other cell types showed relatively low IGF1 expression ^39^. Accordingly, we did not observe significant IGF1^+^ cells in the BM of HMP patients (Figure S9h). In stark contrast, within established bone metastases, *IGF1* was highly expressed by the malignant epithelial cells themselves and by an abnormally expanded osteoblast population (OB1) absent from the normal BM (Figure S9h).

To further investigate the relationship between the primary tumor and systemic IGF1 bioavailability, we performed a series of validation experiments. We first confirmed that IGF1 expression was progressively upregulated in primary prostate tumors and that its bioavailability increased in circulation with disease progression towards bone metastasis in clinical samples ^80^ (Figures 2g, h). Critically, plasma IGF1 bioavailability significantly decreased following surgical removal of the primary tumor, consistent with a contribution from the primary tumor or tumor-associated systemic changes to circulating IGF1 bioavailability (Figure 2i). To evaluate the association between IGF1 and subsequent bone metastasis, we analyzed pre-operative samples from advanced localized PCa patients and monitored them for subsequent bone metastasis development. This analysis revealed that patients who later developed bone metastases had significantly higher IGF1 bioavailability in both their primary tumor tissue and plasma at the time of diagnosis compared to those who remained metastasis-free (Figures 2j, S9i). Meanwhile, Gleason scores and T stages between these two groups exhibited no significant differences (Figure S9j), supporting an association between elevated IGF1 bioavailability and subsequent bone metastasis. Together, these clinical correlations suggest that elevated, primary tumor-related IGF1 is associated with early BM remodeling and bone metastatic progression.

Next, we validated the expression of *IGF1R* in neutrophils. Transcriptomic profiling revealed predominantly higher *IGF1R* expression in BM neutrophils relative to other cell types in human samples, highlighting neutrophils as key candidates for receiving IGF1 signaling in the BM niche (Figures S9k-n). Furthermore, comparing IGF1R expression in neutrophils across the BM, PB and spleen revealed significantly higher IGF1R expression in BM neutrophils at both transcript and protein levels (Figures 2k, S9o).

### An IGF1-expressing epithelial program is associated with early metastatic states

To address whether IGF1 production is a general feature of tumor cells, we analyzed the metastatic malignant epithelial cells within our main dataset. As expected, luminal epithelial cells exhibited high inter-individual heterogeneity (Figures S10a-d) ^81,82^. We thus clustered the malignant epithelial cells based on their transcriptional regulon activity ^82^ (Figures 2l-n, S11a-d). Three distinct subclusters (C1, C2, and C3) were identified and the trajectory inference suggested that these clusters represent a continuum from C1 towards C2/C3 (Figure 2m).

The C1 subcluster, which highly expressed *IGF1* (Figures 2n, S11b), was positioned at the beginning of the trajectory and exhibited high regulation activity of TFs associated with stemness, such as members of the HOX family, NKX3-1, and MYBL2 ^83–85^ (Figure S11c). Conversely, the C2 and C3 subclusters represented later stages, which were characterized by increased activity of regulons implicated in cancer progression, including GRHL2, SRF, EGR1, and GATA2 ^86–89^. Further DEGs enrichment and NMF-based meta-programs analysis revealed enrichment for signatures of cellular stress, hypoxia, and AP-1 activation in these late-stage clusters ^90–93^ (Methods) (Figures S11e-g) (Table S3).

Furthermore, C2 and C3 exhibited stronger cellular communication with other BM components than C1, consistent with their identity as later-stage populations that are more integrated into the BM microenvironment (Figure S11h). Thus, we termed C1 as early metastatic PCa cells (EMPs) with IGF1 as a defined marker and C2/C3 as late metastatic PCa cells (LMPs) (Figure 2n). Metabolic analysis demonstrated EMPs possess early PCa-specific metabolism signature, including fatty acid and keratan sulfate synthesis ^94–96^ (Figure S11i). In the SU2C mPCa bone metastasis samples, EMP-dominant samples were associated with longer overall survival than LMP-dominant samples, consistent with EMP representing a relatively earlier state within established metastatic disease (Figures S11j, k).

On the other hand, when we evaluated the EMP and LMP signature in primary tumors, we found that primary tumors from patients with distant metastasis showed a higher EMP score, whereas no significant difference was observed for the LMP score (Figures S11l-p) ^30,97^. For further validation, we performed paired sequencing of primary tumor and BM samples from another two mCRPC patients (Extended Mets samples) (Figure S11q-v). Across the two Extended Mets samples, malignant epithelial cells from BM showed higher EMP scores than those from the matched primary-prostate samples, suggesting a potential enrichment of EMP-like cells during bone metastatic colonization (Figure S11w). Strikingly, we identified a common C1 cluster with a high EMP score in both patients analyzed (Figures 2o, p, S11x, y). Moreover, RNA velocity analysis computationally supported the inference of these C1 cells as a potential origin state within the metastatic continuum (Figure 2q), highlighting the potential relevance of IGF1-expressing EMP cells in early bone metastasis. Together, these findings identify IGF1-expressing EMP-like cells as a recurrent early metastatic state and a plausible source of tumor-derived IGF1 during bone colonization. This prompted us to next determine whether IGF1 could directly reprogram BM neutrophils toward the TAN phenotype observed in HMP and metastatic BM.

### IGF1 regulates neutrophils towards a pro-tumorigenic state via an MAPK/ERK-KLF4 axis

To directly evaluate the role of IGF1 in regulating neutrophil phenotypes, we treated primary mouse BM neutrophils with recombinant IGF1 (Methods). This treatment promoted the acquisition of a TAN-like phenotype, characterized by the upregulation of TAN-specific markers, the enrichment of TAN derived signatures, and increased hydrogen peroxide production (Figures 3a-d, S12a). In contrast, neutrophils isolated from PB were not responsive to IGF1 under the same conditions (Figure 3c-d). Consequently, we referred to these BM-derived, IGF1 activated cells as IGF1-induced TANs (iTANs).

**Fig. 3.**
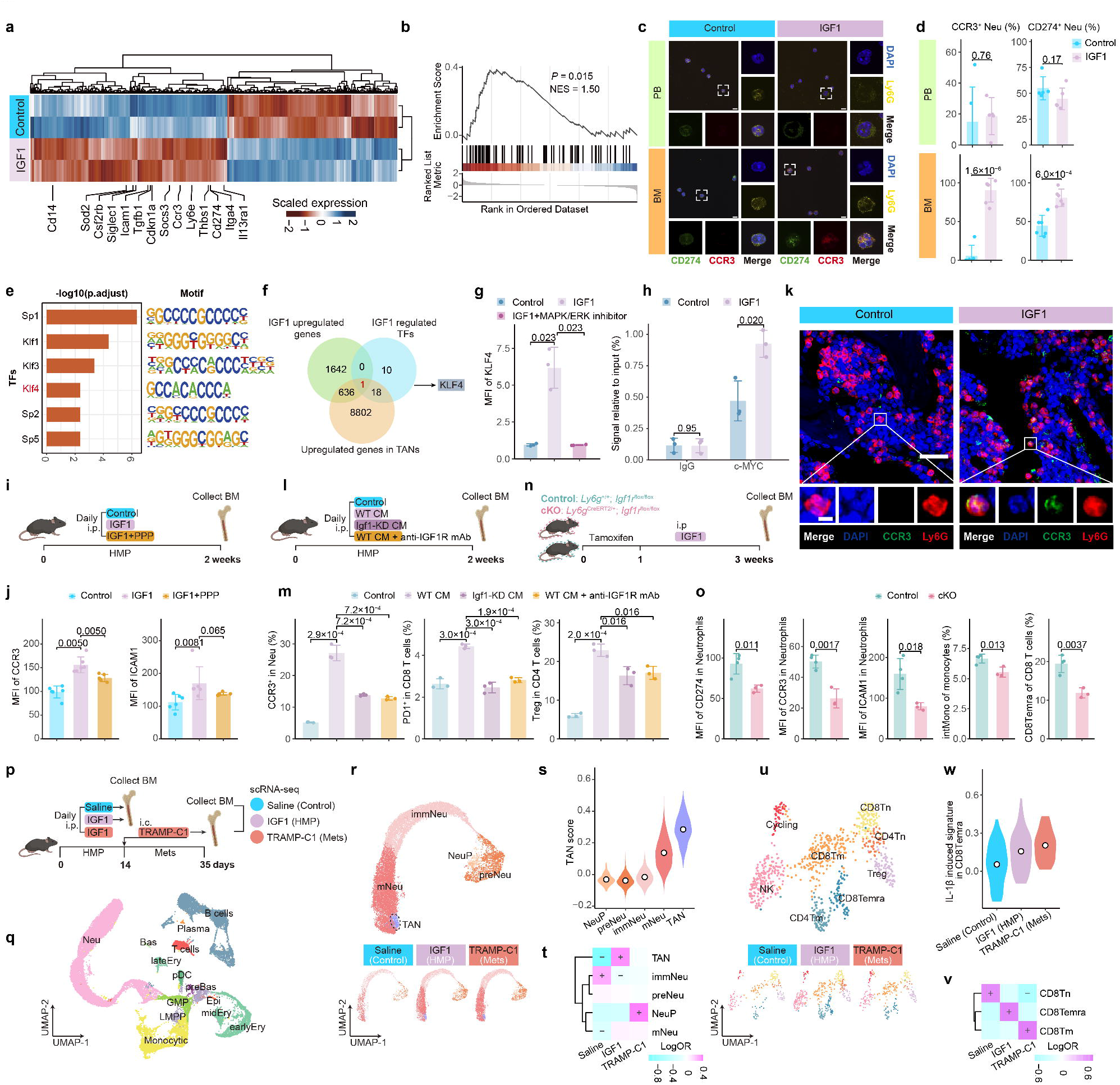
IGF1-IGF1R signaling drives MAPK/ERK-c-MYC-KLF4-dependent TAN-like neutrophil reprogramming and immunosuppressive remodeling. (a) Heatmap of genes differentially expressed in control and IGF1-treated mouse BM neutrophils; values are scaled expressions. Two mice were analysed per group, with each mouse serving as an independent biological replicate. (b) Gene-set enrichment analysis of the TAN signature in IGF1-treated versus control neutrophils using the RNA-seq data shown in a (n = 2 mice per group). (c) Representative immunofluorescence images of peripheral-blood and BM neutrophils from control- or IGF1-treated mice stained for DAPI, CD274, Ly6G and CCR3. Scale bars, 10 µm. (d) Frequencies of CCR3-positive and CD274-positive neutrophils in peripheral blood and BM; n = 3 mice per group. (e) Transcription-factor motifs enriched in accessible chromatin after IGF1 stimulation. n = 2 biologically independent neutrophil preparations per group. (f) Intersection of IGF1-upregulated genes (RNA-seq), IGF1-regulated transcription factors (ATAC-seq) and TAN-upregulated genes (scRNA-seq), identifying KLF4 as a candidate regulator. (g) KLF4 MFI in neutrophils after IGF1 stimulation with or without MAPK/ERK inhibition; n = 3 biologically independent samples per group. (h) ChIP-qPCR analysis of c-Myc occupancy at the KLF4 promoter after IGF1 stimulation; normal IgG served as the negative control. n = 3 biologically independent samples per group. (i) Experimental design for IGF1-induced BM preconditioning in mice receiving matched vehicle (Control), IGF1 or IGF1 plus the IGF1R inhibitor picropodophyllin (PPP). (j) CCR3 and ICAM1 MFIs in BM neutrophils after the indicated treatments; n = 6 mice per group. (k) Representative BM immunofluorescence images stained for DAPI, CCR3 and Ly6G. Scale bars, 50 µm in the main images and 5 µm in the insets. (l) Experimental design of the CM-induced BM-preconditioning model using matched blank medium (Control), wild-type TRAMP-C1 conditioned-medium (WT CM), Igf1-knockdown CM (Igf1-KD CM) or wild-type CM combined with an IGF1R-blocking antibody. (m) Frequencies of CCR3-positive neutrophils, PD-1-positive CD8 T cells and Tregs after the indicated CM treatments; n = 3 mice per group. (n) Experimental design for analysis of IGF1-induced BM remodeling in neutrophil-specific *Igf1r* conditional-knockout (cKO) mice and littermate controls. (o) CD274, CCR3 and ICAM1 MFIs in neutrophils, intermediate-monocyte frequency and CD8Temra frequency in control and cKO mice after IGF1 treatment; n = 3 mice per group. (p) Design of the mouse BM scRNA-seq experiment comprising saline-control, IGF1-induced HMP-like, and IGF1-preconditioned TRAMP-C1 metastatic groups. For each experimental group, BM cells from three mice were pooled to generate one scRNA-seq sample. (q) UMAP of all mouse BM cells, colored by major cell type. (r) UMAP of mouse neutrophil states with group-specific embeddings. (s) Human-derived TAN signature scores across mouse neutrophil states. (t) Differential abundance of mouse neutrophil states across experimental groups. Color indicates the log odds ratio, and “+” or “-” denotes significant enrichment or depletion, respectively. (u) UMAP of mouse BM CD8 T cell subtypes with group-specific embeddings. (v) Differential abundance of CD8 T cell subtypes across experimental groups. (w) IL-1β-induced signature scores in CD8Temra cells across saline-control, IGF1-induced HMP and TRAMP-C1 metastatic groups. *Unless otherwise indicated, bar plots with error bars show mean ± s.d. Statistical comparisons in panels d, g, h, j, m and o were performed using two-sided Student’s t-tests. P values were adjusted using the Benjamini–Hochberg method where multiple comparisons were performed. For the GSEA in panel b, nominal P values and normalized enrichment scores are shown.*

Functionally, iTANs recapitulated key features of native TANs, suppressing CD8 T cell cytotoxicity and proliferation partly through IL-1β-, ROS- and NO-dependent mechanisms, promoting Treg proliferation, and inducing a hypoxia-associated CD8 T cell state (Figs. S12b-d). iTANs also enhanced PCa-cell migration and invasion, which was attenuated by IGF1R suppression (Figs. S12e).

To dissect the underlying mechanism, we performed ATAC-seq and RNA-seq analyses of iTANs and identified the TF KLF4 as a key downstream effector (Figures 3e, f). Inhibition of KLF4 markedly attenuated IGF1-induced TAN-like phenotype (Figure S12f). To delineate the upstream signaling pathway connecting the IGF1R to KLF4, we first evaluated the roles of the MAPK/ERK and AKT pathways, both known to be downstream of IGF1R signaling ^98^. Our results demonstrated that IGF1-induced KLF4 upregulation is dependent on the MAPK/ERK pathway (Figures 3g, S12g).

We next sought to identify the intermediary TF linking MAPK/ERK activation to *Klf4* transcription. After evaluating the Klf4 promoter binding potential of TFs acting downstream of MAPK/ERK ^99^ (Table S4), filtering for candidates upregulated upon IGF1 stimulation in our RNA-seq data, we prioritized c-Myc as the top candidate (Table S4). Its potential intermediary role was supported by Chromatin immunoprecipitation (ChIP)-qPCR and flow cytometry (Figures 3h, S12h). Pharmacological inhibition of either MAPK/ERK or c-Myc phenocopied KLF4 inhibition, preventing iTAN formation (Figure S12i). Together, these in vitro findings support a model where the IGF1R-MAPK/ERK-c-MYC-KLF4 signaling axis mechanistically contributes to the phenotypic shift of naive neutrophils towards a TAN-like state.

### Systemic IGF1 exposure induces HMP-like alterations and enhances metastatic burden in vivo

We next examined the role of IGF1 and IGF1R signaling in TAN formation and BM remodeling in vivo. We first administered recombinant IGF1 systemically to mice. IGF1 treatment increased TAN-like neutrophils in the BM, whereas treatment with the IGF1R inhibitor PPP reduced this effect (Figures 3i-k). We then asked whether IGF1 produced by tumor cells could contribute to this neutrophil phenotype. We co-cultured mouse BM neutrophils with wild-type, *Igf1*-knockdown, or *Igf1*-overexpressing TRAMP-C1 cells. Neutrophils co-cultured with *Igf1*-overexpressing TRAMP-C1 cells showed increased TAN-associated features, whereas *Igf1* knockdown reduced the induction of *Ccr3*, *Icam1*, and *Il1b* (Figure S12j, k). We next tested these findings in vivo using the CM-induced pre-metastatic niche model. Wild-type TRAMP-C1 CM promoted TAN accumulation, increased the frequencies of CD8 T cells with exhaustion-associated features, and increased Treg abundance in the BM. These changes were reduced when *Igf1* was knocked down from tumor-cell CM or when IGF1R was blocked with an antibody (Figures 3l, m; S12l).

To further determine the neutrophil-intrinsic contribution of IGF1R signaling to IGF1-induced BM remodeling, we generated *Ly6g*^CreERT2/+^;*Igf1r*^fl/fl^ mice (cKO) to selectively disrupt IGF1R signaling in neutrophils, with *Ly6g*^+/+^;*Igf1r*^fl/fl^ littermates used as controls (Figure S12m). Following IGF1 treatment, cKO mice showed reduced CD274, CCR3 and ICAM1 expression in BM neutrophils compared with littermate controls, alongside attenuation of the IGF1-associated changes in intermediate monocytes and CD8Temra cells (Figures 3n, o). Together, these experiments support a role for IGF1 and neutrophil IGF1R-intrinsic signaling in TAN formation and the associated remodeling of the BM immune environment.

Building on this, we designed an in vivo model to approximate the stages observed in our human cohorts: a benign group treated with saline as control, a “HMP” group receiving systemic IGF1, and a “Mets” group with IGF1 education followed by intracardiac injection of TRAMP-C1 (Figure 3p). Because intracardiac tumor-cell injection results in immediate systemic dissemination, mice in the metastatic group received a defined two-week IGF1-preconditioning phase before TRAMP-C1 injection, thereby temporally separating BM remodeling from subsequent tumor-cell arrival. We then performed scRNA-seq on the BM from these mice to systematically map the cellular alterations (Figures 3q, S13a). We first confirmed the relatively high expression of *Igf1r* in neutrophils, and emergence of a TAN-like neutrophil population in the HMP and Mets groups (Figures 3r, t; S13b, c), characterized by the upregulation of canonical TAN marker genes, high TAN signature score at both the transcriptomic and regulon levels and highest expression levels of *Lgals9* (Figures 3s, S13d-f). Notably, this model reflected several key immune perturbations observed in our patient data. In the HMP mouse BM, we observed an expansion of CD8Temra cells (Figures 3u, v and S13g, h). After integration with public mouse BM datasets, we further observed an increased representation of CD8Temra cells together with increased activity of the IL-1β-induced signature ^100,101^ (Figures 3w, S13i, j). While a discrete CD8Thx population was not resolved, the murine CD8Tm cells in the Mets group displayed a upregulated CD8Thx signature (Figure S13k). CD4Tregs similarly exhibited increased immunosuppression-associated functional scores despite the absence of numerical expansion (Figure S13k). The mouse monocyte compartment reproduced several features observed in the human HMP cohort, including accumulation of cMono2-like cells in the HMP stage that showed downregulated *Nr4a1* expression and upregulated Havcr2 signaling (Figures S13l-r). Taken together, these in vivo experiments demonstrate that systemic IGF1 is sufficient to initiate the HMP-like alterations and that our mouse model reproduces the key features of immune landscape remodeling seen during human PCa bone metastasis progression.

We next sought to determine its functional consequence on metastasis. To this end, we pre-conditioned mice with either CM from TRAMP-C1 or recombinant IGF1, followed by intracardiac injection of luciferase-tagged TRAMP-C1 to model hematogenous dissemination (Figure 4a, b).

**Fig. 4.**
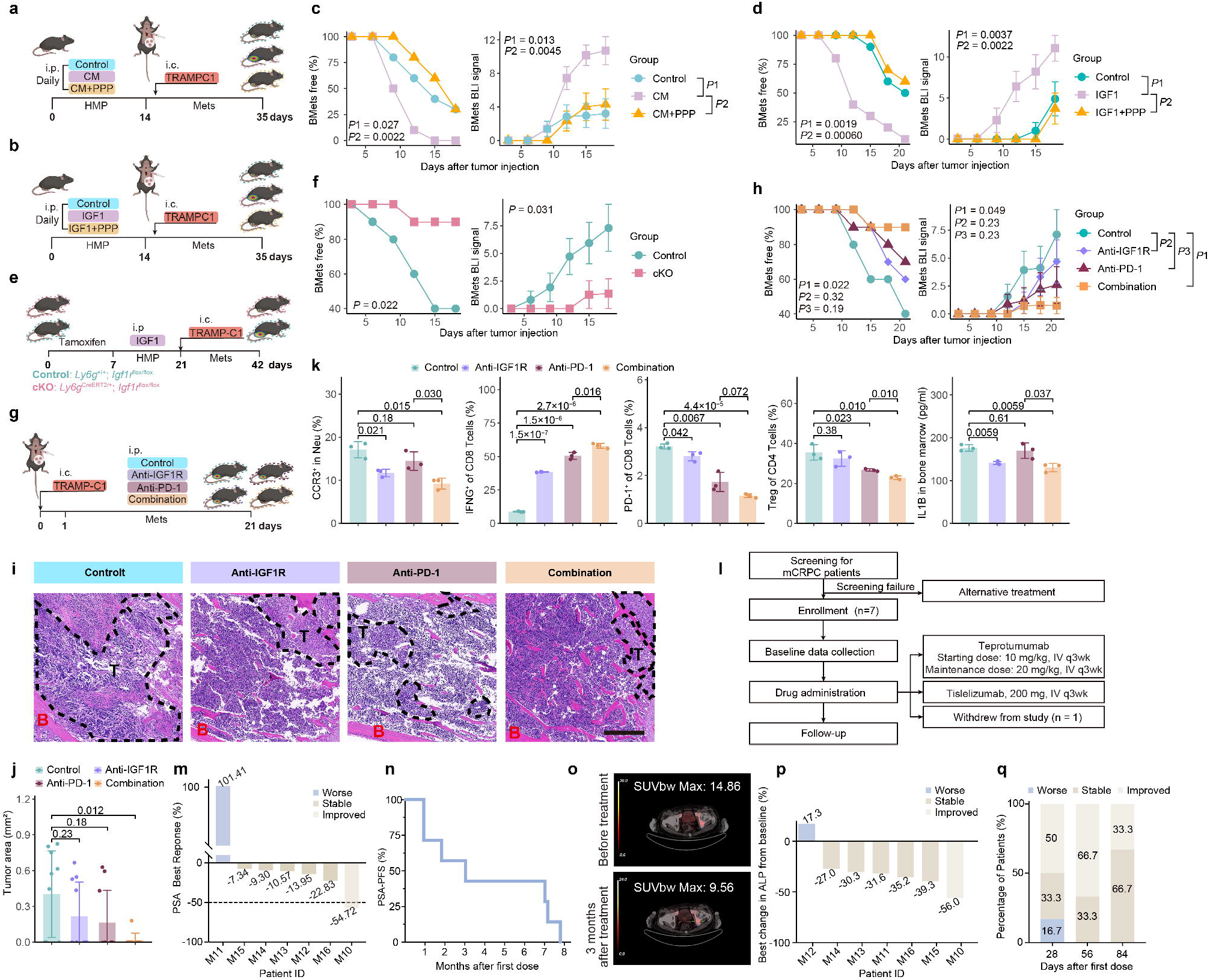
IGF1R targeting suppresses bone metastatic progression and supports combined IGF1R and PD-1 blockade. (a) Experimental design for testing whether conditioned-medium (CM)-induced BM preconditioning promotes bone metastasis and whether PPP prevents this effect. Mice received matched blank medium (Control), wild-type TRAMP-C1 CM or CM plus PPP before intracardiac TRAMP-C1 injection. (b) Experimental design for testing whether recombinant IGF1-induced BM preconditioning promotes bone metastasis and whether PPP prevents this effect. Control mice received matched IGF1 and PPP vehicles. (c, d) Bone-metastasis-free curves and longitudinal bone bioluminescence imaging (BLI) signals after CM (c) or IGF1 (d) preconditioning, with or without PPP. For longitudinal BLI analyses, the area under the curve (AUC) was calculated for each mouse from ln(signal + 1)-transformed serial BLI measurements, and per-mouse AUC values were compared between groups; n = 10 mice per group. *P*1 denotes Control versus CM in c and Control versus IGF1 in d; *P*2 denotes CM versus CM plus PPP in c and IGF1 versus IGF1 plus PPP in d. (e) Experimental design using neutrophil-specific *Igf1r* conditional-knockout (cKO) mice and littermate controls to evaluate IGF1-induced BM preconditioning and subsequent bone metastasis. (f) Bone-metastasis-free curves and longitudinal bone BLI signals in control and cKO mice. BLI signals were natural-log transformed as ln(signal + 1); n = 10 mice per group. (g) Experimental design of the metastatic model evaluating Control, anti-IGF1R, anti-PD-1 or combined anti-IGF1R plus anti-PD-1 treatment. Control mice received matched saline. (h) Bone-metastasis-free curves and longitudinal bone BLI signals across treatment groups. BLI signals were natural-log transformed as ln(signal + 1); n = 10 mice per group. *P* values for the prespecified pairwise comparisons are shown in the panel: *P*1, *P*2 and *P*3 denote Control versus Combination, Saline versus anti-IGF1R and Saline versus anti-PD-1, respectively. (i) Representative haematoxylin-and-eosin-stained bone sections from control, anti-IGF1R-, anti-PD-1- and combination-treated mice. Dashed lines outline tumor regions (T); B denotes bone. Scale bar, 400 µm. (j) Quantification of tumor area in the indicated treatment groups; each dot represents one mouse. (k) Frequencies of CCR3-positive neutrophils, IFNG-positive and PD1-positive CD8 T cells, and Tregs, together with BM IL-1β concentrations, across treatment groups; n = 3 mice per group. (l) Flowchart of the investigator-initiated trial of teprotumumab plus tislelizumab in metastatic castration-resistant prostate cancer; seven patients were enrolled and received treatment; one subsequently withdrew from the study. (m) Best percentage change in serum prostate-specific antigen (PSA) from baseline. (n) PSA progression-free survival from the first dose. (o) Representative axial PET/CT images before treatment and at day 84. (p) Best percentage change in serum alkaline phosphatase (ALP) from baseline. (q) Patient-reported pain outcomes at days 28, 56 and 84 after the first dose, categorized as improved, stable or worse relative to the immediately preceding assessment (day 28 versus baseline, day 56 versus day 28 and day 84 versus day 56); n = 6 evaluable patients at each time point. *Unless otherwise indicated, bar plots with error bars show mean ± s.d. Bone-metastasis-free curves were compared using log-rank tests. Longitudinal BLI data were compared using a two-sided Welch’s t-test. Pairwise cellular or histological comparisons were analyzed using two-sided Student’s t-tests, with Benjamini-Hochberg adjustment where multiple comparisons were performed. Exact P values are shown in the panels.*

Remarkably, both CM and IGF1 preconditioning increased subsequent bone-metastatic progression, as indicated by in vivo bioluminescence imaging, and accelerated tumor growth compared to control mice (Figures 4c, d; S14a, b). This pro-metastatic effect was significantly abrogated when mice were co-treated with an IGF1R inhibitor PPP, suggesting that this phenotype is associated with IGF1R signaling. Moreover, after PPP treatment, the immunosuppressive BM phenotype, including increased FOXP3-positive CD4 T cells and reduced GZMB-positive CD8 T cells, was attenuated.

We next tested the neutrophil-intrinsic contribution of IGF1R signaling using neutrophil-specific *Igf1r* cKO mice. Following IGF1 preconditioning and subsequent TRAMP-C1 challenge, cKO mice showed delayed bone-metastatic development and reduced longitudinal bone BLI signals compared with littermate controls (Figures 4e, f). Endpoint histological analysis independently confirmed a lower metastatic tumor area in cKO mice (Figures S14e, f). Notably, CD8 T cell dysfunction was alleviated, while Treg expansion was reduced (Figures S14g, h). These findings support neutrophil-intrinsic IGF1R signaling as a functionally important contributor to TAN-associated BM remodeling and subsequent bone metastatic progression.

Together, these in vivo observations support the hypothesis that systemic IGF1 exposure can prime the BM via IGF1-IGF1R signaling in neutrophils, contributing to an immunosuppressive environment that may facilitate subsequent metastatic colonization.

### Combined IGF1R and PD-1 blockade suppresses bone metastasis in preclinical models and shows exploratory clinical activity in mCRPC

Our preclinical findings suggest that tumor-associated IGF1 may contribute to an immunosuppressive BM niche, which provides a compelling rationale for a novel therapeutic strategy. Moreover, our human data indicated that this pathway remained prominent after metastatic colonization. Given the limited efficacy of immune checkpoint inhibitors (ICIs) as monotherapy in mPCa ^102,103^, we hypothesized that targeting IGF1R in established bone metastasis could improve the response to immune checkpoint blockade. We therefore sought to evaluate the combination of IGF1 signaling inhibition and anti-PD-1/PD-L1 immune checkpoint blockade. We first evaluated this combinatorial approach in our mouse model of bone metastasis. To model therapeutic intervention after tumor dissemination rather than prevention of BM preconditioning, TRAMP-C1 cells were introduced without prior IGF1 or CM exposure (Figure 4g).

Anti-IGF1R antibody or anti-PD-1 monotherapy produced only modest anti-tumor effects, whereas combined IGF1R and PD-1 blockade more effectively suppressed bone-metastatic progression, as assessed by bone-metastasis-free status, longitudinal BLI and endpoint histological results (Figure 4h-j). This therapeutic benefit was accompanied by partial restoration of the BM immune landscape (Figure 4k). Anti-IGF1R treatment reduced CCR3^+^ neutrophil abundance and was accompanied by increased IFNG^+^ CD8 T cell features (Figure S14i and Figure 4k). The combination treatment further increased IFNG^+^ and GZMB^+^ CD8 T cells while reducing PD-1^+^ and TIM3^+^ CD8 T cells and Treg abundance (Figure 4k, S14i).

BM IL-1β concentrations were reduced following anti-IGF1R treatment and combined anti-IGF1R/anti-PD-1 treatment, whereas anti-PD-1 monotherapy did not produce a significant reduction (Figure 4k). Notably, IGF1 levels in the liver remained stable across control, BM-preconditioned and metastatic mouse groups (Figure S14j), indicating that the observed immune remodeling was not associated with broad alterations in hepatic IGF1 abundance. To further evaluate the robustness of this therapeutic strategy, we applied the same treatment regimen in a MyC-CaP-based PCa model (Figure S14k, l). Although the combination treatment showed a consistent trend toward anti-tumor activity, the effect did not reach statistical significance. However, endpoint histological analysis suggested decreased metastatic burden in the combination treatment group (Figure S14m, n). MyC-CaP cells secreted less IGF1 into CM than TRAMP-C1 cells (Figure S14o), which may contribute to the weaker treatment effect observed in this model.

Motivated by the persistence of the IGF1-TAN-associated program in established metastatic BM and the preclinical activity of combined IGF1R and PD-1 blockade, we initiated an investigator-led, single-center, single-arm, open-label clinical trial (NCT06866548) to evaluate the safety and preliminary clinical activity of combining the IGF1R inhibitor teprotumumab with the PD-1 inhibitor tislelizumab in patients with mCRPC (Figure 4l). The first patient was enrolled on March 10, 2025. A total of seven patients with progressive mCRPC and bone metastases were enrolled. The median age was 73 years. All patients were heavily pretreated, with 100% (7/7) having prior exposure to at least two androgen receptor signaling inhibitors (ARSIs); 42.9% (3/7) had received prior chemotherapy, and 71.4% (5/7) had been treated with a PARP inhibitor (Table S5). All seven enrolled patients received at least one dose of the combination therapy and were included in the safety and efficacy analysis according to the intention-to-treat principle (Figure S15a).

Safety and tolerability constituted the primary endpoint. Treatment-related adverse events (TRAEs) of any grade were observed in 6 of 7 patients (85.7%), predominantly Grade 1-2. One Grade 3 anemia event was reported (14.3%, 1/7), and one patient experienced Grade 3 elevations in both ALT and AST (14.3%, 1/7); no Grade 4 or 5 TRAEs or treatment-related deaths occurred. The most common TRAEs were fatigue (85.7%), weight loss (85.7%), asthenia (85.7%), and tinnitus (57.1%), and AST elevation (57.1%) (Table S6), consistent with the known safety profiles of teprotumumab and tislelizumab. Overall, the combination therapy showed a manageable safety profile in this population.

The combination therapy showed exploratory evidence of clinical activity in this refractory, heavily pretreated population. Specifically, 85.7% (6/7; 95% CI, 42.1-99.6%) of participants experienced a decline in serum PSA levels from baseline (Figure 4m). Longitudinal PSA tracking showed that 6 of 7 patients had a PSA decline or stabilization relative to baseline at 1 month, compared with 4 of 7 at 2 months and 3 of 7 at 3 months after treatment initiation (Figure S15b). One patient achieved a PSA50 response, with a maximal decline of 56.7% (Figure 4m). The median PSA progression-free survival (PSA-PFS) was 3.0 months (range, 1.0-7.8 months) (Figure 4n).

Regarding radiographic outcomes, the median radiographic progression-free survival (rPFS) was 3.6 months (range, 1.0-8.1 months), according to RECIST 1.1 and PCWG3 criteria (Table S7). While no objective response was observed, the disease control rate (DCR) was 42.9% (3/7) at the primary completion assessment. Radiologic assessment revealed reduced metabolic activity in a representative bone lesion after treatment. In one representative lesion, SUVmax decreased from 14.86 at baseline to 9.56 at Day 84, supporting localized metabolic improvement (Figure 4o, S15c).

Consistent with the exploratory PSA and radiographic responses, serum alkaline phosphatase (ALP), a bone-related biomarker, showed a post-treatment reduction in 6 of 7 patients (Figure 4p, S15d), with a maximum decline ranging from 27.0% to 56.0%. Previous clinical studies have linked ALP dynamics with bone disease activity and clinical outcomes in PCa bone metastasis ^104,105^. Together, these concordant changes suggest exploratory clinical activity, although interpretation is limited by the small, single-arm cohort.

Furthermore, all evaluable patients reported improved or stable pain severity at Day 84, as assessed by the Brief Pain Inventory-Short Form (BPI-SF) questionnaire (Figure 4q, S15e, Table S8). As of the data cut-off, no patient had met the criteria for protocol-defined pain progression (TTPP). Additionally, no statistically significant deterioration or improvement in health-related quality of life (QoL) was observed. At the time of this primary analysis, overall survival (OS) data remain immature; the median OS has not been reached, and patients continue to be followed for survival.

Prior to treatment initiation, we collected BM samples from three of the patients enrolled and conducted scRNA-seq (Figure S15f-h). Baseline TAN abundance varied across these patients (Figure S15i, j). Interestingly, the patient with the highest baseline TAN abundance showed the greatest PSA decline, whereas the patient with relatively low TAN abundance showed persistent PSA elevation.

Collectively, this small exploratory cohort provides preliminary information on the safety of combined IGF1R and PD-1 blockade and descriptive evidence of possible clinical activity in mCRPC with bone metastases.

## Discussion

Bone metastasis of PCa marks a critical, often lethal, stage of the disease, for which existing immunotherapies, including ICIs, have demonstrated limited efficacy ^102,103,106^. A comprehensive understanding of the dynamic evolution of the BM microenvironment is therefore crucial for developing new therapeutic strategies. While previous single-cell studies have provided valuable snapshots, they have often been limited to specific disease stages or cell types, leaving the landscape of metastatic initiation and progression largely undefined ^15^. Our study addresses this critical gap by providing a multi-stage single-cell atlas of the human BM spanning benign conditions (Benign), clinically high-risk PCa without pathologically confirmed bone metastasis (HMP), and established bone metastasis (Mets). To avoid implying that all metastasis-negative patients would inevitably progress, we defined the intermediate cohort as having HMP. BM from these patients already exhibited metastasis-associated immune alterations, particularly within the neutrophil compartment, and these alterations became more pronounced in overt metastatic disease. Thus, our findings support the presence of PMN-like BM remodeling in clinically high-risk PCa, without assuming deterministic metastatic progression in every HMP patient. Updated follow-up, in which three of seven HMP patients subsequently developed radiologically confirmed bone metastases, provides longitudinal support for this interpretation. Moreover, for the first time, we delineate a detailed axis wherein primary tumor-associated IGF1 correlates with the emergence of a distinct population of CCR3^+^ TANs, which may play a key role in contributing to this pro-metastatic environment prior to the arrival of tumor cells in the BM microenvironment.

Our investigation into the cellular dynamics of the HMP-associated BM environment reveals a complex interplay of pro- and anti-tumor responses (Figure S15k). We observed an early enrichment of cytotoxic CD8Temra cells, likely representing an intrinsic response to control tumor dissemination. However, this anti-tumor response is gradually subverted as the disease progresses, with CD8 T cells in HMP displaying pre-exhaustion-associated features. Concurrently, the differentiation of anti-metastatic non-classical monocytes is reduced, which may further promote the migration of tumor cells and their seeding in distal sites. TANs were already detectable in the HMP stage and became progressively more abundant and inflammatory in metastatic disease, supporting a continuous model in which an early TAN-associated state is further reinforced after tumor colonization. Our results further suggest that TANs are prominent contributors to this immunosuppressive shift within the BM environment.

Our conditioned-medium and orthotopic prostate tumor models further support a role for tumor-associated soluble signals in inducing HMP-like BM alterations. Integrating the human observations with functional perturbation experiments identified IGF1 as a critical mediator of this process. Consistently, systemic IGF1 exposure promotes key features of the HMP bone niche in vivo, characterized by TAN accumulation, T cell dysfunction, and altered monocyte differentiation. Moreover, IGF1R blockade and neutrophil-specific *Igf1r* deletion attenuated TAN formation and metastatic progression, establishing a functionally important contribution of neutrophil-intrinsic IGF1R signaling.

The preferential emergence of this phenotype in BM despite systemic IGF1 exposure is also consistent with the higher IGF1R abundance and greater IGF1 responsiveness of BM neutrophils compared with circulating neutrophils. Delving deeper into the molecular underpinnings, we uncovered the IGF1-MAPK/ERK-KLF4 signaling cascade that promotes TAN-like neutrophil reprogramming. These TANs, in turn, mediate their immunosuppressive effects in part through IL-1β and ROS/NO-associated mechanisms. Together, these findings support a model in which tumor-associated systemic IGF1 contributes to TAN reprogramming and the development of an immunosuppressive BM environment that facilitates metastatic colonization.

These findings also have therapeutic implications. In our preclinical models, pharmacologically inhibiting the IGF1R attenuated TAN-associated BM remodeling and reduced metastatic colonization and subsequent tumor growth. Combined IGF1R inhibition with anti-PD-1 blockade significantly impedes the development of bone metastasis, providing a strong preclinical rationale for this novel combination therapy.

This is particularly relevant given that ICI-based therapies have consistently struggled in treatment of mCRPC. The landmark KEYNOTE-199 study focusing on ICI monotherapy reported PSA declines in only 15% of patients and a confirmed PSA50 response rate of just 6% ^102^. Similarly, the PD-L1 monotherapy arm of the CCTG IND.232 trial was closed early for futility after observing 0% ORR ^107^. Furthermore, attempts to sensitize mCRPC to ICIs by combining them with standard-of-care agents have been largely unsuccessful ^103,108^.

Our investigator-initiated trial extended this concept into patients with established mCRPC, a setting in which the pre-metastatic phase has already passed but the IGF1-TAN-associated program may remain therapeutically relevant. The trial provided preliminary safety information together with exploratory changes in PSA, ALP and lesion-level PET/CT metabolic activity. Pain outcomes were considered separately as symptomatic observations because IGF1R inhibition may directly affect bone-related symptoms. However, given the small, single-arm cohort, the absence of an anti-IGF1R monotherapy group and the lack of paired on-treatment BM samples, these findings cannot establish efficacy, therapeutic synergy or a TAN-mediated clinical mechanism. They should therefore be regarded as exploratory observations that warrant evaluation in larger, controlled studies.

Apart from identifying the early immune remodeling before metastasis, our study also provides a more comprehensive characterization of the established metastatic lesion itself, including its malignant cells, macrophages, and stromal components—populations often underrepresented in prior scRNA-seq studies of this setting. We identified COL5A2^+^/IGF1^+^ EMPs in metastatic lesions; their early-stage features and association with bone metastasis suggest a role in early metastatic seeding. Additionally, we identified a POSTN^+^/BGLAP^-^ OB1 osteoblast subtype that displayed a woven-bone-associated transcriptional phenotype consistent with the osteoblastic features of PCa bone lesions, offering molecular insights into the unique osteoblastic characteristic of PCa bone lesions. Furthermore, we found that both ECs and macrophages within the metastatic niche acquire a prostate-like transcriptional identity, highlighting a profound transcriptomic remodeling of the BM following metastasis. Whether these cells directly accompany tumor cells during dissemination or emerge as a result of post-colonization remodeling warrants further investigation.

It is important to acknowledge the limitations of our study, which also highlight the complexity of the metastatic process. First, the inherent difficulty in obtaining human BM samples, particularly from healthy and benign individuals, limits the statistical power for some comparisons. Specifically, patients in our benign group had underlying, non-malignant bone-related diseases, which may have introduced a baseline level of inflammation in their BM. Given the difficulty of obtaining BM from healthy donors, our analysis partly relied on multiple published healthy BM datasets to ensure the robustness of our analysis. Nevertheless, these approaches cannot fully substitute for prospectively matched healthy vertebral BM. In addition, neutrophil abundance and recovery are particularly sensitive to sample processing, and residual interpatient and technical variability cannot be excluded. Second, the HMP classification does not imply that every patient will inevitably develop bone metastasis. Four of the seven patients currently have approximately 2 years of follow-up, and longer longitudinal observation will be required to determine the clinical relationship between the identified BM alterations and future metastatic progression. Third, the liver is the major physiological source of circulating IGF1, and the observed decrease in circulating IGF1 bioavailability after prostatectomy does not directly trace the molecular source of circulating IGF1. Our tumor-cell Igf1 perturbation experiments establish that tumor-derived IGF1 can be functionally relevant, but they do not quantify the relative contributions of tumor-derived and hepatic IGF1 in vivo. Fourth, repeated conditioned-medium administration is a simplified model that does not reproduce the concentration gradients, pharmacokinetics or full cellular complexity generated by an intact primary tumor. Fifth, although neutrophil-specific *Igf1r* deletion establishes an important causal role for neutrophil-intrinsic signaling, systemic IGF1R inhibitors may also act on tumor, osteoblastic, endothelial and other immune compartments. In addition, although IGF1R blockade reduces CCR3^+^ TAN accumulation, our endpoint analyses cannot determine whether pre-existing TANs undergo phenotypic reversion, are selectively depleted, or are replaced by newly generated neutrophils after treatment. Sixth, although functional experiments support contributions of IL-1β, ROS and nitric oxide to TAN-mediated T cell suppression, other mediators are likely involved, and the proposed LGALS9-HAVCR2-NR4A1 relationship between TANs and monocyte differentiation remains to be functionally tested. Finally, the investigator-initiated trial addressed treatment of established mCRPC rather than prevention of metastatic initiation. It was a small, single-arm study with safety as the primary endpoint, lacked an anti-IGF1R monotherapy arm and did not include matched on-treatment BM profiling. These features preclude conclusions about therapeutic synergy or the cellular basis of the observed clinical changes. Accordingly, validation of the therapeutic strategy and its underlying clinical mechanism will require larger, controlled studies with longitudinal translational sampling.

In summary, our human and preclinical data support a model in which tumor-associated systemic IGF1 contributes to the reprogramming of BM neutrophils toward a TAN state, promoting an immunosuppressive environment associated with subsequent metastatic colonization. Following tumor arrival, the IGF1-TAN-associated program persists alongside broader malignant, stromal and vascular remodeling of the established metastatic niche. Our functional and therapeutic studies support the IGF1-TAN pathway as a contributor to metastatic progression and a potential therapeutic vulnerability. Our findings provide a rationale for combining IGF1R inhibitors with immune checkpoint blockade in heavily pretreated mCRPC. Plasma IGF1 activity and BM TAN abundance may also represent candidate biomarkers, although their predictive and patient-stratification value will require prospective validation.

## Materials and Methods

### Human participants and specimen collection

#### 1 Human cohorts and ethics

The main BM cohort comprised Benign (n = 6), high-risk bone metastatic potential (HMP; n = 7) and established bone metastasis (Mets; n = 7) groups. Benign BM was obtained during clinically indicated orthopaedic procedures for non-malignant conditions from patients without known systemic inflammatory disease. HMP patients had high-risk localised or locally advanced PCa, no radiologically confirmed bone metastasis at BM sampling and no previous systemic therapy, including ADT. Mets patients had radiologically or histologically confirmed bone metastasis and heterogeneous treatment histories. Additional paired prostate and BM specimens were analysed in the extended cohorts. Clinical characteristics, ADT exposure status at BM collection and follow-up are provided in Table S1. All human tissue acquisition procedures were approved by the Institutional Review Boards of Shanghai Changzheng Hospital and Shanghai Changhai Hospital. All participants provided written informed consent before specimen collection.

#### 2 Collection of BM and prostate specimens

BM was collected by biopsy or during surgery. For biopsy, the target bone was exposed under general anaesthesia and a Jamshidi needle was introduced directly into the bone body; BM was aspirated immediately after cannulation to minimise dilution by PB or surgical irrigation. Surgical BM specimens were obtained from patients undergoing procedures for lumbar disc herniation, lumbar spinal stenosis or PCa bone metastasis. Prostate samples were collected as previously described ^30^, including specimens obtained by radical prostatectomy or needle biopsy.

#### 3 Peripheral blood collection and CD8 T-cell isolation

PB samples were collected from healthy male donors, and human CD8 T cells were isolated using the MojoSort^TM^ Human CD8 T Cell Isolation Kit (BioLegend, #480011) according to the manufacturer’s instructions. The isolated CD8 T cells were washed with MojoSort^TM^ buffer and immediately used for subsequent experiments.

#### 4 Public datasets

Publicly available single-cell and bulk-transcriptomic datasets were used for external validation, cross-dataset integration and survival analyses. Accession numbers and sample characteristics are summarized in Table S9. When raw or count-level single-cell data were available, they were processed using the same quality-control, integration and annotation framework described below.

### Investigator-initiated clinical trial

#### 1 Study design and oversight

This was a single-centre, open-label, single-arm, early phase I investigator-initiated trial conducted at Shanghai Changzheng Hospital. The study commenced on 10 March 2025 and reached primary completion on 11 November 2025. The trial was prospectively registered at ClinicalTrials.gov (NCT06866548). The protocol was approved by the Institutional Review Board of Shanghai Changzheng Hospital (2025SL023), and all participants provided written informed consent before enrolment. The study was conducted in accordance with the Declaration of Helsinki and Good Clinical Practice.

#### 2 Participants and treatment

Seven men aged 18–85 years with histologically confirmed metastatic castration-resistant prostate adenocarcinoma (mCRPC) and radiographic evidence of bone metastases on PSMA-PET/CT or bone scintigraphy were enrolled. Eligibility required castrate serum testosterone levels (<50 ng/dL or <1.7 nmol/L), an Eastern Cooperative Oncology Group (ECOG) performance status of ≤2, a life expectancy of ≥6 months and adequate hematological, renal and hepatic function, including hemoglobin >9.0 g/dL, serum creatinine <1.5× the upper limit of normal (ULN), and alanine aminotransferase and aspartate aminotransferase <3× ULN. Key exclusion criteria included predominant non-adenocarcinoma histology, active infection requiring parenteral antibiotics or associated with fever >38.1 °C within 1 week before enrolment, systemic immunosuppressive therapy within 14 days before study-drug administration, severe cardiovascular disease, previous anti-IGF1R antibody or immune-checkpoint inhibitor therapy, known intolerance to either class of agent, uncontrolled major systemic disease, and investigator-assessed unsuitability for study participation. Full eligibility criteria and laboratory thresholds are provided in the approved study protocol.

Patients continued ongoing ADT throughout the study and received intravenous anti-IGF1R antibody teprotumumab (Teprotumumab N01 Injection; Innovent Biologics) together with the anti-PD-1 antibody tislelizumab (Tislelizumab Injection; BeiGene). Teprotumumab was administered at 10 mg/kg for the first dose and 20 mg/kg every 3 weeks thereafter; tislelizumab was administered at 200 mg every 3 weeks. The planned minimum treatment duration was 12 weeks (four cycles), with treatment continuation permitted in the absence of disease progression or unacceptable toxicity.

#### 3 Assessments, endpoints and analysis populations

Disease assessments were performed according to Prostate Cancer Clinical Trials Working Group 3 (PCWG3) criteria and RECIST v1.1, where applicable. Serum prostate-specific antigen (PSA), alkaline phosphatase (ALP), PSMA-PET/CT and other clinically indicated imaging assessments were performed at baseline and during follow-up, including the prespecified assessment at day 84 ± 3. Pain severity was assessed using the Brief Pain Inventory–Short Form (BPI-SF). Adverse events were recorded throughout treatment and follow-up and graded according to the National Cancer Institute Common Terminology Criteria for Adverse Events (NCI CTCAE), version 5.0.

The primary endpoint was safety, assessed by the incidence and severity of adverse events according to NCI CTCAE v5.0. Secondary endpoints included PSA progression-free survival (PSA-PFS), radiographic progression-free survival (rPFS), PSA response, disease-control rate (DCR), objective-response rate (ORR), overall survival (OS), skeletal-related events and pain severity.

No formal statistical power calculation was performed because of the exploratory nature of this early-phase study. The safety analysis set included all patients who received at least one dose of either study drug. The efficacy-evaluable set included patients who received study treatment and had at least one post-baseline efficacy assessment. The number of patients contributing to each clinical endpoint is reported in the corresponding figure or table.

Time-to-event endpoints, including PSA-PFS, rPFS and OS, were estimated using the Kaplan–Meier method, with median times and 95% confidence intervals reported where estimable. PSA, ALP and pain trajectories were summarized descriptively because of the small cohort size and incomplete repeated measurements. Other clinical outcomes, including PSA response, DCR and ORR, were summarized using descriptive statistics. Statistical analyses were performed using Prism (version 10.3.1) or R (version 4.2.2), as appropriate.

### Single-cell preparation and sequencing

#### 1 Single-cell Dissociation

The tissues were surgically removed and kept in MACS Tissue Storage Solution (Miltenyi Biotec) until processing. The tissue samples were processed as described below. Briefly, samples were first washed with phosphate-buffered saline (PBS), minced into small pieces (approximately 1 mm^3^) on ice and enzymatically digested with 1 mg/mL collagenase I, 2 mg/mL collagenase IV, 1 mg/mL dispase and 30 U/mL DNase I for 30 min at 37 °C, with agitation. After digestion, samples were sieved through a 70 µm cell strainer, and centrifuged at 300 × g for 5 min. After washing with PBS containing 0.04% BSA, the cell pellets were re-suspended in PBS containing 0.04% BSA and re-filtered through a 40μm cell strainer. Dissociated single cells were then stained for viability assessment using Calcein-AM (Thermo Fisher Scientific) and Draq7 (BD Biosciences).

#### 2 Single-cell RNA Sequencing

BD Rhapsody system was used to capture the transcriptomic information of the single cells. Single-cell capture was achieved by random distribution of a single-cell suspension across >200,000 microwells through a limited dilution approach. Beads with oligonucleotide barcodes were added to saturation so that a bead was paired with a cell in a microwell. The cells were lysed in the microwell to hybridize mRNA molecules to barcoded capture oligos on the beads. Beads were collected into a single tube for reverse transcription and ExoI digestion. Upon cDNA synthesis, each cDNA molecule was tagged on the 5′ end (that is, the 3′ end of a mRNA transcript) with a unique molecular identifier (UMI) and cell barcode indicating its cell of origin. Whole transcriptome libraries were prepared using the BD Rhapsody single-cell whole-transcriptome amplification (WTA) workflow including random priming and extension (RPE), RPE amplification PCR and WTA index PCR. The libraries were quantified using a High Sensitivity DNA chip (Agilent) on a Bioanalyzer 2200 and the Qubit High Sensitivity DNA assay (Thermo Fisher Scientific). Sequencing was performed by Illumina sequencer (Illumina, San Diego, CA) on a 150 bp paired-end run.

### Single-cell RNA-seq data analysis

#### 1 Preprocessing of the scRNA-seq data

We applied fastp (v0.23.4) with default parameter filtering the adaptor sequences and removed the low quality reads to achieve the clean data ^109^. UMI-tools was applied for Single-cell transcriptome analysis to identify the cell barcode whitelist ^110^. The UMI-based clean data was mapped to the human genome (Ensembl version 91) utilizing STAR (2.7.10b) mapping with customized parameters from UMI-tools (v1.1.2) standard pipeline to obtain the UMIs counts of each sample ^111^.

After loading UMIs counts matrix as a Raw Seurat (v4.4.0) object, we utilized three steps to generate a clean matrix:

1. Initial Filtering and Doublet Removal:

Cells with fewer than 300 features (nFeature < 300) were removed.

Doublets were identified and removed using DoubletFinder (v2.0.3). The expected doublet number was set to 6% of the recovered cells (nExp = round(0.06 × nCells)).

2. Major Cell Type Annotation and Further Filtering:

Major cell types were annotated, recognizing neutrophils and other cell types.

Cells, except neutrophils, were further filtered under the criterion of nFeature > 800, as previous research reported that neutrophils express a low number of genes ^112^.

3. Correction for Ambient RNA Contamination:

Ambient RNA contamination was corrected using SoupX (v1.6.2). The detailed process is described below.

After completing these three steps, the cleaned counts matrix was used for subsequent procedures.

#### 2 Ambient RNA contamination correction

Potential ambient-RNA marker genes were defined based on lineage-restricted expression patterns observed across the dataset, including:

Plasma-related: IGKC, IGHA1, IGHA2, IGHG1

Erythroid-related: HBB, HBD, HBA2 Neutrophil-related: MPO, ELANE, LTF

Mesenchymal-related: COL1A1, COL1A2, COL3A1

The contamination rate in each sample was calculated using the function estimateNonExpressing from SoupX (v1.6.2). If certain samples did not contain any of the cell types contributing to ambient RNA, these genes were ignored during calculation. We then used SoupX to correct for ambient RNA contamination separately in each sample. The background expression (argument: tod) was set to the raw matrix, which did not undergo any filtering. Other parameters were set to their default values.

#### 3 Data integration, batch effect correction, dimension reduction, and unsupervised clustering

The resulting gene expression matrix, once filtered, was normalized using the NormalizeData function, and the logarithm-transformed normalized matrix was employed for downstream analyses. Batch effect correction was performed using the RunFastMNN function from the SeuratWrappers (v0.3.1) package in R, treating each sample as a separate batch. The generated MNN space was subsequently used for graph-based clustering, utilizing the FindNeighbors function in Seurat to construct the Shared Nearest Neighbor Graph. Unsupervised clustering was then conducted with the FindClusters function in Seurat. For visualization, dimensionality reduction was achieved using Uniform Manifold Approximation and Projection (UMAP) through the RunUMAP function in Seurat, with the reduction parameter set to “mnn”. Differential gene expression analysis in different subsets was generated through the Wilcoxon Rank Sum test, using the FindMarkers function. For integration with published scRNA-seq datasets, analyses were restricted to genes shared across the datasets, after which the same fastMNN-based batch-correction and Seurat clustering workflow was applied.

To mitigate the influence of batch effects on gene expression comparisons, in addition to using the Seurat functions ‘FindMarkers’ and ‘FindAllMarkers’, we also evaluated the difference in the proportion of cells with detectable expression (pct.diff) between clusters. Specifically, when comparing a gene’s expression between clusters, we computed the percentile of positive expression within clusters using Fisher’s exact test.

#### 4 Regulon (transcription factor activity) analysis and regulon activities-based clustering

We used pySCENIC (v0.12.1) to estimate activated regulons with default settings. The AUCell scores generated from the standard pySCENIC pipeline were then loaded for subsequent regulon-related analyses. Due to the high inter-individual heterogeneity of malignant epithelial cells in the BM, fastMNN failed to integrate the transcriptome data effectively. Therefore, as suggested in the pySCENIC tutorial, we constructed a new “Assay” using regulon activity scores as the normalized expression matrix. We then followed the previous Seurat pipeline to conduct batch effect correction, graph-based clustering, and visualization.

#### 5 NicheNet analysis

To identify potential ligands capable of inducing specific signature gene expression, we used the predict_ligand_activities function in R package nichenetr (v2.0.4) ^113^. For the differentiation of OB1 fro m preOB, differentially expressed genes (DEGs) generated by comparing OB1 with preOB were selected according to the criteria: avg_log2FC > 0.5 and p_val_adj < 0.05. For the transition from CapEC1 to CapEC2, DEGs with avg_log2FC > 0.5 and p_val_adj < 0.01 were selected as the targeted gene signature. For TAN formation, DEGs with avg_log2FC > 0.5 and p_val_adj < 0.01 between mNeu and TAN were selected. These targeted gene signatures were then used for subsequent NicheNet analysis.

#### 6 Integration of tumor-derived secreted ligand screening with BM neutrophil response analysis

To explore potential tumor-derived secreted signals associated with BM neutrophil remodeling, we developed a two-step ligand prioritization workflow integrating ligand-receptor annotation with receiver-cell transcriptional response analysis. This analysis was designed for hypothesis generation and was not used as direct evidence of physical cell-cell communication or endocrine signaling between the primary tumor and BM. First, we identified ligands that were enriched in primary PCa cells relative to the comparator group and annotated as secreted signaling molecules. Ligands requiring direct cell-cell contact or extracellular matrix-associated interactions were excluded. This tumor-enrichment filter was used to capture tumor-associated ligand changes, but was not interpreted as evidence of ligand abundance, protein secretion, or circulating concentration. For each retained ligand, the expression of its cognate receptor in BM neutrophil populations was assessed to identify secreted ligand-receptor pairs with potential accessibility to neutrophils. Second, we evaluated whether these candidate ligand-receptor pairs were associated with the transcriptional response observed in BM neutrophils. Differentially expressed genes in BM neutrophils were used as the receiver-cell response signature, and ligand activity was estimated using NicheNet based on the ability of each candidate ligand to explain this neutrophil transcriptional program through prior ligand-receptor, signaling, and gene regulatory networks. Ligands supported by both tumor-side secreted ligand enrichment and neutrophil-side transcriptional response were prioritized for downstream validation. This workflow was used to narrow the list of potential systemic mediators. The biological relevance of IGF1 was subsequently tested experimentally using tumor-CM, recombinant IGF1 stimulation, IGF1R blockade, and neutrophil-specific *Igf1r*-deficient mouse models.

#### 7 Trajectory constructions and differentiation analysis

To explore the cellular differentiation trajectory, we utilized Monocle2 (v2.24.0) ^114^ and destiny (v3.20.0) ^115^. Specifically, for Monocle2, we first identified critical gene lists by FindAllMarkers, and selected genes by criteria: avg_Log2FC>0.5&p_val_adj<0.01. Method “DDRTree” was selected when conducting dimension reduction using the function “reduceDimension”. Pseudotime was calculated after manually setting root cells by function “orderCells”. For the diffusion map generated by destiny, we used the function DiffusionMap and selected the 1-20 dims of mnn reduction to construct the diffusion map.

To evaluate the stemness of cells, we used R package CytoTRACE2 (v1.0.0) ^116^. To avoid batch effect, we used fastMNN corrected normalized counts as input matrix.

#### 8 Scoring cells using gene expression signatures

Gene signature activities were calculated using AddModuleScore with default parameters from Seurat. Gene sets were collected from Molecular Signatures Database (MSigDB; v2023.1.Hs, v2023.1.Mm) and published research (Table S2), mainly including gene sets from: Kyoto Encyclopedia of Genes and Genomes (KEGG), the Pathway Interaction Database (PID), BioCarta, WikiPathways (WP), Gene ontology (GO), Reactome, The Human Phenotype Ontology (HP), Hallmark and Hsiao’s housekeeping gene sets ^117^.

#### 9 Sample-wise validation of the TAN state

To assess inter-patient reproducibility of the TAN state, neutrophils from each HMP and Mets BM sample were re-analysed independently using the same normalization, dimensional-reduction and clustering framework. TAN-related clusters were identified based on enrichment of the predefined TAN signature together with expression of representative TAN-associated genes, including *CCR3*, *ICAM1* and *IL1B*. TAN abundance was then recalculated using these sample-wise assignments and compared with the annotation obtained from the integrated analysis.

#### 10 Group prevalence of cell subsets

Cell-type proportions and mean signature scores were calculated separately for each sample before group comparison. Correlations between cell populations or sample-level programmes were calculated across matched samples using Pearson or Spearman correlation as indicated. To assess the prevalence of cell types across groups, log-transformed odds ratios were calculated using two-sided Fisher’s exact tests ^118^. For each cell type-group combination, a 2 × 2 contingency table was constructed comparing the number of cells of the indicated cell type within versus outside the group of interest against all other cell types. Odds ratios were used to quantify relative cell-type enrichment or depletion across groups, and P values were adjusted for multiple comparisons using the Benjamini-Hochberg method.

#### 11 Infer Copy number variation (CNV) using inferCNV

CNV patterns were identified using inferCNV from R package infercnv (v1.17.0) and followed their standard pipeline. For the analysis of BM data in the main cohort, endothelial cells were used as reference cells because of the absence of normal epithelial cells in BM.

#### 12 Malignant epithelial cells identification

To distinguish malignant from non-malignant epithelial cells within the prostate tissue, we first established a non-malignant reference population. Through a systematic comparison of epithelial cells from our previous dataset (GSE141445) and public normal prostate datasets (GSE201333), we identified cells expressing canonical markers for normal prostate lineages. These included basal cells (*KRT5*^+^), ductal cells (*SCGB1A1*^+^, *SCGB3A1*^+^) ^119^, and a normal luminal cell population characterized by high expression of genes previously demonstrated to be specific to normal prostate tissue, such as *LGALS3*, *MSMB*, *SLC14A1*, and *CD177* ^120–123^. Subsequently, we utilized the Infercnv package to assess large-scale CNVs. The identified non-malignant cell populations were collectively used as the reference group. Epithelial cells exhibiting distinct CNV patterns relative to this reference set were thus validated as malignant.

#### 13 NMF-based clustering

The NMF-based meta-programs analysis was adapted from the procedures of previous research ^93^. Initially, mitochondrial genes, immunoglobulin genes, and ribosomal protein genes were excluded from the expression matrix. To address inter-sample heterogeneity, we included only those genes expressed in more than half of the samples and selected the top 7,000 genes based on average expression levels across all epithelial cells. The data were then normalized to counts per million (CPM) using the ‘NormalizeDatà function, with the ‘normalization.method’ parameter set to “RC” and the ‘scale.factor’ set to 1e6. The normalized data were divided by 10, incremented by 1, and log-transformed. Subsequently, the expression data were centered, with negative values set to zero.

NMF clustering was performed separately for each sample using the function nnmf from R package NNLM (v0.4.4), with the number of clusters (k) ranging from 4 to 9, generating 39 NMF programs. Each NMF program was summarized by the top 50 genes based on NMF coefficients. Intra-sample robust programs were identified by recognizing programs with at least 60% gene overlap (30 out of 50 genes) with any other program from the same sample. These robust programs were then evaluated for inter-sample robustness, where programs with at least 20% gene overlap (10 out of 50 genes) with any other programs from different samples were recognized as robust among samples.

To obtain unique programs, we filtered these robust programs by estimating their similarity. Programs with more than 50% gene overlap (25 out of 50 genes) with any other program were removed. For identifying meta-programs, we initiated the first program as the meta-program. Any subsequent programs with at least 20% overlap (10 out of 50 genes) with the initiated meta-program were added to it sequentially. This process resulted in a large meta-program containing numerous genes. We then sorted the overlapping genes by frequency and selected the top 50 genes as the final meta-program. The remaining programs followed the same procedure. Meta-programs containing fewer than three programs were excluded from further analysis.

#### 14 Metabolism analysis using Compass

To generate reaction activity, we utilized the Compass (v0.9.10.2) ^124^. Initially, UMI counts were transformed into CPM as previously described to create the input matrix. The model used was Recon 2 ^125^, with the parameters set as follows: lambda = 0, num_neighbors = 30, and microcluster_size = 30. All other parameters were set to their default values. This process resulted in the identification of 516 microclusters.

The transformation of reaction penalties into reaction activities followed the methodology outlined in the Compass tutorial. If 90% of the cells within a microcluster belonged to the same cell type, we retained and annotated the microcluster as that cell type. The resulting microcluster-to-reaction activities matrix was then used as the “data” to construct a Seurat object.

We identified significantly regulated reactions among different cell types by Wilcoxon Rank Sum test. Based on the subsystem classification of reactions, we enriched these reactions using the enricher function from the R package clusterProfiler (v4.6.2). This analysis revealed metabolic alterations within specific subsystems, providing insights into metabolic reprogramming across different cell types.

#### 15 Gene signature scoring in bulk RNA-seq samples

We used the function gsva in R package GSVA (v1.46.0) to calculate ssGSEA score of specific gene signature in bulk RNA-seq data. The parameter “Method” was set to “ssgsea”. For gene signatures of specific cell type, we ranked DEGs using their avg_log2FC, and selected the top 50 genes as their signatures.

#### 16 Similarity analysis of cell subsets

Similarity analysis was conducted as in previous research ^126^. Briefly, average expression levels of the 2,000 highly variable genes were calculated using Seurat function AverageExpression in each cell cluster and were used to calculate the pairwise Pearson or Spearman correlation.

#### 17 Survival analysis in the SU2C metastatic PCa (mPCa) bone samples

To avoid gene expressions contributed by other cell components, we first selected epithelial specific genes. using FindMarkers function in Seurat, we compared epithelial cells with all other cells in BM. The resulting DEGs were filtered using the following criteria to generate a list of metastatic epithelial specific DEGs: pct.2<0.1&log2FC>0.5&p_val_adj<0.05. Then we generate subtype specific DEGs comparing EMPs and LMPs using the FindAllMarkers function in Seurat while maintaining overlaps with epithelial specific DEGs. Finally, we selected the top 50 genes from DEGs as EMP and LMP gene signature respectively.

For subsequent analysis, only bone metastasis samples from the SU2C cohort were included. Each sample was assigned an EMP and LMP score using ssGSEA and classified into the respective subgroup based on their higher score. For survival analysis, the statistical test was conducted using R package coin (v1.4.3), The Gehan-Breslow test was used to evaluate significance. Kaplan-Meier survival curves were plotted using the ggsurvplot function in the R package survminer (v.0.4.9).

#### 18 Cellular interaction analysis by CellphoneDB

Cell-cell communication networks were assessed using CellphoneDB (v4) ^127^. Input parameters for the statistical module included a raw gene expression matrix exported from the Seurat object paired with cell type metadata. Interaction frequencies between cellular subsets were visualized via the plot_cpdb_heatmap function (ktplots v2.1.0), while ligand-receptor interaction strengths were predicted based on average expression levels and graphically represented. Statistically significant pairs (P<0.05) were filtered for downstream analysis. This standard CellPhoneDB workflow was used for within-dataset or within-tissue ligand-receptor analyses.

#### 19 RNA velocity analysis and pseudo-time trajectory inference

Spliced and unspliced counts were generated from fastq files using python package Velocyto (v0.17.17). The following analyses were conducted in R using R package anndata (v0.7.5.6) ^128^, reticulate and python package scvelo (v0.2.5) ^129,130^. First, we removed tissue dissociation-associated genes from counts matrix ^131^. Then we filtered and normalized the counts matrix using the function scv.pp.filter_and_normalize, set min_shared_counts=20, n_top_genes=2000. Next, a k nearest-neighbor graph (k=30) was built using top 20 MNN components generated by fastMNN. For each cell, the moments of normalized spliced/unspliced counts were computed using the 30 nearest neighbors by function scv.pp.moments. The estimated velocities were used to construct a velocity graph representing the transition probabilities among cells by function scvelo.tl.velocity_graph.

Finally, the velocity graph was used to embed the velocities into the UMAP or Diffusion map by the function scv.pl.velocity_embedding_grid or scv.pl.velocity_embedding_stream.

### Bulk RNA-seq and ATAC-seq

#### 1 Bulk RNA sequencing

Total RNA was extracted from control and IGF1-treated mouse BM neutrophils using RNA isolater Total RNA Extraction Reagent. RNA quality was assessed using Agilent 2100, and libraries were generated using VAHTS Universal V10 RNA-seq Library Prep Kit and sequenced as paired-end reads on DNASEQ-T7 using PE150. Two mice were analysed per condition, with each mouse serving as an independent biological replicate.

Raw paired-end sequencing reads were first subjected to quality control using fastp to remove adapter sequences and filter low-quality reads. The resulting clean reads were then aligned to the mouse reference genome (GRCm39), with gene annotations from GENCODE release M35, using the STAR.

Gene-level read counts were quantified from the aligned BAM files using featureCounts (v2.1.1) ^132^. Differential gene expression analysis between experimental conditions was performed with the DESeq2 R package (v1.38.3) ^133^. Prior to analysis, genes with low expression (fewer than 10 reads in at least 3 samples) were removed. An adjusted p value < 0.05 was used as the threshold for statistical significance. For visualization and downstream analyses, the raw counts were normalized using the variance-stabilizing transformation (VST) function in DESeq2.

#### 2 ATAC-sequencing

Purified mouse BM neutrophils were treated with vehicle or IGF1 as described above. ATAC-seq was performed according to ATAC-seq protocol by Shanghai Jiayin Biotechnology Ltd. In brief, cells were harvested from cell culture and lysed in a lysis buffer. The Nextera DNA Library Preparation Kit (Illumina) was used to perform the transposition according to the manufacturer’s manual. 50,000 nuclei were pelted and resuspended with transposase, for 30 minutes at 37. The transposed DNA fragments were purified immediately after with a MinElute PCR Purification Kit (Qiagen). Samples were PCR-amplified using 1× NEBNext High-Fidelity PCR Master Mix (New England Biolabs, MA). Subsequent libraries were purified with the MinElute PCR Purification Kit (Qiagen) and subjected to sequencing on Illumina NovaSeq 6000 using PE150. Each condition contained two biologically independent neutrophil preparations that were cultured and treated separately.

Raw paired-end ATAC-seq reads underwent quality control using fastp to trim adapters and filter low-quality sequences. The clean reads were then aligned to the mouse reference genome (GRCm39) using bwa-mem2 (v2.3) ^134^. The resulting SAM files were converted to BAM format, sorted, and indexed using Samtools (v1.17) ^135^. To reduce noise from non-nuclear DNA, reads mapping to the mitochondrial genome (chrM) were removed. PCR duplicates were subsequently marked and removed using Picard (v3.4.0) MarkDuplicates. Finally, a high-confidence set of alignments was generated by filtering for properly paired reads with a mapping quality (MAPQ) score > 30 using Samtools. Chromatin accessibility peaks were called on the final filtered BAM files for each sample using MACS2 (2.2.9.1) with parameters suitable for ATAC-seq data (--nomodel --shift -100 --extsize 200 -g mm) ^136^. To identify differentially accessible regions (DARs) between experimental conditions, we used the DiffBind R package (3.18.0) ^137^. A consensus peakset was generated, and read counts within these peaks were quantified. Statistical analysis was performed using the DESeq2 method integrated within DiffBind. DARs were considered significant at a False Discovery Rate (FDR) < 0.05 and an absolute log2 fold-change > 1.

To identify TF binding motifs enriched within significant DARs, we utilized the findMotifsGenome.pl tool from the HOMER suite (v5.1) ^138^. The genomic location of DARs relative to known gene features (promoters, exons, etc.) was determined using HOMER’s annotatePeaks.pl tool.

To identify TFs that could link MAPK/ERK and KLF4 signaling, we compiled a list of known downstream TFs of the ERK pathway ^99^. We then scanned the promoter region of the mouse *Klf4* gene (2 kb upstream of the transcription start site) for potential binding sites of these TFs using the JASPAR database ^139^. From the resulting candidates, c-MYC was selected for further validation as it was both predicted to bind the KLF4 promoter and was transcriptionally upregulated in TANs upon IGF1 stimulation (Table S4).

### Cell lines and conditioned medium

TRAMP-C1 cells were obtained from ATCC and maintained in DMEM supplemented with 10% FBS, 100 U/ml penicillin, 0.005 mg ml−1 bovine insulin and 10 nM dehydroepiandrosterone at 37 °C with 5% CO2. MyC-CaP cells were kindly provided by Binzhi Qian and cultured under standard conditions as previously described ^74^. To generate luciferase-expressing cell lines, MyC-CaP cells were transduced with a lentiviral vector carrying a firefly luciferase expression cassette driven by the CMV enhancer–MCS– 3×FLAG–hEF1/HTLV promoter and linked to a T2A-puromycin selection marker. Stable MyC-CaP-LUC cells were selected with puromycin. TRAMP-C1 cells were transduced with the Lenti-CMV-Luc-EF1-Blasticidin vector, and stable TRAMP-C1-LUC cells were established by blasticidin selection. Luciferase activity was confirmed before in vivo experiments. All cell lines were confirmed to be mycoplasma-free before use. Stable *Igf1*-knockdown TRAMP-C1 cells were generated using lentiviral shRNA constructs targeting mouse *Igf1* cloned into the GV493 vector (hU6-shRNA-CBh-gcGFP-IRES-puromycin). The *Igf1*-targeting shRNA sequence was 5′-TGATCTGAGGAGACTGGAGAT-3′. A scrambled shRNA construct carrying the sequence 5′-TTCTCCGAACGTGTCACGT-3′ served as the negative control. Mouse Igf1-overexpressing lentivirus was generated by GeneChem (Shanghai, China) using the GV492 lentiviral vector system (Ubi-MCS-3FLAG-CBh-gcGFP-IRES-puromycin). Cells were transduced with lentivirus and selected with puromycin to establish stable Igf1-overexpressing cells.

CM was collected according to our institutional protocol as described in previous studies ^140,141^. Briefly, TRAMP-C1 cells were cultured in standard complete medium until reaching 85% confluence. The cells were then gently washed with PBS, and the medium was replaced with serum-free DMEM. After 48 hours of culture, the serum-free CM was collected, filtered (0.22 μm) to remove cell debris, and concentrated using a Millipore protein concentrator (Millipore, UFC9003). To control for any baseline medium effects, blank serum-free DMEM (without cells) was incubated, filtered, and concentrated in parallel under identical conditions to serve as the matched blank-medium control. Medium was centrifuged at 3500 × g for about 8 minutes until triple the concentration. The concentrated CM was aliquoted to avoid repeated freezing and thawing and stored at −80°C for use within 3 months.

### Mouse models

#### 1 Animals and genetic model

Wild-type C57BL/6J mice, *Ly6g*^CreERT2/+^ mice and *Igf1r*^fl/fl^ mice were obtained from GemPharmatech (Nanjing, China). Tamoxifen-inducible neutrophil-specific *Igf1r* conditional knockout (cKO) mice were generated by crossing *Ly6g*^CreERT2/+^ mice with *Igf1r*^fl/fl^ mice.

*Ly6g*^CreERT2/+^;*Igf1r*^fl/fl^ mice were used as cKO mice, whereas Cre-negative *Ly6g*^+/+^;*Igf1r*^fl/fl^ littermates were used as controls. Genotypes were determined by PCR according to the manufacturer’s instructions. To induce Cre-mediated recombination and neutrophil-specific deletion of *Igf1r*, cKO mice were administered tamoxifen before the indicated experiments; control mice received the same tamoxifen treatment regimen. Tamoxifen (200 mg/kg) was administered by intraperitoneal injection once daily for 5 consecutive days, and experiments were initiated 1 day after the final administration.

Male mice of the ages indicated for each experiment were maintained under specific-pathogen-free conditions with controlled temperature and humidity and a 12-h light-dark cycle, with ad libitum access to food and water. Mice were randomly allocated to experimental groups where applicable. Sample sizes are reported in the corresponding figure legends.

All animal experiments were performed in accordance with institutional guidelines and were approved by the Animal Ethics Committee of Shanghai Changzheng Hospital.

#### 2 BM preconditioning models

For CM-induced BM preconditioning, 5-week-old male C57BL/6J mice received daily intraperitoneal injections of 300 µl concentrated TRAMP-C1 CM or identically processed blank medium for 2 weeks. At the indicated endpoint, mice were euthanized and PB and bone/BM samples were collected for subsequent analyses.

For recombinant IGF1 preconditioning, mice received recombinant IGF1 (Sino Biological, 10598-HNAY1; 1 mg/kg, intraperitoneally, once daily), prepared in saline, for 2 weeks. Picropodophyllin (PPP; MCE, HY-15494; 20 mg/kg, intraperitoneally, once daily), formulated in 15% Cremophor EL and 85% saline (v/v), was administered throughout the preconditioning period until the day before sample collection where indicated. Control mice received matched administrations of saline and PPP vehicle, whereas mice in the IGF1-only group received the matched PPP vehicle according to the same dosing schedule.

#### 3 Orthotopic prostate tumor model

Luciferase-expressing TRAMP-C1 cells (approximately 4 × 10^5^ cells per mouse in 25 μl injection volume containing 20% Matrigel [Corning, 356231]) were injected into the anterior prostate lobe of male C57BL/6J mice; sham-operated mice received vehicle alone. Mice were analysed after 2 weeks, before detectable bone metastasis, and PB, BM, spleen, liver and lung were collected for IGF1 and immune analyses.

#### 4 Bone metastasis and treatment models

For the TRAMP-C1 model, mice received 5 × 10^4^ TRAMP-C1-LUC cells by left-ventricular intracardiac injection. For independent validation, 1 × 10^5^ MyC-CaP-LUC cells were injected using the same route.

Bone metastatic tumor burden was monitored by bioluminescence imaging (BLI) every 3 days. Before imaging, mice received D-luciferin potassium (3 mg/kg; Meilunbio, MB1834), and bioluminescence signals were acquired using a bioluminescence imaging system (BLT Photon Technology). To obtain an integrated measure of tumor burden over time, the area under the curve (AUC) was calculated from serial BLI signal measurements for each mouse. BLI signals were natural-log transformed as ln(signal + 1) before longitudinal visualization and AUC calculation. AUC values were then used for comparisons of tumor burden between experimental groups ^142^.

For BM-preconditioning metastasis experiments, mice first received CM or recombinant IGF1 for 2 weeks as described above, followed by intracardiac injection of TRAMP-C1-LUC cells. PPP (MCE, HY-15494; 15 mg/kg per day, intraperitoneally), formulated in 15% Cremophor EL and 85% saline (v/v), was administered during the preconditioning phase where indicated. Groups not receiving PPP received the matched PPP vehicle according to the same dosing schedule ^143^.

For the mouse BM scRNA-seq experiment shown in Figure 3p-w, three mice were included in each of the saline-control, IGF1-induced HMP-like and IGF1-preconditioned TRAMP-C1 metastatic groups. BM cells from the three mice within each group were pooled to generate one sequencing sample per group before single-cell capture and library preparation.

For neutrophil-specific *Igf1r* cKO metastasis experiments, tamoxifen-treated cKO and littermate-control mice underwent IGF1 preconditioning before intracardiac TRAMP-C1-LUC challenge. For the therapeutic bone-metastasis model, TRAMP-C1-LUC cells were introduced without prior CM or IGF1 preconditioning, and treatment was initiated 1 day after intracardiac tumor-cell injection, anti-IGF1R antibody ganitumab (MCE, HY-P99294, CAS 905703-97-1; 1 mg per mouse loading dose followed by 300 μg per mouse maintenance doses, administered intraperitoneally twice weekly), anti-PD-1 antibody (Bio X Cell, BE0146; 200 μg per dose, intraperitoneally, three doses at 3-day intervals) or the combination according to the treatment schedule shown in Figure 4g. Anti-IGF1R antibody was administered at a loading dose of 1 mg per mouse followed by 300 μg per mouse twice weekly; anti-PD-1 antibody was administered at 200 μg per dose for three doses at 3-day intervals. Saline was used as the matched vehicle for both antibody treatments.

The MyC-CaP validation experiment used intracardiac MyC-CaP-LUC injection followed by the same four treatment groups.

### In vitro functional assays

#### 1 Mouse BM neutrophil isolation and stimulation

BM neutrophils were isolated from the femurs and tibias of 3-month-old male C57BL/6J mice using Histopaque 1077/1119 density-gradient centrifugation as previously described ^144^, with minor modifications. Briefly, BM cells were collected from the femurs and tibias and treated with ACK lysis buffer (Absin, abs9101) to remove red blood cells. Cells were resuspended in 1 ml ice-cold HBSS (Gibco, C14175500BT) supplemented with 0.2% FBS (ExCell Bio, FSP500). The cell suspension was layered onto a discontinuous density gradient consisting of 2 ml Histopaque 1077 (Sigma, 10771) and 3 ml Histopaque 1119 (Sigma, 11191) in a 15-ml tube. Samples were centrifuged at 1,000 × g for 20 min at 4 °C. Neutrophils at the interface between the Histopaque 1077 and Histopaque 1119 layers were collected and washed before subsequent experiments.

Purified BM neutrophils were treated with saline vehicle (Control) or recombinant IGF1 (100 ng/ml; MCE, HY-P70698A). Neutrophils treated with IGF1 for 24 h were used for molecular and functional assays and are referred to as induced TAN-like neutrophils (iTANs). For T cell co-culture experiments, BM neutrophils were pretreated with IGF1 for 12 h before co-culture.

Intracellular reactive oxygen species (ROS) were measured using H2DCFDA (1 µM; MCE, HY-D0940). Where indicated, picropodophyllin (PPP; MCE, HY-15494), kenpaullone (MCE, HY-12302), PD98059 (MCE, HY-12028), MK-2206 (MCE, HY-108232) and 10058-F4 (MCE, HY-12702) were used during IGF1 stimulation to inhibit IGF1R, KLF4-associated signaling, MAPK/ERK, AKT and c-MYC signaling, respectively. N-acetylcysteine (NAC) (MCE, HY-B0215; 10 µM) and L-NAME (MCE, HY-18729A; 50 µM) were used where indicated to suppress ROS-and nitric oxide-associated activity, respectively, during IGF1 stimulation.

#### 2 T cell and Treg co-culture assays

Spleens were collected from male C57BL/6J mice, and red blood cells were removed using ACK lysis buffer (Absin, abs9101). Total T cells were isolated using the EasySep Mouse T Cell Isolation Kit (STEMCELL Technologies, 19851). Where indicated, CD8 T cells were isolated using the EasySep Mouse CD8 T Cell Isolation Kit (STEMCELL Technologies, 19853).

For proliferation assays, T cells were labelled with 2.5 µM CFSE (BD Pharmingen, 565082) for 20 min at room temperature. Cells were cultured in RPMI 1640 medium and activated with anti-mouse CD3 (Invitrogen, 16-0032-82) and anti-mouse CD28 (Invitrogen, 16-0281-82). For neutrophil-T cell co-culture assays, BM neutrophils were pretreated with IGF1 (100 ng/ml) for 12 h to generate iTANs. T cells were then co-cultured with untreated neutrophils or iTANs at a 1:1 ratio for 48 h. Where indicated, an anti-IL-1β neutralising antibody (Selleck, A2142; 5 µg/ml), NAC or L-NAME was added during co-culture to assess the contribution of IL-1β, ROS or nitric oxide-associated activity, respectively.

After co-culture, cells were collected and processed for flow cytometry. Where intracellular staining was required, cells were fixed and permeabilized using the Fixation/Permeabilization Kit (BD Pharmingen, 554714). Fc receptors were blocked using purified rat anti-mouse CD16/CD32 antibody (BD Pharmingen, 553141), and cell viability was assessed using Fixable Viability Stain 510 (BD Pharmingen, 564406). Flow-cytometric analyses included CD3 (PerCP-Cy5.5 rat anti-mouse CD3 molecular complex, clone 17A2; BD Pharmingen, 560527), granzyme B (PE anti-human/mouse Granzyme B recombinant antibody; BioLegend, 372208), perforin (APC anti-mouse Perforin; BioLegend, 154304) and IFN-γ (Brilliant Violet 650 anti-mouse IFN-γ; BioLegend, 505832).

For Treg assays, cell proliferation was assessed by CFSE dilution, and Treg phenotype and function were evaluated by CD25 and FOXP3 expression. CFSE dilution was analysed using the Cell Proliferation Platform in FlowJo software.

#### 3 Tumor-cell migration, invasion and wound-closure assays

For wound-closure assays, TRAMP-C1 cells were cultured in the lower chamber of 6-well plates, and BM neutrophils were placed in 0.4-µm Transwell inserts (LABSELECT). TRAMP-C1 cells and neutrophils were co-cultured for 24 h before wound closure was assessed.

For Transwell migration and invasion assays, TRAMP-C1 cells were seeded into 8-µm Transwell inserts (LABSELECT) in 12-well plates, with BM neutrophils cultured in the lower chamber. Cells were co-cultured for 24 h. For migration assays, inserts were used without extracellular matrix coating. For invasion assays, inserts were coated with Matrigel Matrix (500 µg ml ¹; Corning, 356237). PPP was added where indicated to assess the contribution of IGF1R signalling.

#### 4 Quantitative real-time PCR

Total RNA was isolated using RNA-easy Isolation Reagent (Vazyme, R701-01) according to the manufacturer’s instructions. cDNA was synthesized using HiScript II Q RT SuperMix for qPCR (Vazyme, R223-01). Quantitative real-time PCR was performed in technical duplicate using ChamQ SYBR qPCR Master Mix (Vazyme, Q311-02) on a QuantStudio real-time PCR system (Applied Biosystems).

Relative gene expression was calculated using the 2^−ΔΔCt^ method and normalized to *GAPDH* (human samples) or *Gapdh* (mouse samples), as appropriate. Primer sequences used for qPCR are provided in Table S10.

#### 5 Chromatin immunoprecipitation and ChIP-qPCR

ChIP was performed using the ChIP-IT High Sensitivity Kit (Active Motif, 53040) according to the manufacturer’s instructions. ChIP-enriched DNA was analysed by quantitative PCR as described above. c-MYC occupancy at the Klf4 promoter was quantified as the percentage of input DNA, with normal IgG used as a negative control. Primer sequences used for ChIP-qPCR are provided in Table S10.

#### 6 Histology, immunohistochemistry and immunofluorescence

Formalin-fixed paraffin-embedded (FFPE) tissues were sectioned at 10 µm. Bone specimens were decalcified in 10% EDTA solution (pH 7.4) for 2–4 weeks at 4 °C, with regular replacement of the decalcification solution, before paraffin embedding. Tissue sections were deparaffinized in xylene and rehydrated through graded ethanol solutions.

Haematoxylin and eosin (H&E) staining was performed to identify metastatic lesions and quantify tumor area. Tumor area was quantified using ImageJ in regions located 0.2–2 mm below the growth plate.

For IGF1 immunohistochemistry, heat-mediated antigen retrieval was performed using sodium citrate buffer (10 mM sodium citrate, 0.05% Tween 20, pH 6.0). Sections were stained with anti-IGF1 antibody (1:200; Proteintech, 28530-1-AP), counterstained with haematoxylin and dehydrated through graded ethanol and xylene.

Multiplex immunofluorescence was performed using antibodies against the indicated human or mouse markers, including CCR3, CD66b, ICAM1/CD54, Ly6G, CD8, GZMB, CD4, FOXP3, CD3, ITGAL and CCL5. Antibodies used in the indicated experiments included anti-mouse CCR3 (1:200; Abcam, ab318266), anti-human CCR3 (1:200; Abcam, ab227032), anti-human CD54 (1:500; ABclonal, A26412PM), anti-human CD66b (1:400; BioLegend, 305102) and anti-mouse Ly6G (1:200; BioLegend, 127601). Where conventional indirect immunofluorescence staining was used, sections were incubated with appropriate fluorescently labelled secondary antibodies (1:200; Thermo Fisher Scientific) and nuclei were counterstained with DAPI. Complete information for antibodies used in immunostaining experiments is provided in Table S11. Images within each experiment were acquired using identical imaging settings.

#### 7 Immunofluorescence staining of cultured neutrophils

For immunofluorescence staining of cultured neutrophils, chamber slides (LABSELECT, CS-08-TC) were coated with fibronectin (500 µg ml ¹; Roche, 10838039001) for 40 min at 37 °C. Neutrophils were then seeded onto the coated slides and incubated for an additional 40 min at 37 °C.

Cells were fixed with 4% paraformaldehyde (Beyotime, P0099) for 30 min, permeabilized with Triton X-100 (Beyotime, P0096) for 30 min and blocked with 5% bovine serum albumin (BSA; Sigma, A1933) for 30 min. Cells were incubated with the indicated primary antibodies overnight at 4 °C, followed by fluorescently conjugated secondary antibodies for 45 min at room temperature. Nuclei were counterstained using an antifade mounting medium containing DAPI (Beyotime, P0131).

Primary antibodies included anti-CD274 (Proteintech, 66248-1-Ig), anti-Ly6G (Cell Signaling Technology, 31469S) and anti-CCR3 (Abcam, ab318266). Secondary antibodies included Alexa Fluor 488 goat anti-mouse IgG (Invitrogen, A11029), Alexa Fluor 555 goat anti-rat IgG (H+L) (Invitrogen, A48263) and Alexa Fluor 633 goat anti-rabbit IgG (H+L) (Invitrogen, A21071).

Images were acquired using a Nikon confocal microscope under identical settings within each experiment. Complete antibody information is provided in Table S11.

#### 8 Western blotting

BM neutrophils were isolated as described above and lysed in RIPA buffer supplemented with protease and phosphatase inhibitor cocktails. Cell lysates were incubated on ice for 30 min with intermittent vortexing and centrifuged at 12,000 × g for 15 min at 4 °C to remove insoluble debris. Protein concentrations were determined using a bicinchoninic acid (BCA) assay. Equal amounts of protein were separated by SDS–PAGE and transferred onto PVDF membranes. Membranes were blocked with 5% non-fat milk in TBST and incubated with the indicated primary antibodies overnight at 4 °C, followed by HRP-conjugated secondary antibodies. Protein signals were detected using an enhanced chemiluminescence (ECL) substrate and acquired with a chemiluminescence imaging system.

#### 9 Flow cytometry

Single-cell suspensions were prepared from mouse BM, spleen, PB and the indicated tissues, as well as from human PB. Samples were analysed using a BD LSRFortessa flow cytometer, and data were processed using FlowJo v10.8.

Erythrocytes were lysed before antibody staining. For human peripheral-blood samples, erythrocytes were lysed using RBC lysis buffer containing 155 mM NH4Cl, 12 mM NaHCO3 and 0.1 mM EDTA. Cells were washed twice with PBS and filtered through a 70-µm nylon mesh before staining. Live single cells were selected before subsequent lineage-specific gating.

Flow-cytometric panels included markers for neutrophils and TANs, including CD11b, Ly6G or CD66b, CCR3, ICAM1/CD54, PD-L1/CD274 and IGF1R; monocytes, including Ly6C and CD115; CD8 T cell activation or dysfunction, including CD8, GZMB, IFNγ, PD-1 and TIM-3; and regulatory T cells, including CD4 and FOXP3. Additional markers were included as indicated for individual experiments. Complete antibody information, including antibodies used in additional panels, is provided in Table S11.

Gating strategies for the indicated neutrophil, monocyte and T cell populations are shown in the Supplementary Figures.

#### 10 IGF1 and IGFBP3 measurements

IGF1 and IGFBP3 concentrations in human plasma and the indicated mouse samples, including PB, BM supernatant and liver extracts, as well as CM, were measured by ELISA according to the manufacturers’ instructions.

Human plasma IGF1 and IGFBP3 concentrations were measured using ELISA kits from Jianglaibio (IGF1, JLO11625; IGFBP3, JL10882). For collection of mouse BM supernatant, mice were euthanized and both ends of the tibiae were opened. Tibiae were centrifuged at 800 × g for 15 min at 4 °C, and the recovered BM supernatants were stored at −80 °C until analysis. Mouse BM IGF1 and IGFBP3 were measured using the corresponding ELISA kits (Jianglaibio, JL12620 and JL10973). [

The IGF1 bioavailability index was calculated as [IGF1 (ng/ml) × 0.13]/[IGFBP3 (ng/ml) × 0.036] ^145^.

#### 11 Statistical analysis

Unless otherwise specified, statistical tests were two-sided. Data are presented as mean ± s.d. or as otherwise indicated in the corresponding figure legends. Paired or unpaired Student’s t-tests, Wilcoxon rank-sum tests, Pearson or Spearman correlation analyses, log-rank tests, Gehan-Breslow tests and beta-binomial regression were used as appropriate.

Where multiple t-tests or related comparisons were performed within an analysis, P values were adjusted for multiple comparisons using the Benjamini-Hochberg method. The patient, mouse or independent cell preparation was treated as the biological replicate. For single-cell abundance and signature-score analyses, values were summarized at the patient or sample level wherever biological replication permitted.

Exact sample sizes, definitions of centre and dispersion, and the statistical tests used for individual experiments are provided in the corresponding figure legends. P < 0.05 was considered statistically significant. Statistical analyses were performed using R v4.2.2. (2019).

## Acknowledgments

We thank Lai Guan Ng for his insightful discussion. We thank NovelBio Co., Ltd. for assistance with single-cell RNA sequencing. We thank Innovent Biologics, Inc. for providing teprotumumab and BeiGene, Ltd. for providing tislelizumab for this study.

## Funding

National Key Research and Development Program of China (2024ZD0525200)

National Natural Science Foundation of China (82125025)

National Natural Science Foundation of China (82330091)

Shanghai Municipal Education Commission (2023ZKZD46)

Noncommunicable Chronic Diseases-National Science and Technology Major Project (2023ZD0505600, 2023ZD0505601)

National Natural Science Foundation of China (82373106)

National Natural Science Foundation of China (92474108)

National Natural Science Foundation of Sichuan (2024NSFSC0758)

National Natural Science Foundation of China (82372987)

Shanghai Rising-Star Program (23QA1408100)

Natural Science Foundation of Shanghai (23ZR1451800)

Shanghai Municipal Health Commission (2022YQ065)

National Natural Science Foundation of China (32200532)

## Author contributions

S.R., S.C., G.Z. and Z.Y. designed the study and supervised the experimental framework. Y.T., X.Z., Y.L. and D.L. performed data analysis and statistical modeling under the supervision of S.C. and G.Z. X.Z., Y.L., X.W., J.W. and D.Z. conducted experiments and validated the results. Y.Z., H.H., Z.L. and B.Q. contributed to methodology development and technical implementation. S.R. and J.X. provided clinical samples and related resources. S.R., S.C., G.Z. and Z.Y. critically reviewed the experimental design, coordinated quality control, and oversaw project conceptualization and funding acquisition. Y.T., X.Z., Y.L., D.L. and S.C. drafted the manuscript. All authors participated in revising the manuscript and approved the final version.

## Competing interests

The authors declare no competing interests.

## Data and materials availability

Raw sequencing data have been uploaded to the GSA- Human database under accession code PRJCA043924 (https://ngdc.cncb.ac.cn/bioproject/browse/PRJCA043924). Our analysis code has been uploaded to github (https://github.com/iceautumn/PCa_BM). Accession numbers and source information for all public datasets are listed in Table S9. Any additional information required to reanalyze the data reported in this paper is available from the lead contact upon request.

## Supplementary Materials

Figs. S1 to S15

Tables S1 to S11

## Notes

### Competing Interest Statement

The authors have declared no competing interest.

## References

1 Sung, H. et al. Global Cancer Statistics 2020: GLOBOCAN Estimates of Incidence and Mortality Worldwide for 36 Cancers in 185 Countries. 71, 209–249, doi: 10.3322/caac.21660 (2021).

2 Ferlay, J. et al. Cancer statistics for the year 2020: An overview. 149, 778–789, doi: 10.1002/ijc.33588 (2021).

3 Siegel, R. L., Kratzer, T. B., Giaquinto, A. N., Sung, H. & Jemal, A. Cancer statistics, 2025. CA: A Cancer Journal for Clinicians 75, 10–45, doi: 10.3322/caac.21871 (2025).

4 Bubendorf, L. et al. Metastatic patterns of prostate cancer: An autopsy study of 1,589 patients. Human Pathology 31, 578–583, doi: 10.1053/hp.2000.6698 (2000).

5 Liu, D. et al. Prognosis of prostate cancer and bone metastasis pattern of patients: a SEER-based study and a local hospital based study from China. Scientific Reports 10, 9104, doi:10.1038/s41598-020-64073-6 (2020).

6 Nieder, C., Haukland, E., Pawinski, A. & Dalhaug, A. Pathologic fracture and metastatic spinal cord compression in patients with prostate cancer and bone metastases. BMC Urology 10, 23, doi:10.1186/1471-2490-10-23 (2010).

7 Ribelli, G. et al. Osteoblasts Promote Prostate Cancer Cell Proliferation Through Androgen Receptor Independent Mechanisms. 11, doi:10.3389/fonc.2021.789885 (2021).

8 Kirschenbaum, A., Liu, X.-H., Yao, S., Leiter, A. & Levine, A. C. Prostatic acid phosphatase is expressed in human prostate cancer bone metastases and promotes osteoblast differentiation. 1237, 64–70, doi:10.1111/j.1749-6632.2011.06198.x (2011).

9 Chen, G. et al. Expression of RANKL/RANK/OPG in primary and metastatic human prostate cancer as markers of disease stage and functional regulation. 107, 289–298, doi:10.1002/cncr.21978 (2006).

10 Corey, E. et al. Osteoprotegerin in Prostate Cancer Bone Metastasis. Cancer Research 65, 1710–1718, doi:10.1158/0008-5472.CAN-04-2033%J Cancer Research (2005).

11 Udagawa, N. et al. Osteoprotegerin Produced by Osteoblasts Is an Important Regulator in Osteoclast Development and Function*. Endocrinology 141, 3478–3484, doi:10.1210/endo.141.9.7634%J Endocrinology (2000).

12 Wang, S. et al. A pan-cancer single-cell transcriptomic atlas of human bone metastases. Cell Reports Medicine 7, doi:10.1016/j.xcrm.2025.102583 (2026).

13 Costanzo-Garvey, D. L. et al. Neutrophils are mediators of metastatic prostate cancer progression in bone. Cancer Immunology, Immunotherapy 69, 1113–1130, doi:10.1007/s00262-020-02527-6 (2020).

14 Alsamraae, M. et al. Androgen receptor inhibition suppresses anti-tumor neutrophil response against bone metastatic prostate cancer via regulation of TβRI expression. Cancer Letters 579, 216468, doi:10.1016/j.canlet.2023.216468 (2023).

15 Kfoury, Y. et al. Human prostate cancer bone metastases have an actionable immunosuppressive microenvironment. Cancer Cell 39, 1464–1478.e1468, doi:10.1016/j.ccell.2021.09.005 (2021).

16 Liu, Y. & Cao, X. Characteristics and Significance of the Pre-metastatic Niche. Cancer Cell 30, 668–681, doi:10.1016/j.ccell.2016.09.011 (2016).

17 Wang, Y. et al. Pre-metastatic niche: formation, characteristics and therapeutic implication. Signal Transduction and Targeted Therapy 9, 236, doi:10.1038/s41392-024-01937-7 (2024).

18 Monteiro, A. C. et al. Cooperation between T and B cells reinforce the establishment of bone metastases in a mouse model of breast cancer. Bone 178, 116932, doi:10.1016/j.bone.2023.116932 (2024).

19 Yuan, X. et al. Breast cancer exosomes contribute to pre-metastatic niche formation and promote bone metastasis of tumor cells. Theranostics 11, 1429–1445, doi:10.7150/thno.45351 (2021).

20 Ravi, P. et al. Long-term Outcomes and Prognostic Impact of Residual Cancer Burden After Intensified Neoadjuvant Therapy in High-risk Prostate Cancer. European Urology 87, 643–650, doi:10.1016/j.eururo.2025.01.015 (2025).

21 Vermeulen, A. H., Vermeer, C. & Bosman, F. T. Histochemical detection of osteocalcin in normal and pathological human bone. Journal of Histochemistry & Cytochemistry 37, 1503–1508, doi:10.1177/37.10.2789247 (1989).

22 Roudier, M. P. et al. Histopathological assessment of prostate cancer bone osteoblastic metastases. J Urol 180, 1154–1160, doi:10.1016/j.juro.2008.04.140 (2008).

23 Roudier, M. P. et al. in The Biology of Skeletal Metastases (eds Evan T. Keller & Leland W. K. Chung) 311–339 (Springer US, 2004).

24 Gorski, J. P. Is All Bone the Same? Distinctive Distributions and Properties of Non-Collagenous Matrix Proteins in Lamellar Vs. Woven Bone Imply the Existence of Different Underlying Osteogenic Mechanisms. Critical Reviews in Oral Biology & Medicine 9, 201–223, doi:10.1177/10454411980090020401 (1998).

25 Kikuchi, Y. et al. The Niche Component Periostin Is Produced by Cancer-Associated Fibroblasts, Supporting Growth of Gastric Cancer through ERK Activation. The American Journal of Pathology 184, 859–870, doi:10.1016/j.ajpath.2013.11.012 (2014).

26 Sasaki, Y. et al. Expression of asporin reprograms cancer cells to acquire resistance to oxidative stress. Cancer Science 112, 1251–1261, doi:10.1111/cas.14794 (2021).

27 Chiavarina, B. et al. Metastatic colorectal cancer cells maintain the TGFβ program and use TGFBI to fuel angiogenesis. Theranostics 11, 1626–1640, doi:10.7150/thno.51507 (2021).

28 Yang, H. et al. Multi-scale integrative analyses identify THBS2(+) cancer-associated fibroblasts as a key orchestrator promoting aggressiveness in early-stage lung adenocarcinoma. Theranostics 12, 3104–3130, doi:10.7150/thno.69590 (2022).

29 Abida, W. et al. Genomic correlates of clinical outcome in advanced prostate cancer. Proceedings of the National Academy of Sciences 116, 11428–11436, doi:10.1073/pnas.1902651116 (2019).

30 Chen, S. et al. Single-cell analysis reveals transcriptomic remodellings in distinct cell types that contribute to human prostate cancer progression. Nature Cell Biology 23, 87–98, doi:10.1038/s41556-020-00613-6 (2021).

31 Rocha, S. F. et al. Esm1 Modulates Endothelial Tip Cell Behavior and Vascular Permeability by Enhancing VEGF Bioavailability. Circulation Research 115, 581–590, doi:10.1161/CIRCRESAHA.115.304718 (2014).

32 Zhang, H. et al. Targeting Endothelial Cell-Specific Molecule 1 Protein in Cancer: A Promising Therapeutic Approach. Frontiers in Oncology 11 (2021).

33 Walker, A. M. N. et al. Endothelial Insulin Receptors Promote VEGF-A Signaling via ERK1/2 and Sprouting Angiogenesis. Endocrinology 162, bqab104, doi:10.1210/endocr/bqab104 (2021).

34 Salcedo, R. et al. Angiogenic effects of prostaglandin E2 are mediated by up-regulation of CXCR4 on human microvascular endothelial cells. Blood 102, 1966–1977, doi:10.1182/blood-2002-11-3400 (2003).

35 Uutela, M. et al. PDGF-D induces macrophage recruitment, increased interstitial pressure, and blood vessel maturation during angiogenesis. Blood 104, 3198–3204, doi:10.1182/blood-2004-04-1485 (2004).

36 Gordon, S. & Plüddemann, A. Tissue macrophages: heterogeneity and functions. BMC Biology 15, 53, doi:10.1186/s12915-017-0392-4 (2017).

37 Jones, R. C. et al. The Tabula Sapiens: A multiple-organ, single-cell transcriptomic atlas of humans. Science 376, eabl4896, doi:10.1126/science.abl4896.

38 Zhang, H. et al. Single-Cell RNA Sequencing Reveals B Cells Are Important Regulators in Fracture Healing. Frontiers in Endocrinology Volume 12 **-** 2021 (2021).

39 Bandyopadhyay, S. et al. Mapping the cellular biogeography of human bone marrow niches using single-cell transcriptomics and proteomic imaging. Cell 187, 3120–3140.e3129, doi:10.1016/j.cell.2024.04.013 (2024).

40 Bancaro, N. et al. Apolipoprotein E induces pathogenic senescent-like myeloid cells in prostate cancer. Cancer Cell 41, 602–619.e611, doi:10.1016/j.ccell.2023.02.004 (2023).

41 Calì, B. et al. Coagulation factor X promotes resistance to androgen-deprivation therapy in prostate cancer. Cancer Cell 42, 1676–1692.e1611, doi:10.1016/j.ccell.2024.08.018 (2024).

42 Payne, K. K. et al. BTN3A1 governs antitumor responses by coordinating αβ and γδ T cells. 369, 942–949, doi:doi:10.1126/science.aay2767 (2020).

43 Smith, I. A. et al. BTN1A1, the Mammary Gland Butyrophilin, and BTN2A2 Are Both Inhibitors of T Cell Activation. The Journal of Immunology 184, 3514–3525, doi:10.4049/jimmunol.0900416 %J The Journal of Immunology (2010).

44 Huang, Y.-C. et al. Single-Cell Transcriptomic Profiles of Lung Pre-Metastatic Niche Reveal Neutrophil and Lymphatic Endothelial Cell Roles in Breast Cancer. 15, 176 (2023).

45 Liu, C.-Y. et al. Population alterations of l-arginase- and inducible nitric oxide synthase- expressed CD11b+/CD14−/CD15+/CD33+ myeloid-derived suppressor cells and CD8+ T lymphocytes in patients with advanced-stage non-small cell lung cancer. Journal of Cancer Research and Clinical Oncology 136, 35–45, doi:10.1007/s00432-009-0634-0 (2010).

46 Cemerski, S., Cantagrel, A., van Meerwijk, J. P. M. & Romagnoli, P. Reactive Oxygen Species Differentially Affect T Cell Receptor-signaling Pathways*. Journal of Biological Chemistry 277, 19585–19593, doi:10.1074/jbc.M111451200 (2002).

47 Yarosz, E. L. & Chang, C.-H. The Role of Reactive Oxygen Species in Regulating T Cell-mediated Immunity and Disease. Immune Netw 18 (2018).

48 Godoy-Tundidor, S. et al. Interleukin-6 and oncostatin M stimulation of proliferation of prostate cancer 22Rv1 cells through the signaling pathways of p38 mitogen-activated protein kinase and phosphatidylinositol 3-kinase. 64, 209–216, doi:10.1002/pros.20235 (2005).

49 Smith, D. A., Kiba, A., Zong, Y. & Witte, O. N. Interleukin-6 and Oncostatin-M Synergize with the PI3K/AKT Pathway to Promote Aggressive Prostate Malignancy in Mouse and Human Tissues. Molecular Cancer Research 11, 1159–1165, doi:10.1158/1541-7786.MCR-13-0238 %J Molecular Cancer Research (2013).

50 Oetjen, K. A. et al. Human bone marrow assessment by single-cell RNA sequencing, mass cytometry, and flow cytometry. JCI Insight 3, doi:10.1172/jci.insight.124928 (2018).

51 Hajaj, E. et al. SLAMF6 deficiency augments tumor killing and skews toward an effector phenotype revealing it as a novel T cell checkpoint. eLife 9, e52539, doi:10.7554/eLife.52539 (2020).

52 Kared, H. et al. SLAMF7 defines subsets of human effector CD8 T cells. Scientific Reports 14, 30779, doi:10.1038/s41598-024-80971-5 (2024).

53 Reading, S. M. et al. Abstract PR01: Regulation of exhausted CD8+ T cell differentiation by IKZF transcription factors. Cancer Immunology Research 11, PR01–PR01, doi:10.1158/2326-6074.TUMIMM23-PR01 (2023).

54 Zou, Y. et al. IKZF3 deficiency potentiates chimeric antigen receptor T cells targeting solid tumors. Cancer Letters 524, 121–130, doi:10.1016/j.canlet.2021.10.016 (2022).

55 Zhang, W. et al. ICAM-1-mediated adhesion is a prerequisite for exosome-induced T-cell suppression. Developmental Cell 57, 329–343.e327, doi:10.1016/j.devcel.2022.01.002 (2022).

56 Jakubzick, C. V., Randolph, G. J. & Henson, P. M. Monocyte differentiation and antigen-presenting functions. Nature Reviews Immunology 17, 349–362, doi:10.1038/nri.2017.28 (2017).

57 Dunsmore, G. et al. Timing and location dictate monocyte fate and their transition to tumor-associated macrophages. Science Immunology 9, eadk3981, doi:10.1126/sciimmunol.adk3981 (2024).

58 Giral, H. et al. Rapid Inflammasome Activation Is Attenuated in Post-Myocardial Infarction Monocytes. Frontiers in Immunology 13 (2022).

59 Thomas, G., Tacke, R., Hedrick, C. C. & Hanna, R. N. Nonclassical Patrolling Monocyte Function in the Vasculature. *Arteriosclerosis*, Thrombosis, and Vascular Biology 35, 1306–1316, doi:10.1161/ATVBAHA.114.304650 (2015).

60 Hanna, R. N. et al. The transcription factor NR4A1 (Nur77) controls bone marrow differentiation and the survival of Ly6C− monocytes. Nature Immunology 12, 778–785, doi:10.1038/ni.2063 (2011).

61 Mildner, A. et al. Genomic Characterization of Murine Monocytes Reveals C/EBPβ Transcription Factor Dependence of Ly6C− Cells. Immunity 46, 849–862.e847, doi:10.1016/j.immuni.2017.04.018 (2017).

62 Balasubramanian, S., Kota, S. K., Kuchroo, V. K., Humphreys, B. D. & Strom, T. B. TIM Family Proteins Promote the Lysosomal Degradation of the Nuclear Receptor NUR77. Science Signaling 5, ra90–ra90, doi:10.1126/scisignal.2003200 (2012).

63 Lee, S. O. et al. The orphan nuclear receptor NR4A1 (Nur77) regulates oxidative and endoplasmic reticulum stress in pancreatic cancer cells. Molecular cancer research : MCR 12, 527–538, doi:10.1158/1541-7786.Mcr-13-0567 (2014).

64 Kim, K. S. et al. TIM-3 on myeloid cells promotes pulmonary inflammation through increased production of galectin-3. Communications biology 7, 1090, doi:10.1038/s42003-024-06762-w (2024).

65 Chu, Y. et al. Pan-cancer T cell atlas links a cellular stress response state to immunotherapy resistance. Nature Medicine 29, 1550–1562, doi:10.1038/s41591-023-02371-y (2023).

66 Ross, S. H., Rollings, C. M. & Cantrell, D. A. Quantitative Analyses Reveal How Hypoxia Reconfigures the Proteome of Primary Cytotoxic T Lymphocytes. 12, doi:10.3389/fimmu.2021.712402 (2021).

67 Scharping, N. E. et al. Mitochondrial stress induced by continuous stimulation under hypoxia rapidly drives T cell exhaustion. Nature Immunology 22, 205–215, doi:10.1038/s41590-020-00834-9 (2021).

68 Cheng, J. et al. ETV7 limits the antiviral and antitumor efficacy of CD8+ T cells by diverting their fate toward exhaustion. Nature Cancer 6, 338–356, doi:10.1038/s43018-024-00892-0 (2025).

69 Liu, X. et al. Regulatory T cells trigger effector T cell DNA damage and senescence caused by metabolic competition. Nature Communications 9, 249, doi:10.1038/s41467-017-02689-5 (2018).

70 Ivashkiv, L. B. The hypoxia–lactate axis tempers inflammation. Nature Reviews Immunology 20, 85–86, doi:10.1038/s41577-019-0259-8 (2020).

71 Staron, Matthew M. et al. The Transcription Factor FoxO1 Sustains Expression of the Inhibitory Receptor PD-1 and Survival of Antiviral CD8^+^ T Cells during Chronic Infection. Immunity 41, 802–814, doi:10.1016/j.immuni.2014.10.013 (2014).

72 Brinster, C. & Shevach, E. M. Costimulatory effects of IL-1 on the expansion/differentiation of CD4+CD25+Foxp3+ and CD4+CD25+Foxp3– T cells. Journal of Leukocyte Biology 84, 480–487, doi:10.1189/jlb.0208085 %J Journal of Leukocyte Biology (2008).

73 Monteran, L. et al. Combining TIGIT Blockade with MDSC Inhibition Hinders Breast Cancer Bone Metastasis by Activating Antitumor Immunity. Cancer Discovery 14, 1252–1275, doi:10.1158/2159-8290.CD-23-0762 %J Cancer Discovery (2024).

74 Guo, J. et al. KMT2C deficiency drives transdifferentiation of double-negative prostate cancer and confer resistance to AR-targeted therapy. Cancer Cell 43, 1261–1278.e1210, doi:10.1016/j.ccell.2025.04.002 (2025).

75 Luo, H. et al. Pan-cancer single-cell analysis reveals the heterogeneity and plasticity of cancer-associated fibroblasts in the tumor microenvironment. Nat Commun 13, 6619, doi:10.1038/s41467-022-34395-2 (2022).

76 Ohlsson, C. et al. The Role of Liver-Derived Insulin-Like Growth Factor-I. Endocrine Reviews 30, 494–535, doi:10.1210/er.2009-0010 (2009).

77 van Golen, C. M. et al. Insulin-Like Growth Factor-I Receptor Expression Regulates Neuroblastoma Metastasis to Bone. Cancer Research 66, 6570–6578, doi:10.1158/0008-5472.CAN-05-1448 (2006).

78 Hiraga, T. et al. Bone-Derived IGF Mediates Crosstalk between Bone and Breast Cancer Cells in Bony Metastases. Cancer Research 72, 4238–4249, doi:10.1158/0008-5472.CAN-11-3061 (2012).

79 Wang, J. et al. Bone marrow-derived IGF-1 orchestrates maintenance and regeneration of the adult skeleton. Proceedings of the National Academy of Sciences 120, e2203779120, doi:10.1073/pnas.2203779120 (2023).

80 Seck, T. et al. Concentration of Insulin-Like Growth Factor (IGF)-I and -II in Iliac Crest Bone Matrix from Pre- and Postmenopausal Women: Relationship to Age, Menopause, Bone Turnover, Bone Volume, and Circulating IGFs1. The Journal of Clinical Endocrinology & Metabolism 83, 2331–2337, doi:10.1210/jcem.83.7.4967 (1998).

81 Abeshouse, A. et al. The Molecular Taxonomy of Primary Prostate Cancer. Cell 163, 1011–1025, doi:10.1016/j.cell.2015.10.025 (2015).

82 Joanito, I. et al. Single-cell and bulk transcriptome sequencing identifies two epithelial tumor cell states and refines the consensus molecular classification of colorectal cancer. Nature Genetics 54, 963–975, doi:10.1038/s41588-022-01100-4 (2022).

83 Bhatlekar, S., Fields, J. Z. & Boman, B. M. Role of HOX Genes in Stem Cell Differentiation and Cancer. Stem Cells International 2018, 3569493, doi:10.1155/2018/3569493 (2018).

84 Wang, X. et al. A luminal epithelial stem cell that is a cell of origin for prostate cancer. Nature 461, 495–500, doi:10.1038/nature08361 (2009).

85 German, B. et al. MYBL2 Drives Prostate Cancer Plasticity: Inhibiting Its Transcriptional Target CDK2 for RB1-Deficient Neuroendocrine Prostate Cancer. Cancer Research Communications 4, 2295–2307, doi:10.1158/2767-9764.CRC-24-0069 (2024).

86 Paltoglou, S. et al. Novel Androgen Receptor Coregulator GRHL2 Exerts Both Oncogenic and Antimetastatic Functions in Prostate Cancer. Cancer Research 77, 3417–3430, doi:10.1158/0008-5472.CAN-16-1616 (2017).

87 Prencipe, M. et al. Role of serum response factor expression in prostate cancer biochemical recurrence. The Prostate 78, 724–730, doi:10.1002/pros.23516 (2018).

88 Li, L. et al. EGR1 regulates angiogenic and osteoclastogenic factors in prostate cancer and promotes metastasis. Oncogene 38, 6241–6255, doi:10.1038/s41388-019-0873-8 (2019).

89 Vidal, Samuel J. et al. A Targetable GATA2-IGF2 Axis Confers Aggressiveness in Lethal Prostate Cancer. Cancer Cell 27, 223–239, doi:10.1016/j.ccell.2014.11.013 (2015).

90 Chen, S. et al. Widespread and Functional RNA Circularization in Localized Prostate Cancer. Cell 176, 831–843.e822, doi:10.1016/j.cell.2019.01.025 (2019).

91 Tang, D. G. et al. Prostate cancer stem/progenitor cells: Identification, characterization, and implications. 46, 1–14, doi:10.1002/mc.20255 (2007).

92 Kirk, J. S. et al. Integrated single-cell analysis defines the epigenetic basis of castration-resistant prostate luminal cells. Cell Stem Cell 31, 1203–1221.e1207, doi:10.1016/j.stem.2024.05.008 (2024).

93 Gavish, A. et al. Hallmarks of transcriptional intratumour heterogeneity across a thousand tumours. Nature 618, 598–606, doi:10.1038/s41586-023-06130-4 (2023).

94 Liu, Y. Fatty acid oxidation is a dominant bioenergetic pathway in prostate cancer. Prostate Cancer and Prostatic Diseases 9, 230–234, doi:10.1038/sj.pcan.4500879 (2006).

95 Cohen, R. J., Holland, J. W., Redmond, S. L., McNeal, J. E. & Dawkins, H. J. S. Identification of the glycosaminoglycan keratan sulfate in the prostatic secretory cell. The Prostate 44, 204–209, doi:10.1002/1097-0045(20000801)44:3<204::AID-PROS4>3.0.CO;2-W (2000).

96 Holland, J. W., Meehan, K. L., Redmond, S. L. & Dawkins, H. J. S. Purification of the keratan sulfate proteoglycan expressed in prostatic secretory cells and its identification as lumican. The Prostate 59, 252–259, doi:10.1002/pros.20002 (2004).

97 Ren, S. et al. Whole-genome and Transcriptome Sequencing of Prostate Cancer Identify New Genetic Alterations Driving Disease Progression. European Urology 73, 322–339, doi:10.1016/j.eururo.2017.08.027 (2018).

98 Guntur, A. R. & Rosen, C. J. IGF-1 regulation of key signaling pathways in bone. BoneKEy reports 2, 437, doi:10.1038/bonekey.2013.171 (2013).

99 Hazzalin, C. A. & Mahadevan, L. C. MAPK-regulated transcription: a continuously variable gene switch? Nature Reviews Molecular Cell Biology 3, 30–40, doi:10.1038/nrm715 (2002).

100 van Gastel, N. et al. Lipid availability determines fate of skeletal progenitor cells via SOX9. Nature 579, 111–117, doi:10.1038/s41586-020-2050-1 (2020).

101 Stumpf, P. S. et al. Transfer learning efficiently maps bone marrow cell types from mouse to human using single-cell RNA sequencing. Communications biology 3, 736, doi:10.1038/s42003-020-01463-6 (2020).

102 Antonarakis, E. S. et al. Pembrolizumab for Treatment-Refractory Metastatic Castration-Resistant Prostate Cancer: Multicohort, Open-Label Phase II KEYNOTE-199 Study. Journal of Clinical Oncology 38, 395–405, doi:10.1200/JCO.19.01638 (2019).

103 Powles, T. et al. Atezolizumab with enzalutamide versus enzalutamide alone in metastatic castration-resistant prostate cancer: a randomized phase 3 trial. Nature Medicine 28, 144–153, doi:10.1038/s41591-021-01600-6 (2022).

104 Sartor, O. et al. An exploratory analysis of alkaline phosphatase, lactate dehydrogenase, and prostate-specific antigen dynamics in the phase 3 ALSYMPCA trial with radium-223. Annals of oncology : official journal of the European Society for Medical Oncology 28, 1090–1097, doi:10.1093/annonc/mdx044 (2017).

105 Romero-Laorden, N. et al. Prospective Assessment of Bone Metabolism Biomarkers and Survival in Metastatic Castration-resistant Prostate Cancer Patients Treated with Radium-223: The PRORADIUM Study. European urology oncology 7, 447–455, doi:10.1016/j.euo.2023.09.015 (2024).

106 Graf, R. P. et al. Comparative Effectiveness of Immune Checkpoint Inhibitors vs Chemotherapy by Tumor Mutational Burden in Metastatic Castration-Resistant Prostate Cancer. JAMA Network Open 5, e225394–e225394, doi:10.1001/jamanetworkopen.2022.5394 (2022).

107 Hotte, S. J. et al. LBA51 - CCTG IND 232: A phase II study of durvalumab with or without tremelimumab in patients with metastatic castration resistant prostate cancer (mCRPC). Annals of Oncology 30, v885, doi:10.1093/annonc/mdz394.044 (2019).

108 Graff, J. N. et al. Pembrolizumab plus enzalutamide versus placebo plus enzalutamide for chemotherapy-naive metastatic castration-resistant prostate cancer: the randomized, double-blind, phase III KEYNOTE-641 study. Annals of Oncology 36, 976–987, doi:10.1016/j.annonc.2025.05.007 (2025).

109 Chen, S. Ultrafast one-pass FASTQ data preprocessing, quality control, and deduplication using fastp. iMeta 2, e107, doi:10.1002/imt2.107 (2023).

110 Smith, T., Heger, A. & Sudbery, I. UMI-tools: modeling sequencing errors in Unique Molecular Identifiers to improve quantification accuracy. Genome research 27, 491–499 (2017).

111 Dobin, A. et al. STAR: ultrafast universal RNA-seq aligner. Bioinformatics 29, 15–21, doi:10.1093/bioinformatics/bts635 (2013).

112 Karlsson, M. et al. A single–cell type transcriptomics map of human tissues. Science Advances 7, eabh2169, doi:10.1126/sciadv.abh2169.

113 Browaeys, R., Saelens, W. & Saeys, Y. NicheNet: modeling intercellular communication by linking ligands to target genes. Nature Methods 17, 159–162, doi:10.1038/s41592-019-0667-5 (2020).

114 Trapnell, C. et al. The dynamics and regulators of cell fate decisions are revealed by pseudotemporal ordering of single cells. Nature Biotechnology 32, 381–386, doi:10.1038/nbt.2859 (2014).

115 Angerer, P. et al. destiny: diffusion maps for large-scale single-cell data in R. Bioinformatics 32, 1241–1243, doi:10.1093/bioinformatics/btv715 (2016).

116 Kang, M. et al. Mapping single-cell developmental potential in health and disease with interpretable deep learning. bioRxiv, 2024.2003.2019.585637, doi:10.1101/2024.03.19.585637 (2024).

117 Hsiao, L.-L. et al. A compendium of gene expression in normal human tissues. Physiological Genomics 7, 97–104, doi:10.1152/physiolgenomics.00040.2001 (2001).

118 Zheng, L. et al. Pan-cancer single-cell landscape of tumor-infiltrating T cells. *Science (New York*, N.Y.) 374, abe6474, doi:10.1126/science.abe6474 (2021).

119 Hu, J. et al. Spatially resolved transcriptomic analysis of the adult human prostate. Nature Genetics 57, 922–933, doi:10.1038/s41588-025-02139-9 (2025).

120 Lima, T. et al. Gal-3 Protein Expression and Localization in Prostate Tumours. Current Oncology 30, 2729–2742 (2023).

121 Whitaker, H. C., Warren, A. Y., Eeles, R., Kote-Jarai, Z. & Neal, D. E. The potential value of microseminoprotein-β as a prostate cancer biomarker and therapeutic target. The Prostate 70, 333–340, doi:10.1002/pros.21059 (2010).

122 Ye, B., Ding, K., Li, K. & Zhu, Q. Study on the role of SLC14A1 gene in biochemical recurrence of prostate cancer. Scientific Reports 12, 17064, doi:10.1038/s41598-022-20775-7 (2022).

123 Kluz, P. N. et al. Cancer cell-intrinsic function of CD177 in attenuating β-catenin signaling. Oncogene 39, 2877–2889, doi:10.1038/s41388-020-1203-x (2020).

124 Wagner, A. et al. Metabolic modeling of single Th17 cells reveals regulators of autoimmunity. Cell 184, 4168–4185.e4121, doi:10.1016/j.cell.2021.05.045 (2021).

125 Thiele, I. et al. A community-driven global reconstruction of human metabolism. Nature Biotechnology 31, 419–425, doi:10.1038/nbt.2488 (2013).

126 Sun, K. et al. scRNA-seq of gastric tumor shows complex intercellular interaction with an alternative T cell exhaustion trajectory. Nature Communications 13, 4943, doi:10.1038/s41467-022-32627-z (2022).

127 Efremova, M., Vento-Tormo, M., Teichmann, S. A. & Vento-Tormo, R. CellPhoneDB: inferring cell–cell communication from combined expression of multi-subunit ligand– receptor complexes. Nature Protocols 15, 1484–1506, doi:10.1038/s41596-020-0292-x (2020).

128 Wolf, F. A., Angerer, P. & Theis, F. J. SCANPY: large-scale single-cell gene expression data analysis. Genome Biology 19, 15, doi:10.1186/s13059-017-1382-0 (2018).

129 Ushey, K., Allaire, J. & Tang, Y. reticulate: Interface to ‘Python’. (2024).

130 Bergen, V., Lange, M., Peidli, S., Wolf, F. A. & Theis, F. J. Generalizing RNA velocity to transient cell states through dynamical modeling. Nature Biotechnology 38, 1408–1414, doi:10.1038/s41587-020-0591-3 (2020).

131 O’Flanagan, C. H. et al. Dissociation of solid tumor tissues with cold active protease for single-cell RNA-seq minimizes conserved collagenase-associated stress responses. Genome Biology 20, 210, doi:10.1186/s13059-019-1830-0 (2019).

132 Liao, Y., Smyth, G. K. & Shi, W. featureCounts: an efficient general purpose program for assigning sequence reads to genomic features. Bioinformatics 30, 923–930, doi:10.1093/bioinformatics/btt656 (2014).

133 Love, M. I., Huber, W. & Anders, S. Moderated estimation of fold change and dispersion for RNA-seq data with DESeq2. Genome Biology 15, 550, doi:10.1186/s13059-014-0550-8 (2014).

134 Vasimuddin, M., Misra, S., Li, H. & Aluru, S. in 2019 IEEE International Parallel and Distributed Processing Symposium (IPDPS). 314–324.

135 Danecek, P. et al. Twelve years of SAMtools and BCFtools. GigaScience 10, giab008, doi:10.1093/gigascience/giab008 (2021).

136 Zhang, Y. et al. Model-based Analysis of ChIP-Seq (MACS). Genome Biology 9, R137, doi:10.1186/gb-2008-9-9-r137 (2008).

137 Stark R, B. G. DiffBind: differential binding analysis of ChIP-Seq peak data. doi:10.18129/B9.bioc.DiffBind (2011).

138 Heinz, S. et al. Simple Combinations of Lineage-Determining Transcription Factors Prime cis-Regulatory Elements Required for Macrophage and B Cell Identities. Molecular Cell 38, 576–589, doi:10.1016/j.molcel.2010.05.004 (2010).

139 Rauluseviciute, I. et al. JASPAR 2024: 20th anniversary of the open-access database of transcription factor binding profiles. Nucleic Acids Research 52, D174–D182, doi:10.1093/nar/gkad1059 (2024).

140 Kaplan, R. N. et al. VEGFR1-positive haematopoietic bone marrow progenitors initiate the pre-metastatic niche. Nature 438, 820–827, doi:10.1038/nature04186 (2005).

141 Yue, Z. et al. RSPO2 and RANKL signal through LGR4 to regulate osteoclastic premetastatic niche formation and bone metastasis. The Journal of clinical investigation 132, doi:10.1172/JCI144579 (2022).

142 Duan, F. et al. Area under the curve as a tool to measure kinetics of tumor growth in experimental animals. Journal of Immunological Methods 382, 224–228, doi:10.1016/j.jim.2012.06.005 (2012).

143 Li, R. et al. Therapeutically reprogrammed nutrient signalling enhances nanoparticulate albumin bound drug uptake and efficacy in KRAS-mutant cancer. Nature Nanotechnology 16, 830–839, doi:10.1038/s41565-021-00897-1 (2021).

144 Ouyang, S. et al. Targeting IL-17A/glucocorticoid synergy to CSF3 expression in neutrophilic airway diseases. JCI Insight 5, doi:10.1172/jci.insight.132836 (2020).

145 Hang, D. et al. Plasma Biomarkers of Insulin and the Insulin-like Growth Factor Axis, and Risk of Colorectal Adenoma and Serrated Polyp. JNCI Cancer Spectrum 3, pkz056, doi:10.1093/jncics/pkz056

