## Supplemental Figures 1-15 for "Neutrophils orchestrate pre-metastatic immunosuppressive bone marrow remodeling during prostate cancer metastasis"

**The PDF file includes:**

Figs. S1 to S15

**Other Supplementary Materials for this manuscript include the following:**

Tables S1 to S11 (Excel files)

Supplementary Figures and Tables
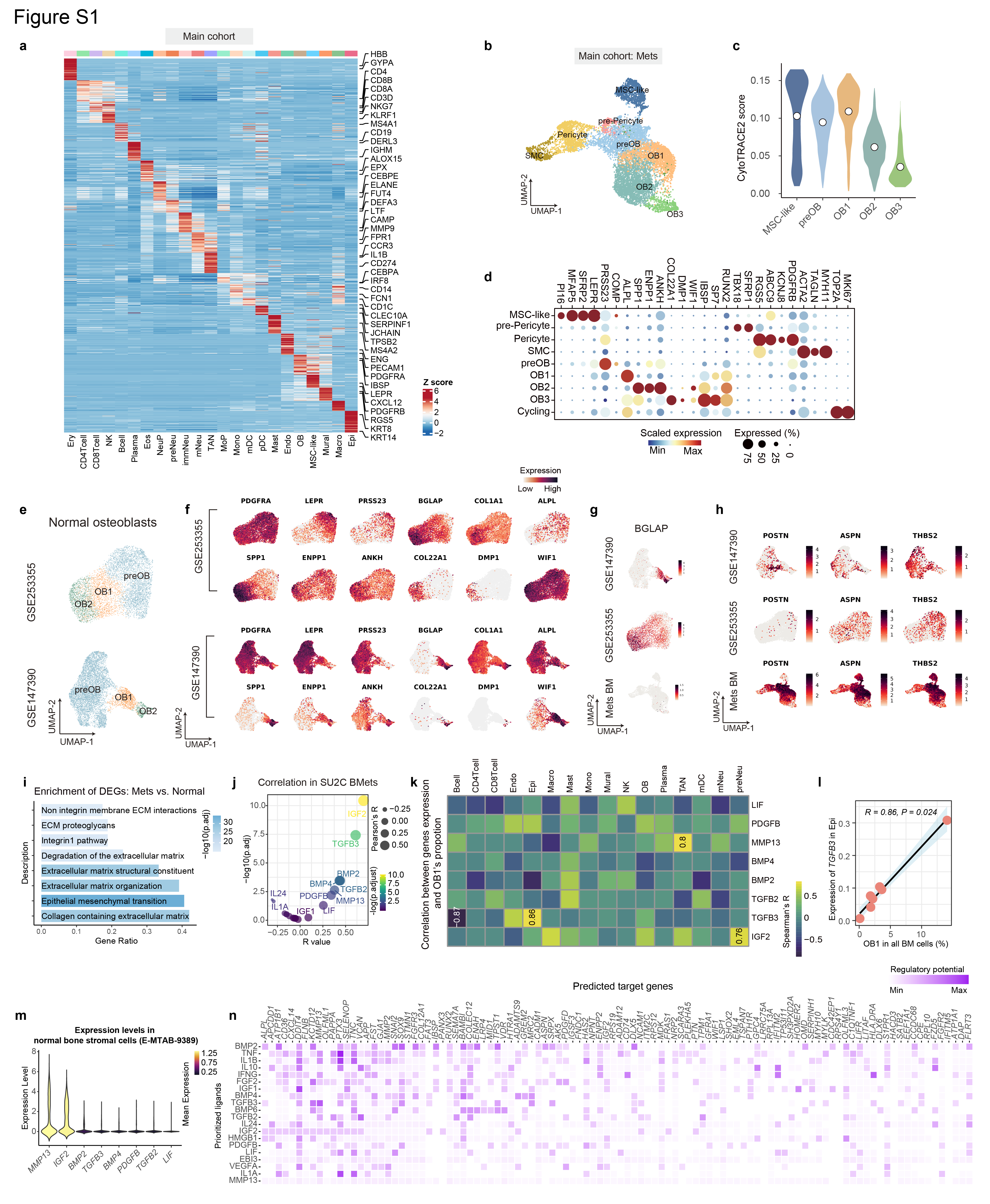

**Fig. S1. Characterization and trajectory analysis of osteoblast subtypes.**

(a) Heatmap showing the expression of the top marker genes for each major cell type identified in the main cohort. Color intensity represents the row-scaled mean expression of each gene across cell types. (b) UMAP of mesenchymal populations from metastatic BM, colored by subtype. (c) Violin plots showing CytoTRACE2 scores across mesenchymal and osteoblast subtypes; white dots indicate median values. (d) Dot plot showing canonical marker-gene expression across mesenchymal, osteoblast and cycling populations. Dot size indicates the percentage of cells expressing the gene, and color indicates the scaled average expression. (e) UMAPs of osteoblast populations from two published normal bone marrow (BM) datasets (GSE253355 and GSE147390), annotated according to the osteoblast states defined in the main cohort. (f) Feature plots showing the expression of mesenchymal and osteoblast-lineage markers in the two normal BM datasets. (g, h) Feature plots showing *BGLAP* expression (g) and the expression of *POSTN*, *ASPN* and *THBS2* (h) in the public normal BM datasets and metastatic BM from the main cohort. Color intensity indicates normalized expression. (i) Over-representation analysis of genes differentially expressed in metastatic versus normal osteoblasts. The x-axis indicates the gene ratio, and bar color indicates -log10(adjusted *P* value). (j) Correlation of candidate signaling molecules with the OB1 signature score in Stand Up To Cancer (SU2C) bone-metastasis samples. The x-axis indicates the Pearson correlation coefficient and the y-axis indicates -log10(adjusted *P* value). (k) Heatmap showing Spearman correlations between ligand expression in major BM cell types and the proportion of OB1 cells across samples. Numbers denote correlation coefficients with *P* values below 0.05. (l) Spearman correlation between *TGFB3* expression in epithelial cells and the proportion of OB1 cells in the main cohort. (m) Violin plots showing the expression of candidate ligands in normal BM stromal cells from E-MTAB-9389; violin color indicates the mean normalized expression. (n) NicheNet analysis predicting ligands and target genes associated with the transition from preOB to OB1. Color intensity indicates the predicted regulatory potential.

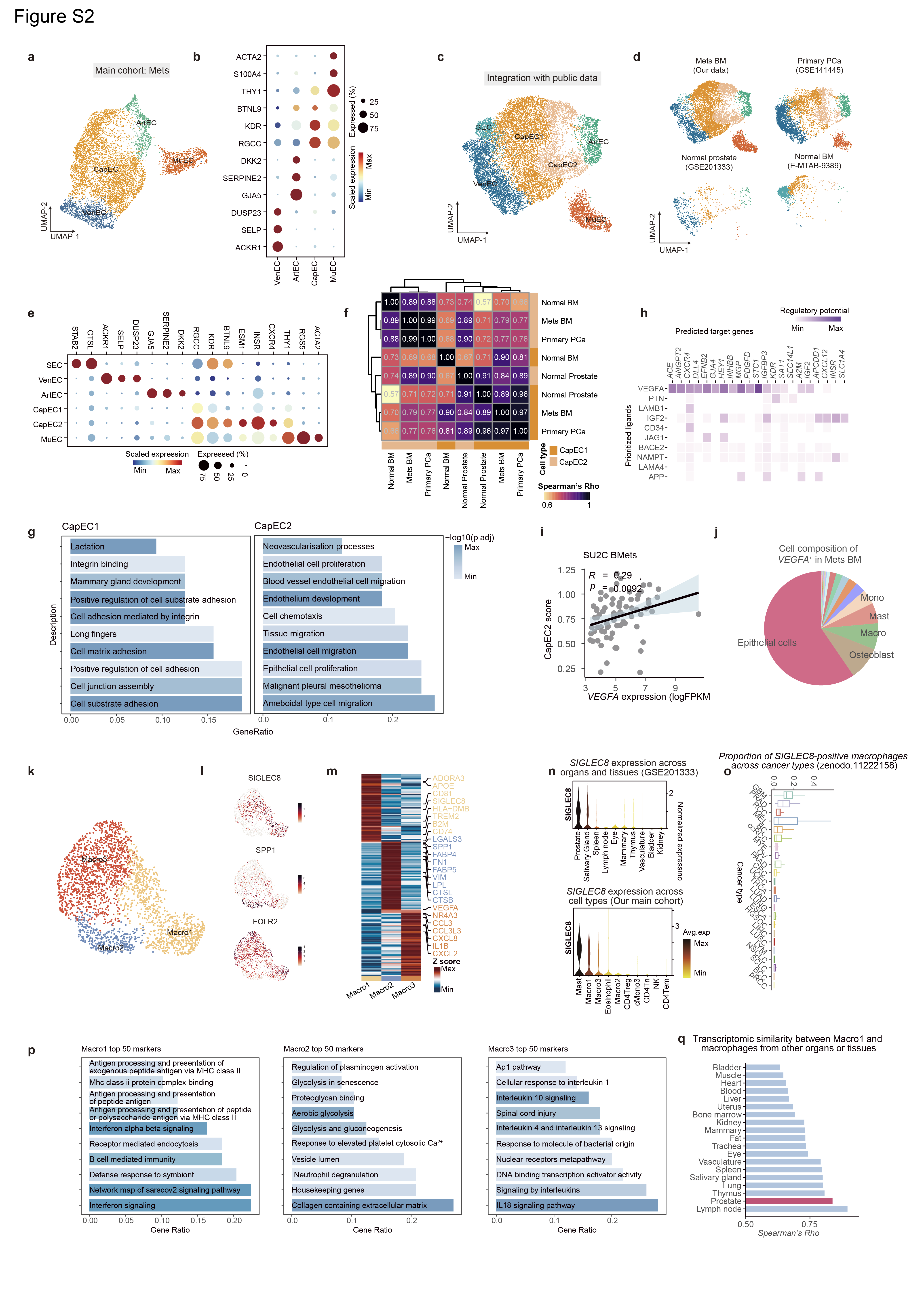

**Fig. S2. Characterization of endothelial and macrophage subtypes in the BM.**

(a) UMAP of endothelial cells (ECs) from metastatic bone marrow (BM) in the main cohort, colored by subtype. (b) Dot plot showing canonical marker-gene expression across EC subtypes. Dot size indicates the percentage of cells expressing the gene, and color indicates the scaled average expression. (c) UMAP of ECs integrated from metastatic BM, primary prostate cancer, normal prostate and normal BM datasets, colored by subtype. (d) Dataset-specific UMAPs of the integrated EC populations. (e) Dot plot showing marker-gene expression across integrated EC subtypes. (f) Spearman-correlation matrix showing transcriptomic similarity between EC subtype profiles across tissues and datasets. (g) Over-representation analysis of marker genes from CapEC1 and CapEC2. The x-axis indicates the gene ratio, and bar color indicates -log10(adjusted *P* value). (h) NicheNet analysis predicting ligands and target genes associated with CapEC2 remodeling. Color intensity indicates the predicted regulatory potential. (i) Pearson correlation between *VEGFA* expression and the CapEC2 signature score in Stand Up To Cancer (SU2C) bone-metastasis samples. (j) Cellular composition of *VEGFA*-expressing cells in metastatic BM; slice size indicates the contribution of each cell type. (k) UMAP of macrophage subtypes from the main cohort. (l) Feature plots showing *SIGLEC8*, *SPP1* and *FOLR2* expression across macrophages. (m) Heatmap showing genes differentially expressed across macrophage subtypes; selected markers are labeled. Color intensity represents row-scaled mean expression. (n) Violin plots showing *SIGLEC8* expression in macrophages across normal organs and tissues in GSE201333 (top) and across major cell types in the main cohort (bottom). (o) Proportions of *SIGLEC8*-positive macrophages across cancer types in a pan-cancer dataset. BC, breast cancer; BCC, basal cell carcinoma; ccRCC, clear cell renal cell carcinoma; CRC, colorectal cancer; ESCA, esophageal carcinoma; GBM, glioblastoma; HGSOC, high-grade serous ovarian carcinoma; LIHC, liver hepatocellular carcinoma; LUAD, lung adenocarcinoma; LUNG, lung cancer; LUSC, lung squamous cell carcinoma; LYM, lymphoma; MEL, melanoma; MYE, myeloma; NSCLC, non-small cell lung cancer; OV, ovarian cancer; PAAD, pancreatic adenocarcinoma; PRAD, prostate adenocarcinoma; pRCC, papillary renal cell carcinoma; RCC, renal cell carcinoma; SCLC, small cell lung cancer; THCA, thyroid carcinoma; UCEC, uterine corpus endometrial carcinoma. (p) Over-representation analysis of the top marker genes of Macro1, Macro2 and Macro3. The x-axis indicates the gene ratio, and bar color indicates -log10(adjusted *P* value). (q) Transcriptomic similarity of Macro1 to macrophages from normal organs and tissues, quantified by Spearman's ρ; similarity to prostate macrophages is highlighted.

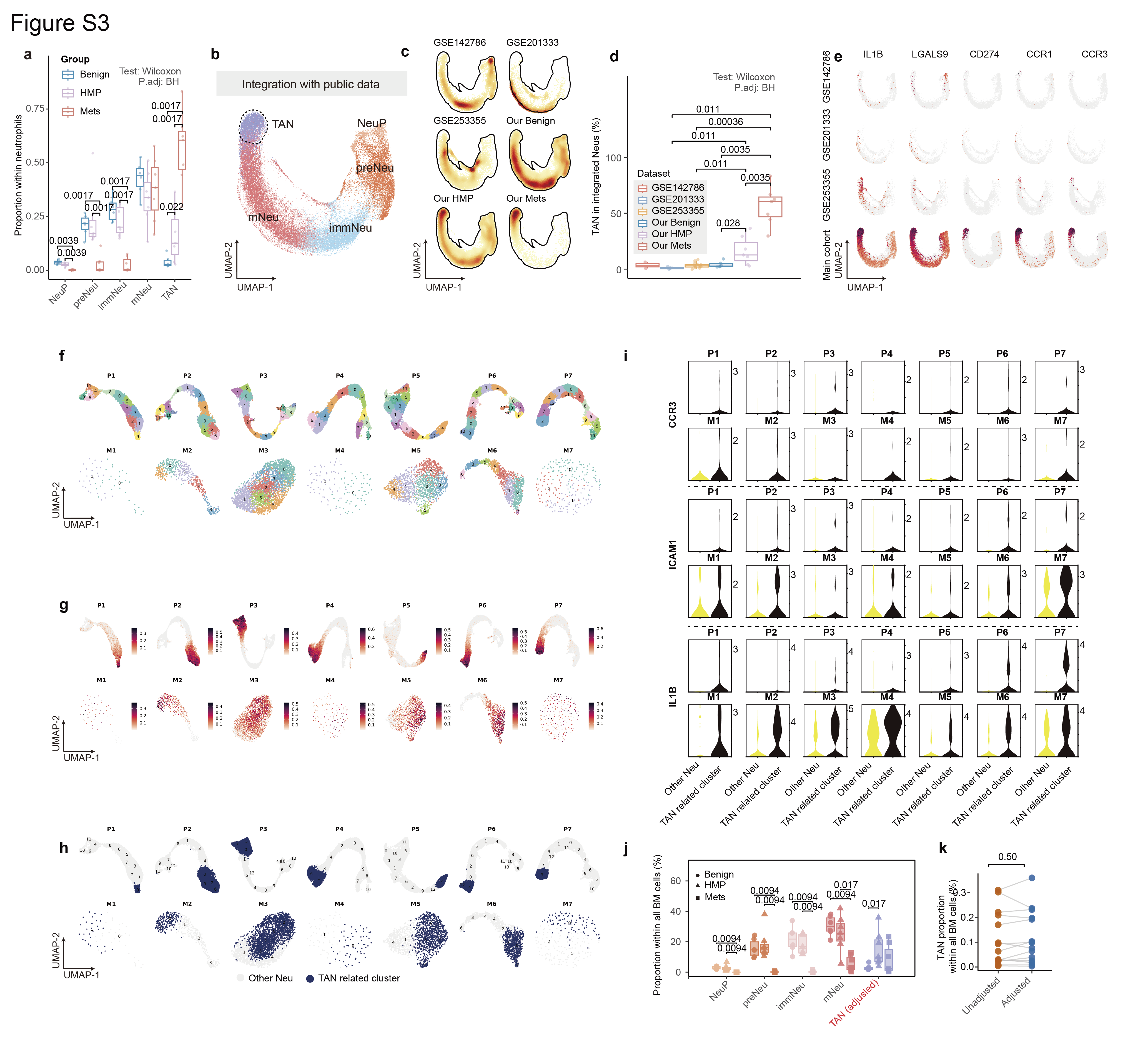

**Fig. S3. Cross-dataset and sample-wise validation of the TAN state.**

(a) Proportions of neutrophil subtypes within the neutrophil compartment across Benign, HMP and Mets samples. Each point represents one sample. (b) UMAP of neutrophils integrated from public datasets (GSE142786, GSE201333 and GSE253355) and the main cohort, colored by subtype. (c) Cell-density maps of the integrated neutrophils by dataset or disease group. (d) Proportion of TANs among integrated neutrophils across datasets and disease groups. (e) Feature plots showing IL1B, LGALS9, CD274, CCR1 and CCR3 expression in neutrophils from each dataset. (f) UMAPs of neutrophils analysed independently within individual HMP and Mets bone marrow (BM) samples. (g) TAN-signature scores across neutrophils in each independently analysed sample. (h) TAN-related clusters identified independently within individual HMP and Mets samples. (i) Expression of CCR3, ICAM1 and IL1B in sample-wise TAN-related clusters compared with other neutrophils. (j) Patient-level proportions of neutrophil subtypes across Benign, HMP and Mets samples, with TAN abundance recalculated using the sample-wise TAN assignments. (k) Paired comparison of TAN proportions among total BM cells obtained using the integrated annotation and the sample-wise TAN reassignment. *Comparisons in panels a, d and j were performed using two-sided Wilcoxon rank-sum tests; the paired comparison in panel k was performed using a two-sided Wilcoxon signed-rank test.*

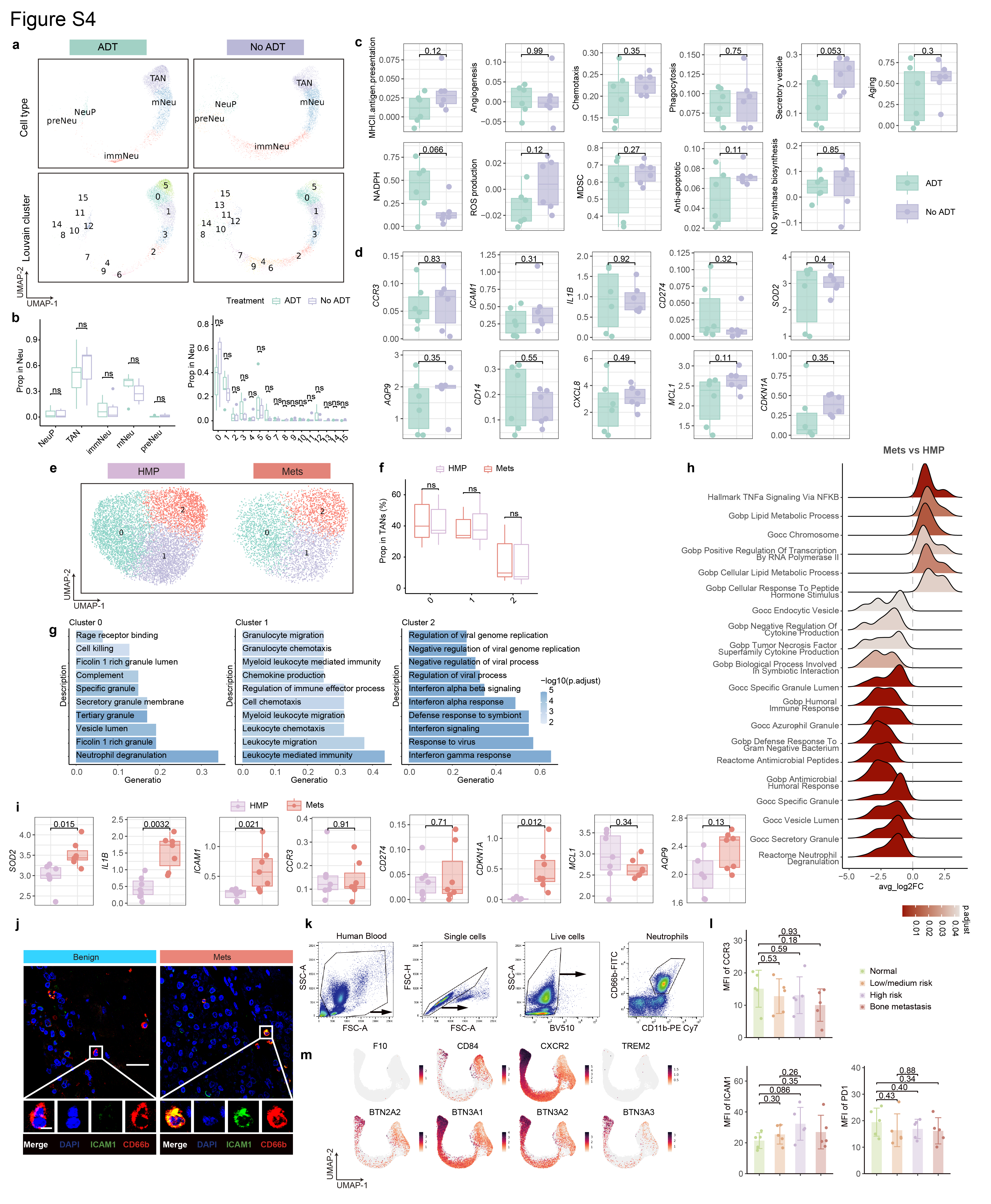

**Fig. S4. Phenotypic and transcriptional characterization of TANs across disease stages and treatment status.**

(a) UMAPs showing neutrophil annotations (top) and Louvain clusters (bottom) obtained with or without androgen deprivation therapy (ADT) treatment. (b) Comparison of neutrophil-subtype and Louvain-cluster proportions between the ADT and no-ADT groups. (c, d) Comparison of neutrophil functional-module scores (c), TAN-associated genes (d) in TANs between the ADT and no-ADT groups. (e) UMAPs of TAN subclusters in HMP and Mets samples. (f) Proportions of TAN clusters 0-2 in HMP and Mets samples. (g) Over-representation analysis of the top markers of TAN clusters 0-2. Bar length indicates the gene ratio, and bar color indicates -log10(adjusted *P* value). (h) Ridge plots of GSEA results comparing Mets and HMP TANs. Curves show the distributions of average log2 fold changes of genes within each gene set and are colored according to adjusted P values. The dashed line indicates an average log2 fold change of zero. (i) Expression of selected TAN-associated genes in HMP and Mets TANs. (j) Representative multiplex immunofluorescence images of bone marrow (BM) from Benign and Mets groups stained for DAPI, ICAM1 and CD66b. Scale bars, 25 µm in the main images and 5 µm in the insets. (k) Flow-cytometry gating strategy for live, single CD66b-positive neutrophils from human peripheral blood. (l) CCR3, ICAM1 and PD-1 mean fluorescence intensities (MFIs) in peripheral-blood neutrophils from individuals without prostate cancer (normal) and patients with low/medium risk, high risk or established bone metastasis. Each dot represents one participant; n = 5 participants per group. (m) Feature plots showing previously reported prostate-cancer TAN markers and butyrophilin-family genes in BM neutrophils. *Comparisons were performed using two-sided Student’s t-tests (b-d, f, i and l), and P values were adjusted using the Benjamini-Hochberg method. “ns” in panel b means “no statistical significance (P > 0.05)”.*

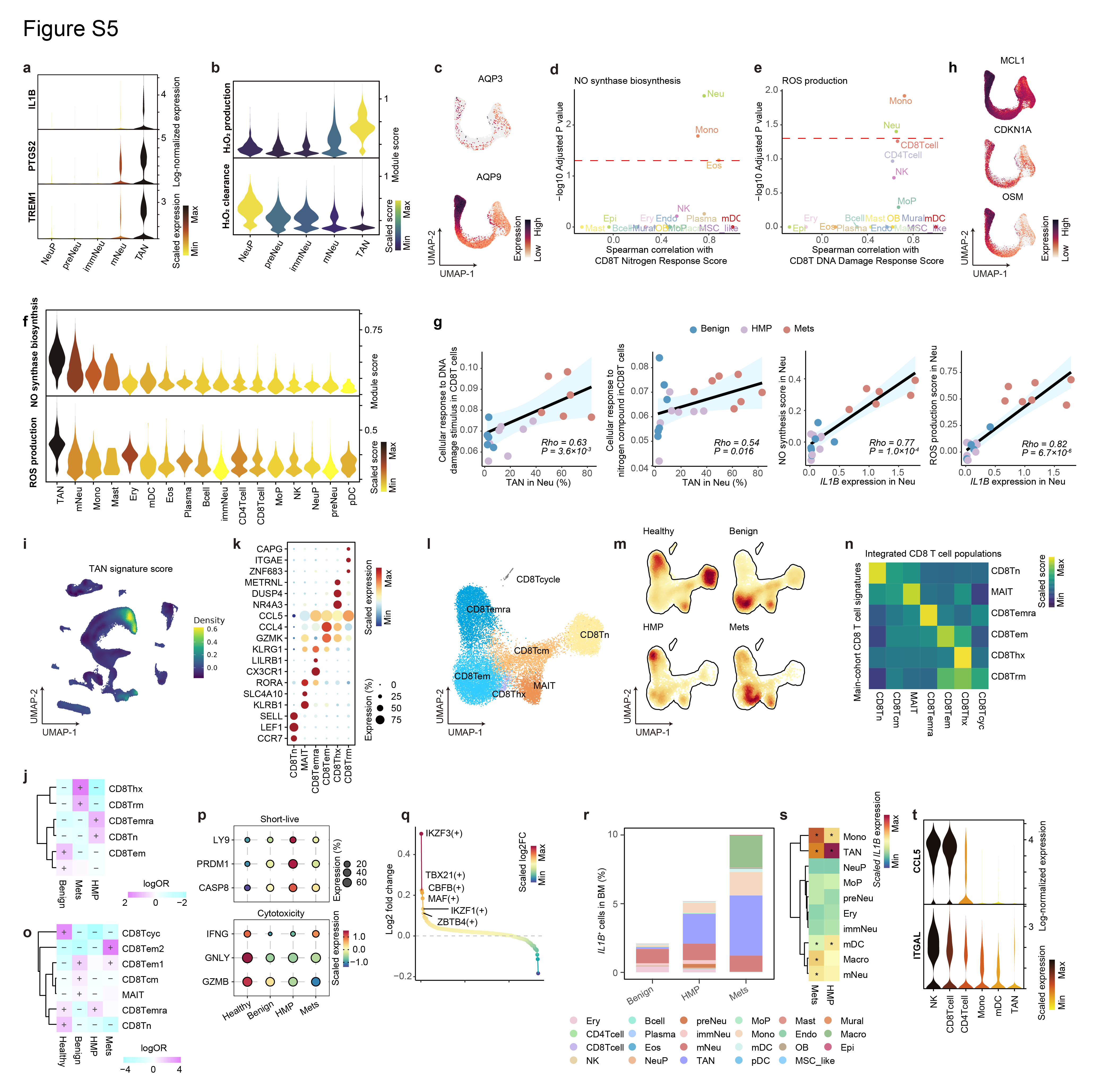

**Fig. S5. TAN-associated oxidative stress and CD8 T cell remodeling across disease stages.**

(a) Violin plots showing *IL1B*, *PTGS2* and *TREM1* expression across neutrophil subtypes. Violin color indicates the mean normalized expression. (b) Violin plots showing hydrogen-peroxide-production and hydrogen-peroxide-clearance signature scores across neutrophil subtypes. (c) Feature plots showing *AQP3* and *AQP9* expression in neutrophils. (d, e) Associations between nitric-oxide-synthesis scores in each major bone marrow (BM) cell type and the nitrogen-response score in CD8 T cells (d), and between ROS-production scores in each major BM cell type and the DNA-damage-response score in CD8 T cells (e). The x-axis indicates the Spearman correlation coefficient and the y-axis indicates −log10(adjusted *P* value); the dashed line denotes adjusted *P* = 0.05. Statistical correlations were performed using patient-level values. (f) Violin plots showing nitric-oxide-synthesis and ROS-production scores across major BM cell types; violin color indicates the mean score. (g) Spearman correlations between TAN abundance and DNA-damage or nitrogen-response programs in CD8 T cells, and between neutrophil *IL1B* expression and nitric-oxide-synthesis or ROS-production programs in neutrophils. Points are colored by disease group. Each point represents one patient sample. (h) Feature plots showing *MCL1*, *CDKN1A* and *OSM* expression in neutrophils. (i) Density of TAN signature scores across all BM cells in the main cohort. (j) Differential abundance of CD8 T cell subtypes across Benign, HMP and Mets samples. Color indicates the log odds ratio, and “+” or “-” denotes significant enrichment or depletion, respectively. (k) Dot plot showing canonical marker-gene expression across CD8 T cell subtypes. Dot size indicates the percentage of cells expressing the gene, and color indicates the scaled average expression. (l) UMAP of CD8 T cells integrated with public BM datasets, colored by subtype. (m) Cell-density maps of integrated CD8 T cells across Healthy, Benign, HMP and Mets groups. (n) Heatmap showing the signature scores of CD8 T cell states defined in the main cohort when projected onto the integrated CD8 T cell populations. (o) Differential abundance of integrated CD8 T cell subtypes across Healthy, Benign, HMP and Mets groups, displayed as in panel j. (p) Dot plots showing short-lived and cytotoxicity-associated genes in CD8Temra cells across disease stages. (q) Differentially activated regulons in CD8Temra cells between HMP and healthy BM. Each point represents a regulon. The color intensity represents the scaled log2 fold change. Regulons and their activities were identified by SCENIC. (r) Cell type-resolved percentages of *IL1B*-positive cells among all BM cells across Benign, HMP and Mets samples. (s) Heatmap showing mean *IL1B* expression across cell types expressing *IL1B* in HMP and Mets BM. Within each disease stage, *IL1B* expression in each cell type was compared with that in all remaining cell types using a Wilcoxon rank-sum test. Asterisks indicate significantly higher *IL1B* expression (Bonferroni-adjusted *P* < 0.05). (t) Violin plots showing the six cell types with the highest average expression of *CCL5* and *ITGAL* in the main BM cohort. Violin color indicates the average normalized expression of each gene across cell types. *Correlations in panels d, e and g were assessed using Spearman correlation; adjusted P values in panels d and e were obtained using the Benjamini-Hochberg method. Wilcoxon rank-sum test was used when comparing regulon activity score (q). Differential abundance in panels j and o was assessed using two-sided Fisher’s exact tests with Benjamini–Hochberg adjustment for multiple comparisons.*

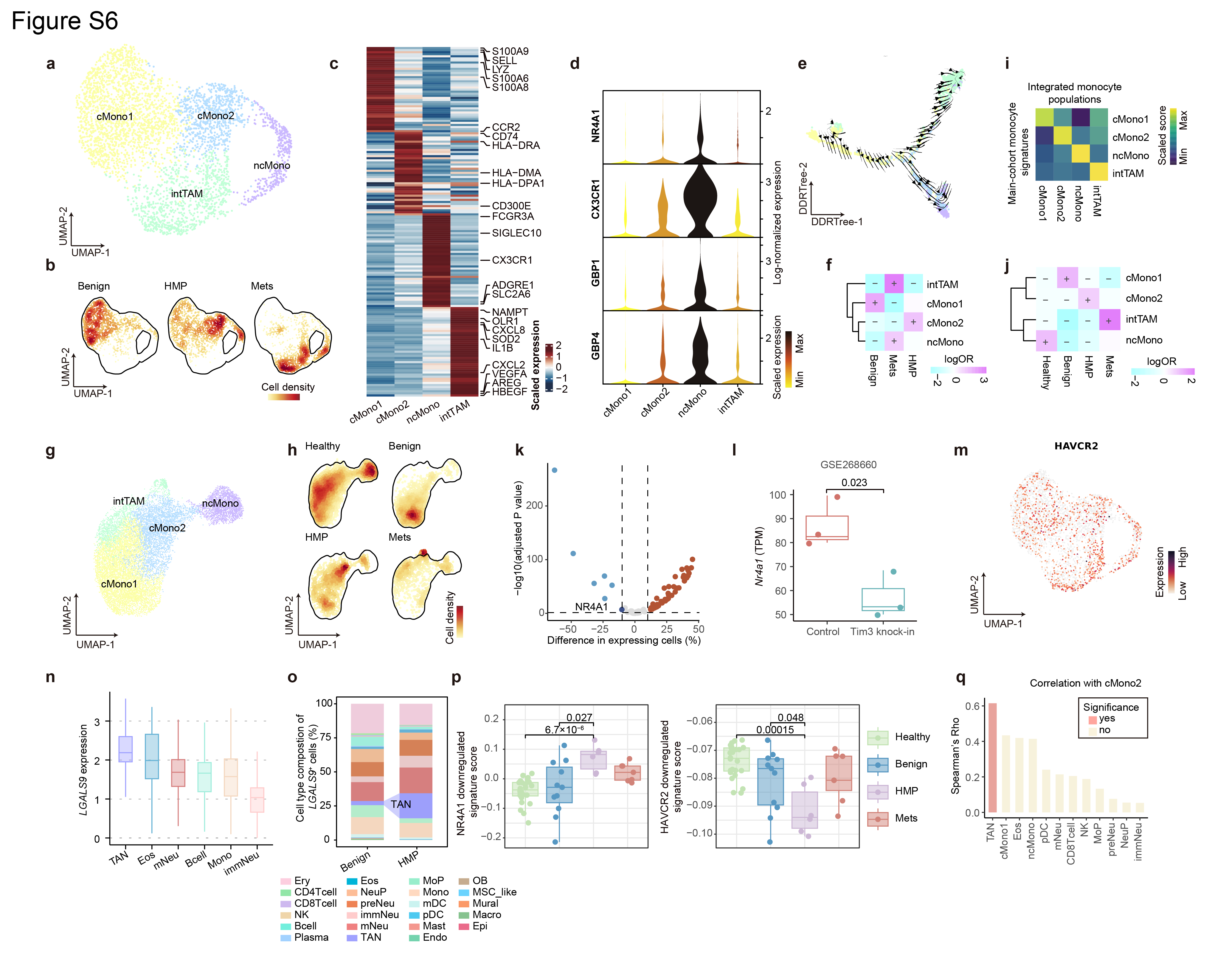

**Fig. S6. Detailed characterization of monocyte alterations.**

(a) UMAP of monocyte subtypes from the main cohort, colored by subtype. (b) Cell-density maps of monocytes across Benign, HMP and Mets samples. (c) Heatmap showing genes differentially expressed across monocyte subtypes; selected markers are labeled. Color intensity represents row-scaled mean expression. (d) Violin plots showing *NR4A1*, *CX3CR1*, *GBP1* and *GBP4* expression across monocyte subtypes. Violin color indicates the scaled mean expression. (e) DDRTree embedding of monocytes with overlaid RNA-velocity vectors indicating the inferred direction of differentiation. (f) Differential abundance of monocyte subtypes across Benign, HMP and Mets samples. Color indicates the log odds ratio, and '+' or '-' denotes significant enrichment or depletion, respectively. (g) UMAP of monocytes integrated with public bone marrow (BM) datasets, colored by subtype. (h) Cell-density maps of integrated monocytes across Healthy, Benign, HMP and Mets groups. (i) Heatmap showing the signature scores of monocyte subtypes defined in the main cohort when projected onto integrated monocyte populations. (j) Differential abundance of integrated monocyte subtypes across Healthy, Benign, HMP and Mets groups, displayed as in panel f. (k) Differential expression analysis of genes associated with non-classical-monocyte differentiation in cMono2 cells from HMP versus Healthy BM. The x-axis indicates the difference in the percentage of expressing cells and the y-axis indicates -log10(adjusted *P* value); NR4A1 is labeled. (l) *Nr4a1* expression in control and Tim3-knock-in monocytes from GSE268660. (m) Feature plot showing *HAVCR2* expression across monocytes. (n) *LGALS9* expression across the indicated BM cell types; expression was calculated among *LGALS9*-positive cells. (o) Cellular composition of *LGALS9*-positive cells in Benign and HMP BM. (p) Signature scores of genes downregulated by NR4A1 or HAVCR2 in integrated cMono2 cells across Healthy, Benign, HMP and Mets groups. (q) Spearman correlations between cMono2 abundance and other BM cell populations; significant correlations are highlighted (P < 0.05 was considered significant). *For panels l and p, comparisons were performed using two-sided Student's t-tests and P values were adjusted using the Benjamini-Hochberg method. Differential abundance in panels f and j was assessed using two-sided Fisher’s exact tests with Benjamini-Hochberg adjustment for multiple comparisons.*

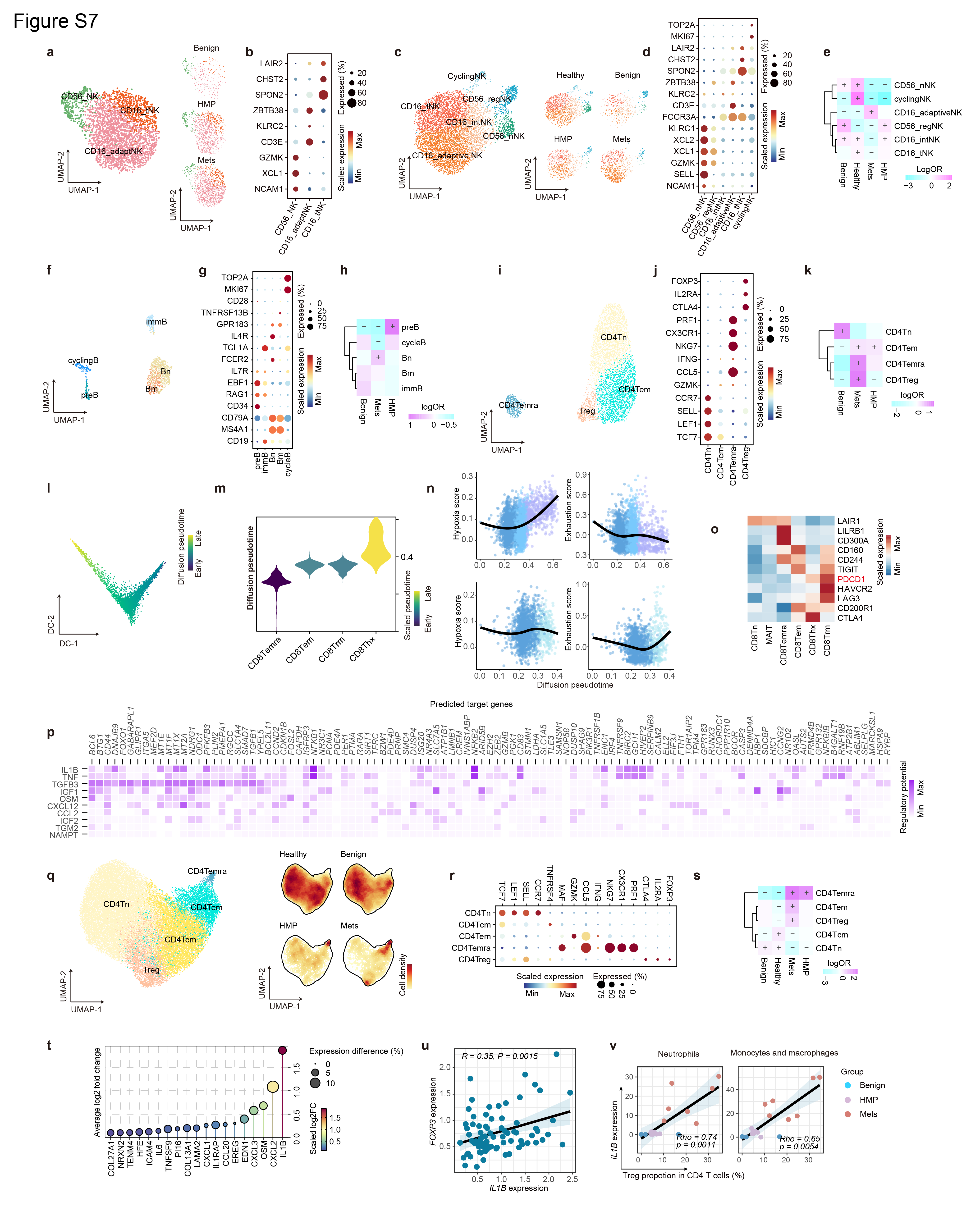

**Fig. S7. UMAP visualization, marker expression and population distribution of immune-cell subsets.**

(a) UMAP of NK cells from the main cohort, colored by subtype (left), with group-specific embeddings for Benign, HMP and Mets samples (right). (b) Dot plot showing canonical marker-gene expression across NK cell subtypes. Dot size indicates the percentage of cells expressing the gene, and color indicates the scaled average expression. (c) UMAP of NK cells integrated with public bone marrow (BM) datasets, colored by subtype, with cell-density maps across Healthy, Benign, HMP and Mets groups. (d) Dot plot showing marker-gene expression across integrated NK cell subtypes. (e) Differential abundance of integrated NK cell subtypes across disease groups. Color indicates the log odds ratio, and '+' or '-' denotes significant enrichment or depletion, respectively. (f) UMAP of B cell subtypes from the main cohort. (g) Dot plot showing canonical marker-gene expression across B cell subtypes. (h) Differential abundance of B cell subtypes across Benign, HMP and Mets samples. (i) UMAP of CD4 T cell subtypes from the main cohort. (j) Dot plot showing canonical marker-gene expression across CD4 T cell subtypes. (k) Differential abundance of CD4 T cell subtypes across Benign, HMP and Mets samples. (l) Diffusion map of CD8Temra, CD8Tem, CD8Trm and CD8Thx cells, colored by diffusion pseudotime (DPT). (m) Distribution of DPT across CD8 T cell subtypes; violin color indicates the mean DPT. (n) Changes in selected functional scores along DPT; black lines indicate fitted trends. (o) Heatmap showing inhibitory-receptor expression across CD8 T cell subtypes. (p) NicheNet analysis predicting ligands and target genes associated with the transition from CD8Tem to CD8Thx. Color intensity indicates the predicted regulatory potential. (q) UMAP of CD4 T cells integrated with public BM datasets, colored by subtype, with cell-density maps across Healthy, Benign, HMP and Mets groups. (r) Dot plot showing canonical marker-gene expression across integrated CD4 T cell subtypes. (s) Differential abundance of integrated CD4 T cell subtypes across disease groups. (t) Ligands predicted to act on Tregs and upregulated in Mets versus HMP samples. Dot position indicates average log2 fold change, dot size indicates the difference in the percentage of expressing cells, and color indicates the scaled average log2 fold change. *(u) Pearson correlation between IL1B and FOXP3 expression in Stand Up To Cancer (SU2C) bone-metastasis samples. (v) Spearman correlations between the proportion of Tregs within CD4 T cells and IL1B expression in neutrophils or monocytes/macrophages in the main cohort. Points are colored by disease group. Differential abundance in panels e, h, k and s was assessed using two-sided Fisher’s exact tests with Benjamini-Hochberg adjustment for multiple comparisons.*

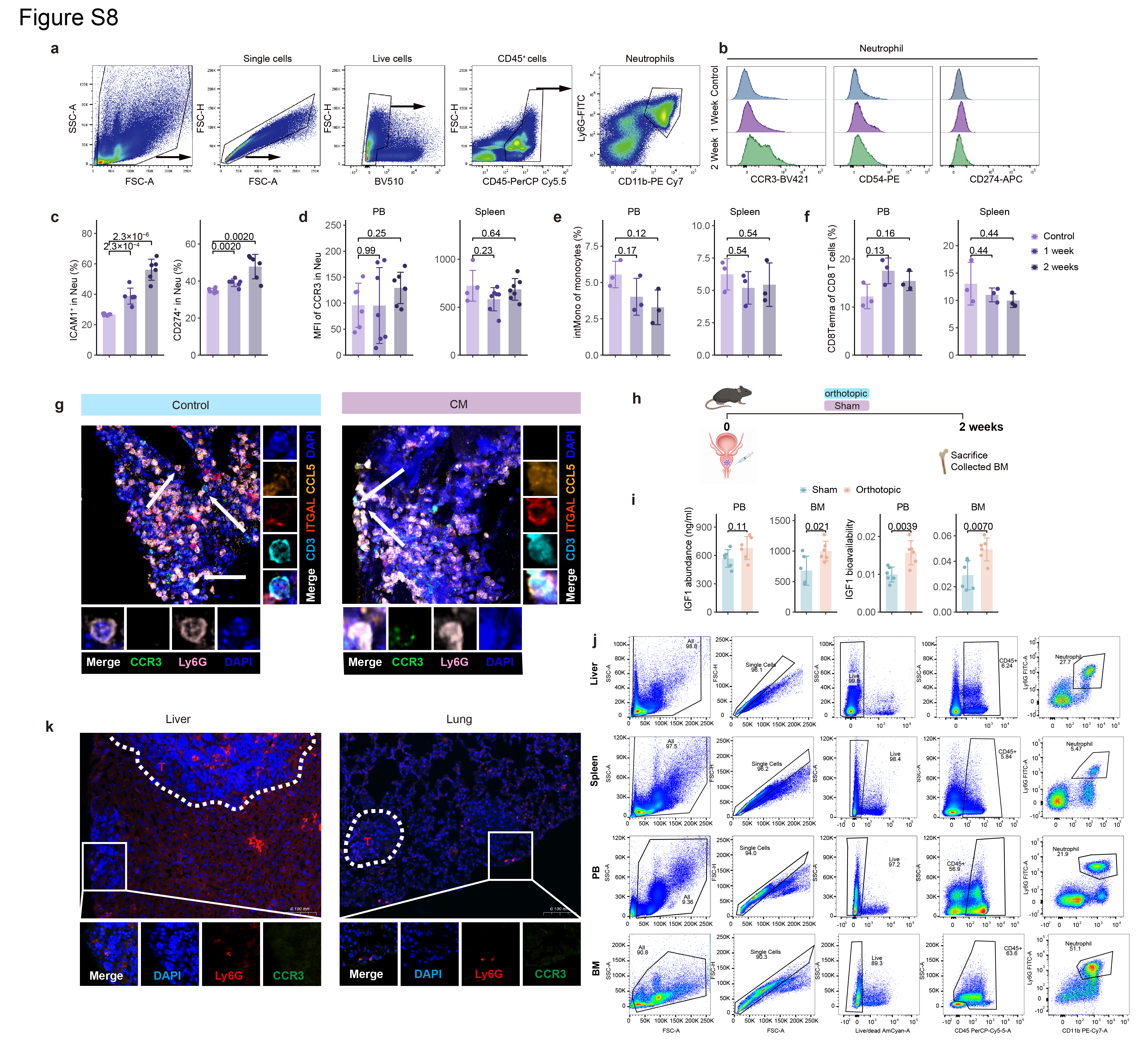

**Fig. S8. Systemic and BM-specific alterations in conditioned-medium and orthotopic prostate cancer models.**

(a) Flow-cytometry gating strategy for live CD45-positive CD11b-positive Ly6G-positive mouse bone marrow (BM) neutrophils. (b) Representative flow-cytometry distributions of CCR3, CD54 and CD274 in BM neutrophils from control mice and mice treated with conditioned medium (CM) for 1 or 2 weeks. (c) Frequencies of ICAM1-positive and CD274-positive BM neutrophils during CM-induced BM preconditioning. Each dot represents one mouse. (d) CCR3 MFI in neutrophils from peripheral blood and spleen at 1 and 2 weeks after conditioned medium (CM) treatment. For peripheral blood, control, n = 6 mice; 1 week and 2 weeks, n = 7 mice per group. For spleen, control, n = 4 mice; 1 week and 2 weeks, n = 7 mice per group. (e) Frequencies of intermediate monocytes among total monocytes in peripheral blood and spleen; n = 3 mice per group. (f) Frequencies of CD8Temra cells among total CD8 T cells in peripheral blood and spleen; n = 3 mice per group. (g) Representative multiplex immunofluorescence images showing spatial proximity between CCR3-positive Ly6G-positive neutrophils and CCL5-positive ITGAL-positive CD3-positive T cells in BM from CM-preconditioned mice. Scale bars, 50 µm in the main images and 5 µm in the insets. (h) Experimental design of the orthotopic prostate cancer model. BM was collected 2 weeks after implantation of TRAMP-C1 cells or sham surgery. (i) IGF1 abundance and bioavailability in peripheral blood and BM from sham-operated and orthotopic tumor-bearing mice; n = 6 mice per group. (j) Representative flow-cytometry gating of live CD45-positive CD11b-positive Ly6G-positive neutrophils from liver, spleen, peripheral blood and BM. (k) Representative immunofluorescence images of spontaneous liver and lung prostate cancer metastatic lesions from previous research (Guo et al., 2025), stained for DAPI, Ly6G and CCR3. Dashed outlines indicate tumor regions. Liver: n = 4; lung: n = 3. Scale bars, 100 µm. *Comparisons in panels c–f and i were performed using two-sided Student’s t-tests, with Benjamini-Hochberg adjustment where multiple comparisons were performed.*

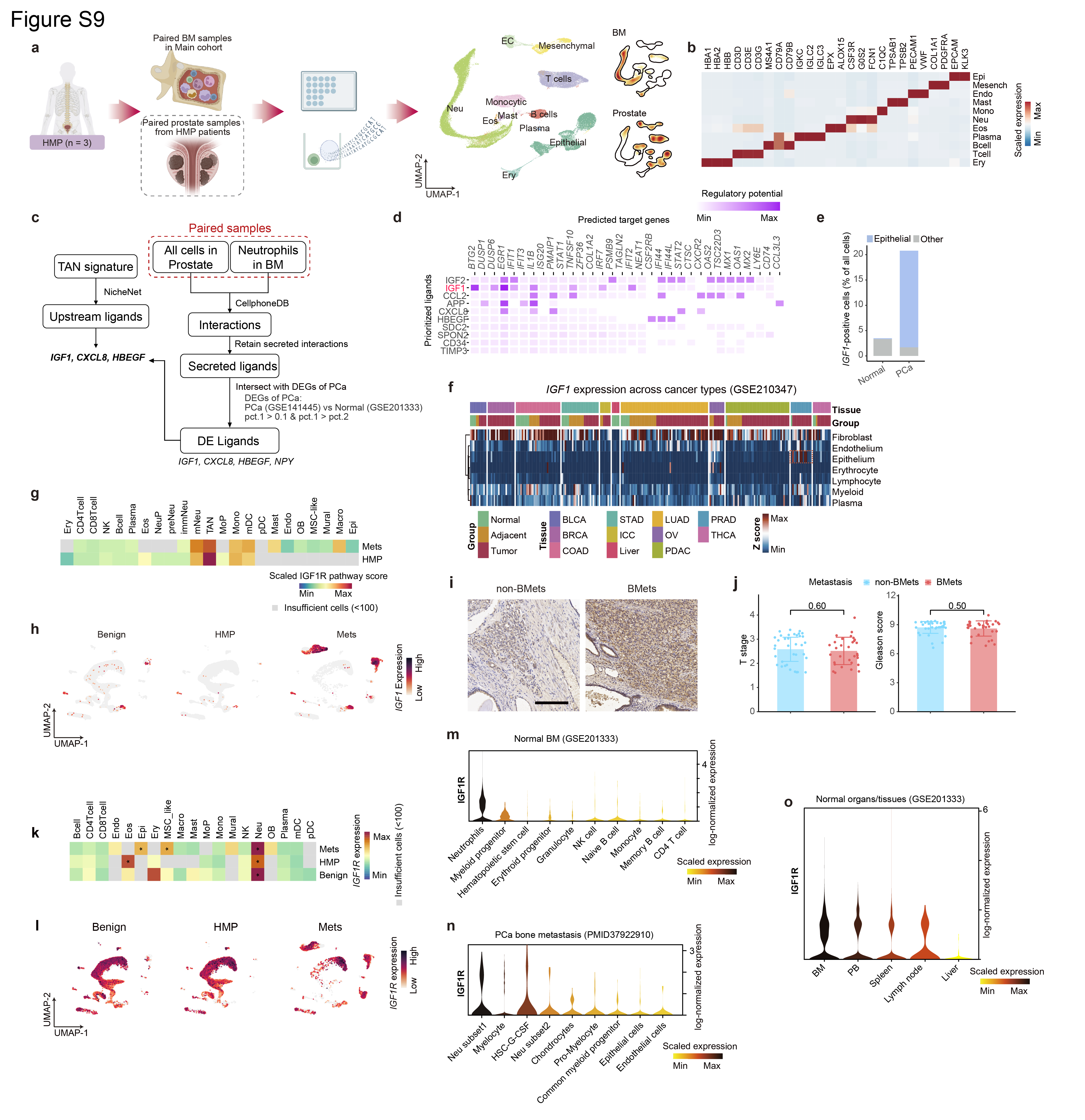

**Fig. S9. Prioritization of tumor-associated IGF1 signaling and validation of IGF1R expression across bone marrow cell populations.**

(a) Schematic of paired single-cell profiling of bone marrow (BM) and primary-prostate samples from 3 HMP patients. UMAPs on the right show major cell types and tissue-specific cell-density distributions. (b) Heatmap showing marker-gene expression across major cell types in the paired samples. Color intensity represents row-scaled mean expression. (c) Analytical strategy integrating NicheNet, CellPhoneDB and differential-expression analysis of prostate cancer versus normal prostate to identify secreted prostate-derived ligands predicted to act on BM neutrophils. (d) NicheNet analysis showing prioritized ligands and predicted target genes associated with the TAN program in Mets BM from the main cohort. Color intensity indicates predicted regulatory potential. (e) Epithelial and non-epithelial composition of *IGF1*-positive cells in normal prostate and prostate cancer tissues. The epithelial fraction among *IGF1*-positive cells was compared between groups using a two-sided beta-binomial regression based on sample-level epithelial and non-epithelial *IGF1*-positive cell counts, with samples treated as biological replicates (*P* = 2.86 × 10^-4^). (f) *IGF1* expression across cell populations and cancer types in the GSE210347 pan-cancer dataset. Rows denote cell populations and columns denote samples; annotations indicate tissue and tumor status. BLCA: Bladder urothelial carcinoma; BRCA: Breast cancer; COAD: Colon adenocarcinoma; STAD: Stomach adenocarcinoma; ICC: Intrahepatic cholangiocarcinoma; LUAD: Lung adenocarcinoma; OV: Ovarian cancer; PDAC: Pancreatic ductal adenocarcinoma; PRAD: Prostate adenocarcinoma; THCA: Thyroid carcinoma. (g) IGF1R pathway scores across major BM cell types and disease stages. Gray indicates cell type-group combinations with fewer than 100 cells. (h) Feature plots showing IGF1 expression across all BM cells in Benign, HMP and Mets samples of our main cohort. (i) Representative IGF1 immunohistochemistry in primary prostate tumors from patients without bone metastasis (non-BMets) and patients with bone metastasis (BMets). Scale bar, 100 µm. (j) Pathological T stage and Gleason score in patients without or with bone metastasis. Each dot represents one patient; non-BMets, n = 36; BMets, n = 33. (k) *IGF1R* expression levels across major BM cell types and disease stages. Within each disease stage, *IGF1R* expression in each cell type was compared with that in all remaining cell types using a Wilcoxon rank-sum test. Asterisks indicate significantly higher IGF1R expression (Bonferroni-adjusted *P* < 0.05). Gray indicates cell type–group combinations containing fewer than 100 cells. (l) Feature plots showing *IGF1R* expression across the integrated BM UMAP in Benign, HMP and Mets samples. (m) *IGF1R* expression across cell types in normal human BM from GSE201333. (n) *IGF1R* expression across cell types in an independent prostate-cancer bone-metastasis dataset (PMID: 37922910). (o) *IGF1R* expression in neutrophils from human BM, blood, spleen, lymph node and liver in GSE201333. *Statistical tests and exact P values are shown in the panels. P values from multiple Student’s t-test comparisons were adjusted using the Benjamini-Hochberg method.*

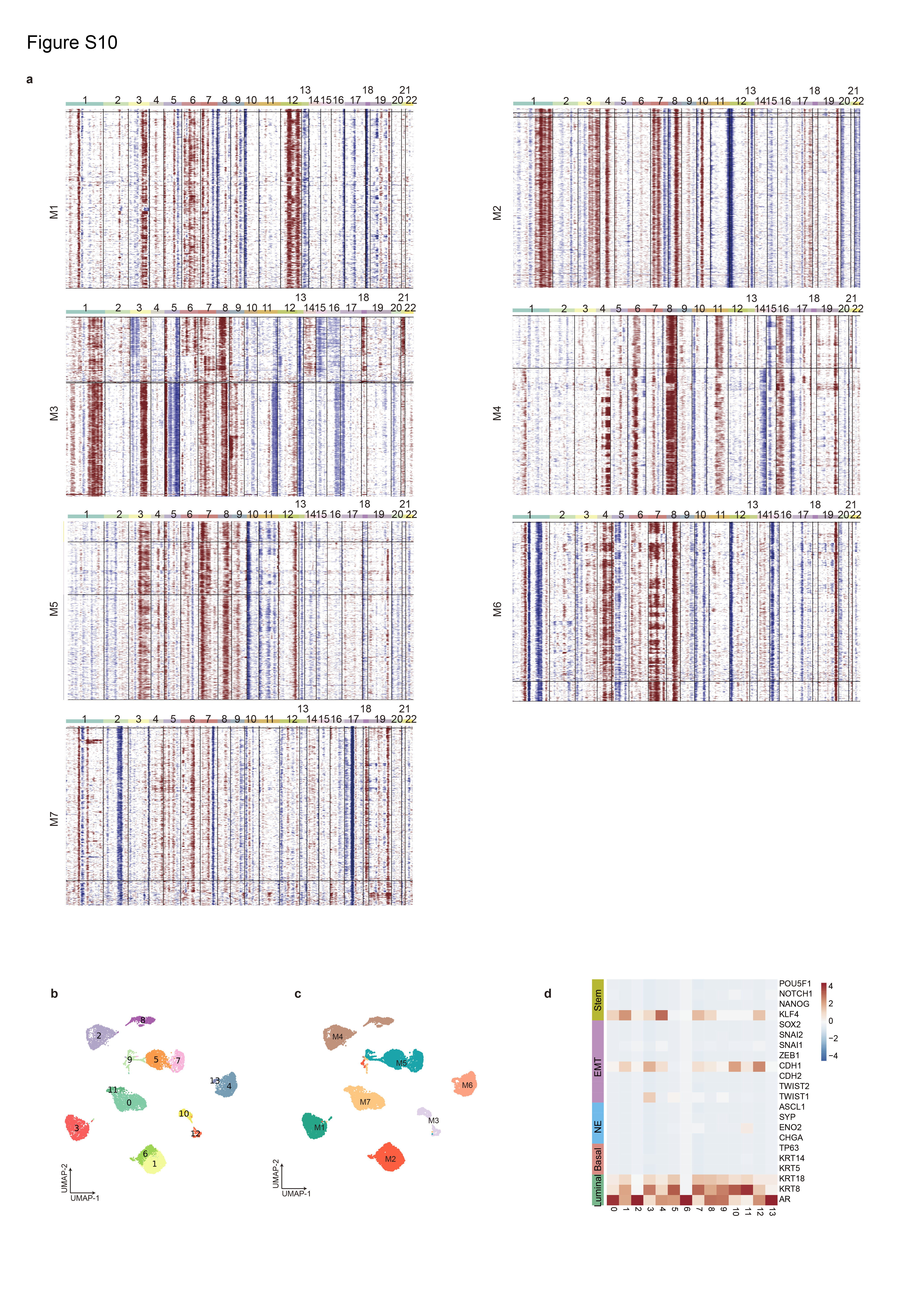

**Fig. S10. Copy-number-variation analysis and heterogeneity of metastatic epithelial cells.**

(a) Inferred copy-number-variation profiles of epithelial cells from metastatic bone marrow samples M1-M7 across chromosomes 1-22. Non-malignant reference cells were used to define the baseline; red and blue indicate relative chromosomal gains and losses, respectively. (b,c) UMAPs of epithelial cells without batch correction, colored by Louvain cluster (b) or patient identity (c). (d) Heatmap showing the expression of luminal, basal, neuroendocrine, epithelial-mesenchymal-transition and stemness-associated genes across epithelial clusters. Color intensity represents the scaled mean expression.

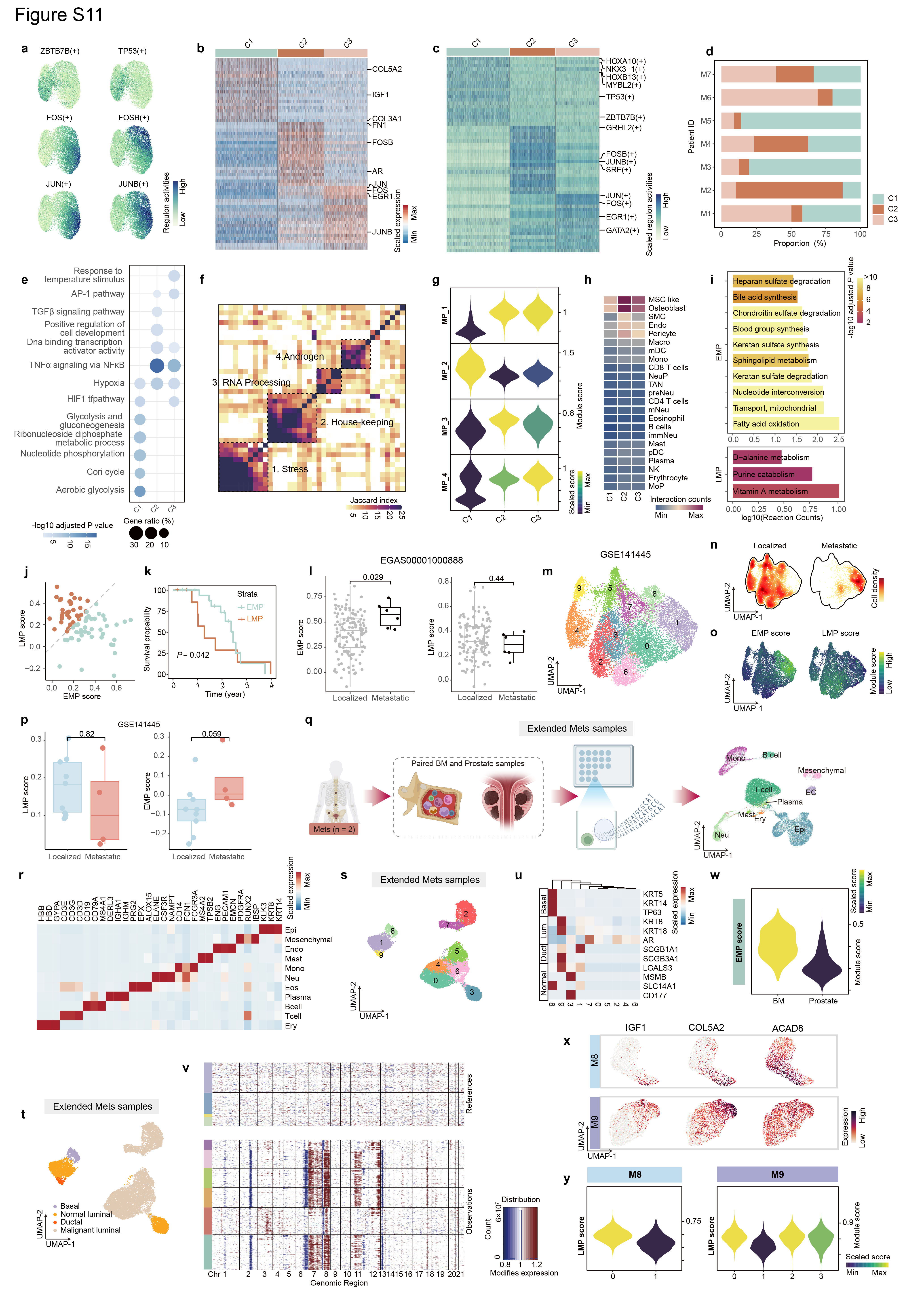

**Fig. S11. Characterization of metastatic epithelial subpopulations and their clinical associations.**

(a) Feature plots showing the activity of selected regulons across metastatic epithelial cells. (b, c) Heatmaps showing differentially expressed genes (b) and differential regulon activity (c) across epithelial clusters C1-C3. Color intensity represents row-scaled expression or regulon activity. (d) Proportional composition of epithelial clusters C1-C3 across metastatic bone marrow (BM) samples M1-M7. (e) Over-representation analysis of DEGs across epithelial subclusters using gene-set collections from MSigDB. Each row represents an enriched pathway. Dot size indicates the gene ratio, and color indicates −log10(adjusted P value). (f) Heatmap showing the similarity between gene programs identified by NMF-based clustering, quantified using the Jaccard index. Meta-programs, representing groups of similar programs, are indicated by dashed rectangles. (g) Violin plots showing meta-program scores across epithelial clusters; violin color indicates the scaled mean score. (h) Heatmap showing inferred interaction counts between epithelial clusters C1-C3 (columns) and other major BM cell types (rows). Color intensity indicates the number of inferred interactions. (i) Metabolic-subsystem enrichment analysis of differentially regulated metabolic reactions between EMP and LMP epithelial cells. Bar length indicates the log10-transformed reaction count, and color indicates −log10(adjusted P value). (j) EMP and LMP signature scores in Stand Up To Cancer (SU2C) bone-metastasis samples. The dashed diagonal separates EMP-dominant and LMP-dominant samples. (k) Overall survival of SU2C bone-metastasis samples stratified by EMP- or LMP-dominant status; the *P* value was calculated using the Gehan-Breslow test. (l) EMP and LMP scores in localized and metastatic prostate cancer samples from EGAS00001000888. (m-o) UMAP plots of epithelial cells from the GSE141445 primary prostate cancer dataset, colored according to (m) Louvain-defined subclusters. Corresponding plots show cell density (n), stratified by patient metastatic status (localized vs. metastatic), and EMP and LMP signature scores (o). (p) Patient-level EMP and LMP scores in localized and metastatic samples from GSE141445. Signature scores were averaged across malignant epithelial cells within each patient, and each point represents one patient. (q) Schematic of paired primary-prostate and BM single-cell profiling in the Extended Mets cohort (n = 2), with a UMAP of all major cell types shown on the right. (r) Heatmap showing marker-gene expression across major cell types in the Extended Mets samples. (s-t) UMAP visualization of epithelial cells from the Extended Mets samples, colored by subclusters identified by the Louvain algorithm (s) or annotated subtypes (t). (u) Heatmap shows the expression levels (z-score) of selected phenotype-related genes across epithelial subclusters from Extended Mets samples. (v) Inferred copy-number-variation profiles of epithelial cells in the Extended Mets samples. Columns represent genomic regions, rows represent reference and epithelial cells, and the annotation bar indicates epithelial subclusters. (w) Violin plot shows the EMP scores in malignant epithelial cells from Extended Mets samples. (x) UMAP plots display the expression pattern of EMP marker genes in malignant epithelial cells across individuals in the Extended Mets samples, colored by normalized expression levels. (y) Violin plots show the LMP scores within malignant epithelial subclusters from two individuals in the Extended Mets samples, colored by the average score of each subcluster. *Comparisons in panel l were performed using two-sided unpaired Student’s t-tests, whereas comparisons in panel p were performed using one-sided unpaired Student’s t-tests. P values were adjusted using the Benjamini–Hochberg method.*

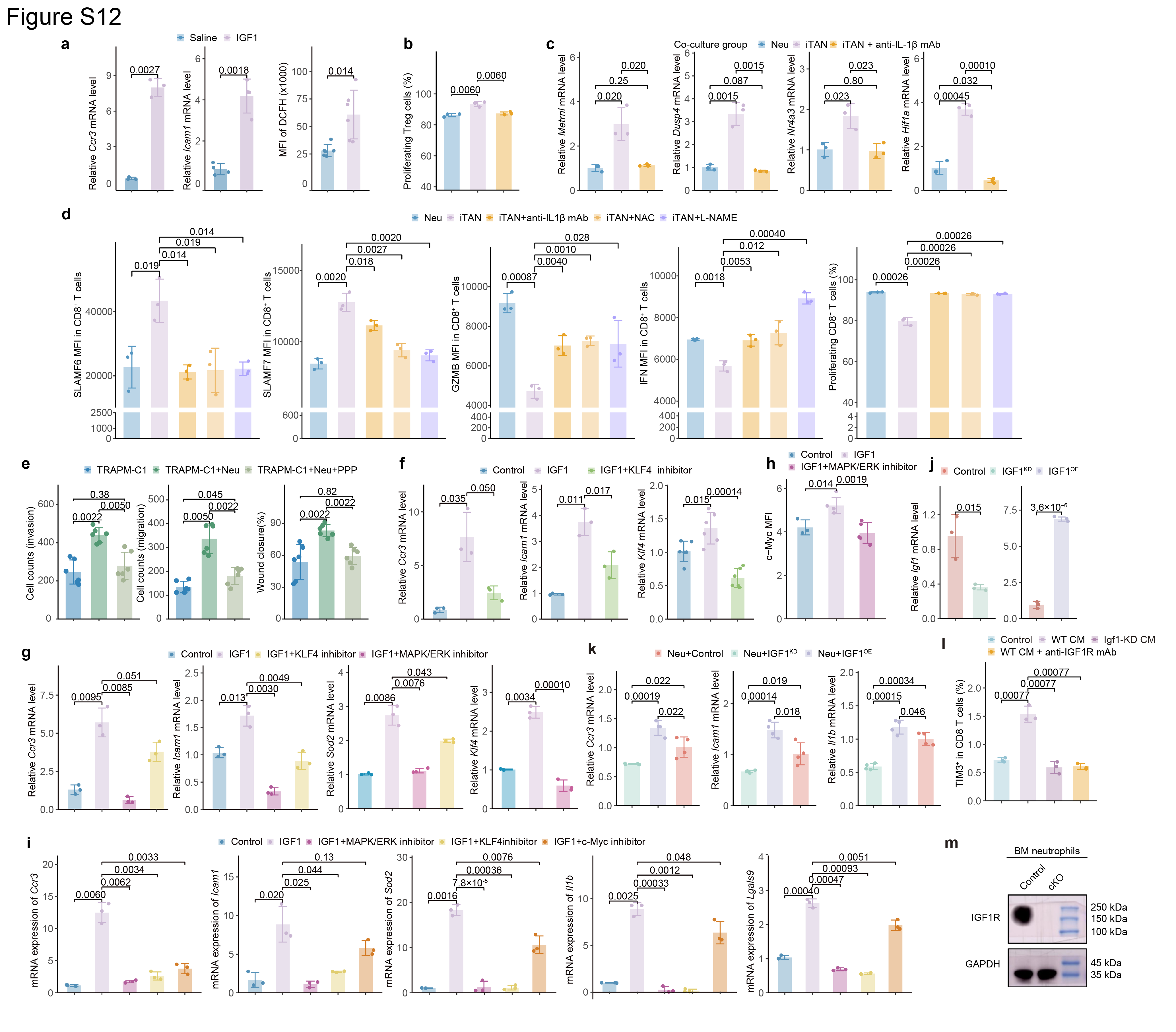

**Fig. S12. Molecular mechanisms and functional consequences of IGF1-induced TAN polarization.**

(a) Relative *Ccr3* and *Icam1* mRNA expression and DCFH fluorescence MFI in vehicle-treated control and IGF1-stimulated mouse BM-derived neutrophils. For *Ccr3*, n = 3 biologically independent samples per group; for *Icam1*, n = 4; for DCFH, n = 6. (b) Percentage of proliferating Tregs after co-culture with unstimulated neutrophils, IGF1-induced TANs, or IGF1-induced TANs treated with an IL-1β-neutralizing antibody; n = 3 biologically independent samples per group. (c) Relative *Metrnl*, *Dusp4*, *Nr4a3* and *Hif1a* mRNA expression in CD8 T cells co-cultured with unstimulated neutrophils, IGF1-induced TANs, or IGF1-induced TANs in the presence of an IL-1β-neutralizing antibody; n = 3 biologically independent samples per group. (d) SLAMF6, SLAMF7, GZMB and IFNG MFIs and percentage of proliferating CD8 T cells after co-culture with neutrophils, IGF1-induced TANs, or IGF1-induced TANs treated with an IL-1β-neutralizing antibody, N-acetylcysteine (NAC) or L-NAME; n = 3 biologically independent samples per group. (e) Migration, invasion and wound-closure assays of TRAMP-C1 cells cultured alone or co-cultured with neutrophils, with or without PPP; n = 6 biologically independent experiments per group. (f) Relative *Ccr3*, *Icam1* and *Klf4* mRNA abundance after IGF1 stimulation with or without KLF4 inhibition. For *Ccr3* and *Icam1*, n = 3 biologically independent samples per group; for *Klf4*, n = 6. (g) Relative *Ccr3*, *Icam1*, *Sod2* and *Klf4* mRNA abundance after IGF1 stimulation with KLF4 or MAPK/ERK pathway inhibition; n = 3 biologically independent samples per group. (h) c-MYC MFI after IGF1 stimulation with or without MAPK/ERK pathway inhibition. Control, n = 3; IGF1, n = 4; IGF1 plus MAPK/ERK inhibitor, n = 5 biologically independent samples. (i) Relative *Ccr3*, *Icam1*, *Sod2*, *Il1b* and *Lgals9* mRNA abundance after IGF1 stimulation with MAPK/ERK, KLF4 or c-MYC inhibition; n = 3 biologically independent samples per group. (j) Relative *Igf1* mRNA abundance in TRAMP-C1 cells with wild-type, knocked-down or overexpressed Igf1; n = 3 biologically independent experiments per group. (k) Relative *Ccr3*, *Icam1* and *Il1b* mRNA abundance in neutrophils exposed to conditioned medium (CM) from TRAMP-C1 cells with wild-type, knocked-down or overexpressed Igf1; n = 4 biologically independent experiments per group. (l) Frequencies of TIM3-positive CD8 T cells after treatment with matched blank medium (Control), wild-type TRAMP-C1 CM, Igf1-knockdown CM or wild-type CM plus an IGF1R-blocking antibody in vivo; n = 3 mice per group. Data in bar plots are presented as mean ± s.d. (m) Immunoblot analysis of IGF1R protein expression in BM neutrophils from control and cKO mice. GAPDH served as a loading control. Molecular mass markers are indicated in kDa. *Comparisons were performed using two-sided Student’s t-tests and P values were adjusted using the Benjamini-Hochberg method.*

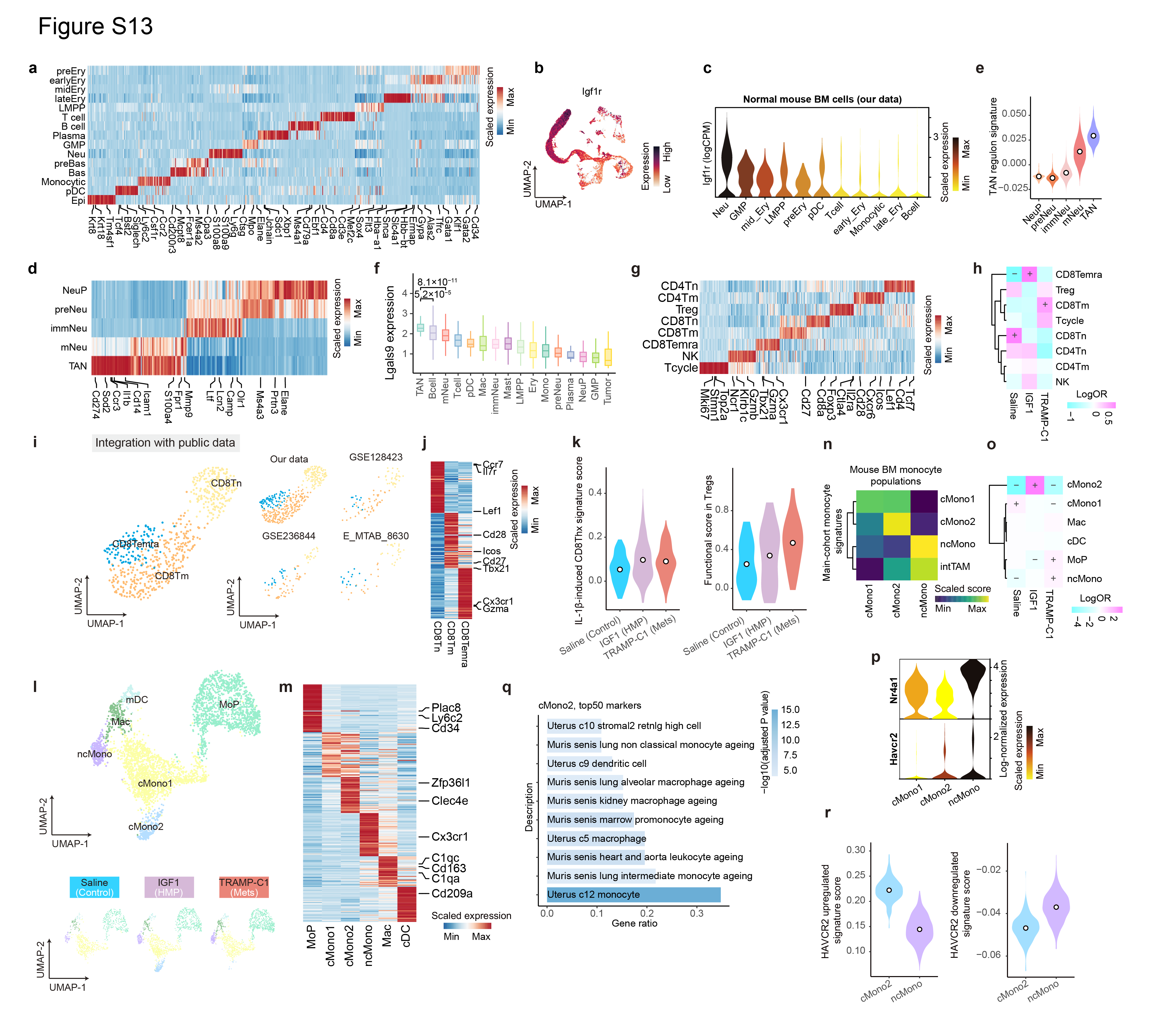

**Fig. S13. Cross-species validation of neutrophil, CD8 T cell and monocyte states in mouse BM.**

(a) Heatmap showing marker-gene expression across major mouse bone marrow (BM) cell types. Color intensity represents row-scaled mean expression. (b) Feature plot showing *Igf1r* expression across all mouse BM cells. (c) Violin plots showing *Igf1r* expression across major mouse BM cell types; violin color indicates scaled mean expression. (d) Heatmap showing marker-gene expression across mouse neutrophil subtypes. (e) Violin plots showing TAN-regulon-signature scores across mouse BM neutrophil subtypes. The signature was derived from the human main cohort, and violin color indicates subtype. (f) *Lgals9* expression across major mouse BM cell types; expression was calculated among *Lgals9*-positive cells. (g) Heatmap showing marker-gene expression across mouse CD8 T cell subtypes. (h) Differential abundance of CD8 T cell subtypes across saline-control, IGF1-induced HMP and TRAMP-C1 metastatic groups. Color indicates the log odds ratio, and “+” or “−” denotes significant enrichment or depletion, respectively. (i) UMAP of mouse CD8 T cells integrated with public mouse BM datasets, colored by subtype, with dataset-specific embeddings shown on the right. (j) Heatmap showing marker-gene expression across integrated mouse CD8 T cell subtypes. (k) IL-1β-induced CD8Thx signature scores in integrated CD8Tm cells, and functional scores in integrated Tregs across saline-control, IGF1-induced HMP and TRAMP-C1 metastatic groups. (l) UMAP of mouse monocytic subtypes, colored by subtype, with group-specific embeddings shown below. (m) Heatmap showing genes differentially expressed across mouse monocytic subtypes. (n) Heatmap showing the signature scores of human monocyte states when projected onto mouse monocyte populations. (o) Differential abundance of mouse monocytic subtypes across experimental groups, displayed as in panel h. (p) Violin plots showing *Nr4a1* and *Havcr2* expression across mouse monocytic subtypes. (q) Over-representation analysis of DEGs of cMono2 using the signature collections from MSigDB. The x-axis indicates the gene ratio, and bar color indicates −log10(adjusted *P* value). (r) Scores of HAVCR2-upregulated and HAVCR2-downregulated gene signatures in cMono2 and ncMono cells. *Exact P values and statistical tests are shown in the panels; Student’s t-test P values were adjusted using the Benjamini-Hochberg method. Differential abundance in panels h and o was assessed using two-sided Fisher’s exact tests with Benjamini-Hochberg adjustment for multiple comparisons.*

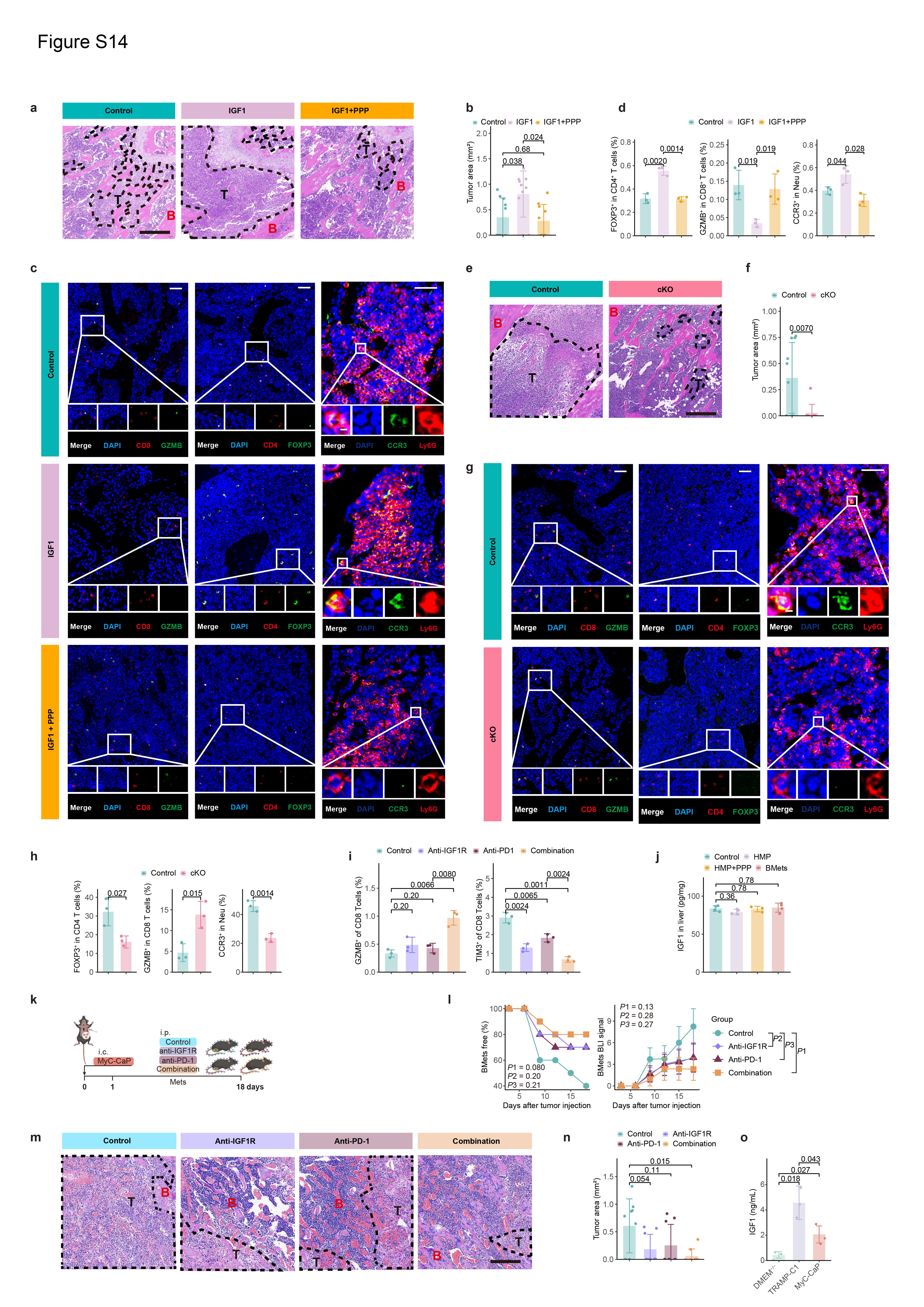

**Fig. S14. Histological and immune validation of IGF1R targeting in bone-metastasis models.**

(a) Representative haematoxylin-and-eosin-stained bone sections from Control, IGF1 and IGF1 plus PPP groups after tumor challenge. For panels a–d, Control denotes mice receiving matched IGF1 and PPP vehicles. Dashed lines outline tumor regions (T); B denotes bone. Scale bar, 400 µm. (b) Quantification of tumor area in the indicated groups; n = 10 mice per group. (c) Representative bone immunofluorescence images stained for DAPI and CD8/GZMB, CD4/FOXP3 or CCR3/Ly6G in Control, IGF1 and IGF1 plus PPP groups. Scale bars: for CD8/GZMB, CD4/FOXP3: 40 µm; for CCR3/Ly6G: 25 μm. (d) Quantification of FOXP3-positive CD4 T cells, GZMB-positive CD8 T cells and CCR3-positive neutrophils in the indicated groups; n = 3 mice per group. (e) Representative haematoxylin-and-eosin-stained bone sections from littermate control and neutrophil-specific *Igf1r* conditional-knockout mice (cKO) after tumor challenge. Dashed lines outline tumor regions (T); B denotes bone. Scale bar, 200 µm. (f) Quantification of tumor area in control and cKO mice. n = 10 mice per group. (g) Representative bone immunofluorescence images from control and cKO mice stained for DAPI and CD8/GZMB, CD4/FOXP3 or CCR3/Ly6G. Scale bars are the same as panel c. (h) Quantification of FOXP3-positive CD4 T cells, GZMB-positive CD8 T cells and CCR3-positive neutrophils in control and cKO mice; n = 3 mice per group. (i) Frequencies of GZMB-positive and TIM3-positive CD8 T cells after Control, anti-IGF1R, anti-PD-1 or combined anti-IGF1R plus anti-PD-1 treatment; n = 3 mice per group. (j) IGF1 concentrations in liver samples from control mice, mice at the HMP stage, PPP-treated mice at the HMP stage, and mice with bone metastases (BMets). Liver samples from the control, HMP, and PPP-treated HMP groups were collected from the mice used in Fig. 4a, whereas liver samples from the BMets group were collected from the Control mice used in Fig. 4g. Control, n = 4 mice; HMP, n = 4 mice; PPP-treated HMP, n = 3 mice; BMets, n = 4 mice. (k) Experimental design of an independent MyC-CaP bone-metastasis model evaluating Control, anti-IGF1R, anti-PD-1 or combined treatment. (l) Bone-metastasis-free curves and longitudinal bone BLI signals across treatment groups. Per-mouse AUCs of ln(signal + 1)-transformed serial BLI measurements were compared; n = 10 mice per group. *P*1, *P*2 and *P*3 denote Control versus Combination, Control versus anti-IGF1R and Control versus anti-PD-1, respectively. (m) Representative haematoxylin-and-eosin-stained bone sections from Control, anti-IGF1R-, anti-PD-1- and combination-treated mice (BMets established using MyC-CaP cell line). Dashed lines outline tumor regions (T); B denotes bone. Scale bar, 400 µm. (n) Quantification of tumor area in the indicated treatment groups; each dot represents one mouse. n = 10 mice per group. (o) IGF1 concentrations in matched serum free DMEM control medium and conditioned medium (CM) from TRAMP-C1 and MyC-CaP cells; n = 3 biologically independent samples per group. *Data in bar plots are presented as mean ± s.d. Bone-metastasis-free curves were compared using log-rank tests; longitudinal BLI data were compared using a two-sided Welch's t-test. Other comparisons used two-sided Student’s t-tests with Benjamini-Hochberg-adjusted P values.*

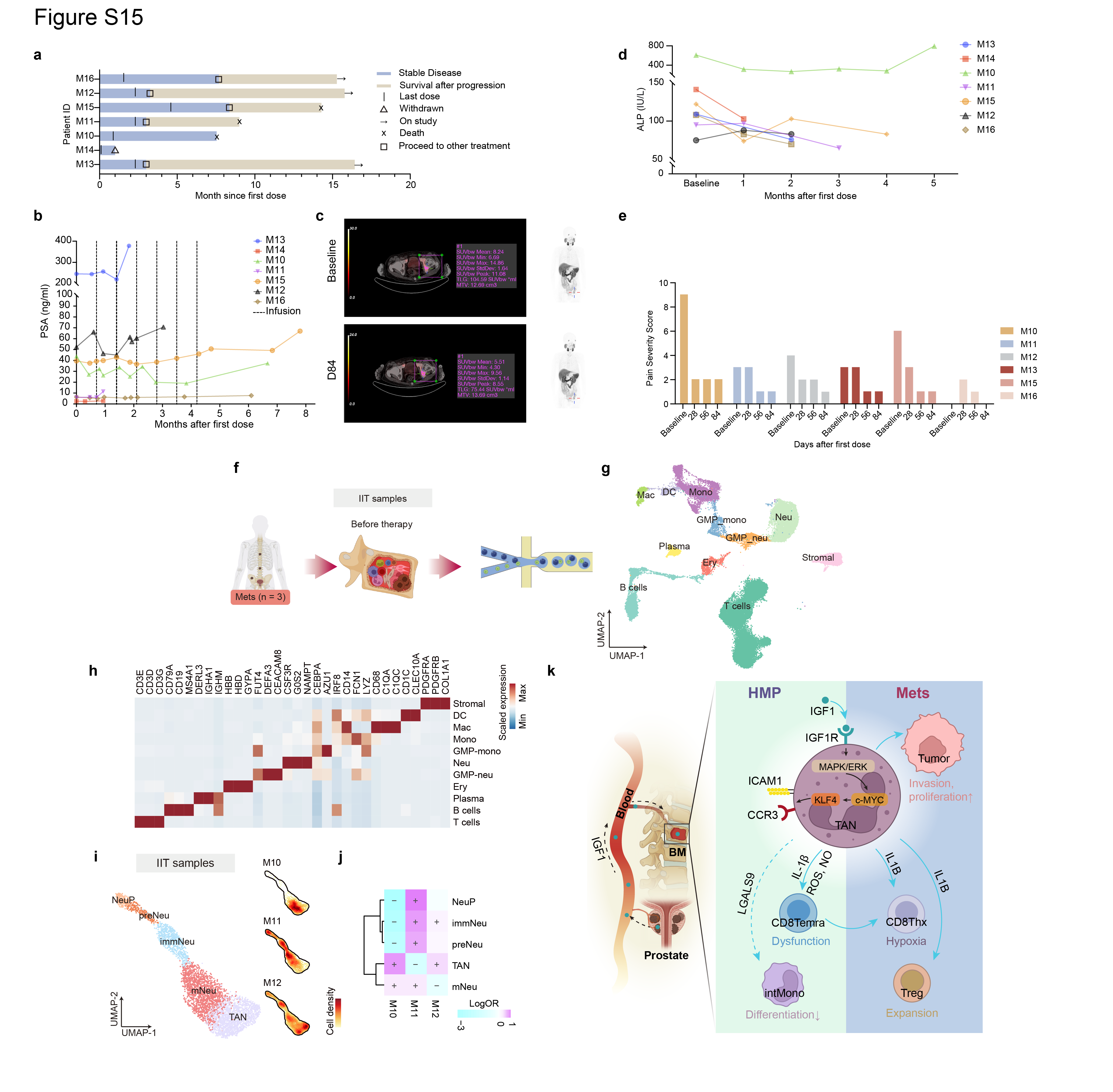

**Fig. S15. Clinical responses to combined IGF1R and PD-1 blockade and the pretreatment bone marrow environment.**

(a) Swimmer plot showing treatment duration, stable disease, survival after progression and clinical events for each participant receiving teprotumumab plus tislelizumab. (b) Longitudinal serum prostate-specific antigen (PSA) concentrations after the first dose; dashed vertical lines indicate drug infusions. (c) Representative PET/CT images at baseline and day 84, with lesion standardized uptake values shown. (d) Longitudinal serum alkaline-phosphatase (ALP) concentrations after the first dose. (e) Patient-reported pain-severity scores at baseline and during treatment, colored by patient identity. n = 6 evaluable participants. (f) Workflow for pretreatment bone marrow (BM) collection and single-cell RNA sequencing from three participants in the investigator-initiated trial. (g) UMAP of all pretreatment BM cells, colored by major cell type. (h) Heatmap showing marker-gene expression across the major BM cell types. Color intensity represents row-scaled mean expression. (i) UMAP of pretreatment BM neutrophil subtypes with patient-specific cell-density maps for M10–M12. (j) Differential abundance of neutrophil subtypes across patients M10–M12. Color indicates the log odds ratio, and “+” or “−” denotes significant enrichment or depletion, respectively. (k) Proposed model of prostate tumor-to-BM communication. Tumor-associated systemic IGF1 is increased during prostate cancer progression and can act on IGF1R-expressing BM neutrophils, inducing MAPK/ERK-c-MYC-KLF4-dependent TAN differentiation. During the HMP stage, TAN-derived IL-1β promotes a pre-exhausted CD8Temra state, whereas LGALS9-HAVCR2 signaling is associated with altered monocyte differentiation. In established bone metastasis, TANs promote CD8Thx hypoxia and Treg expansion, thereby supporting tumor invasion and proliferation. intMono, intermediate monocytes. *Differential abundance in panel j was assessed using two-sided Fisher’s exact tests with Benjamini-Hochberg adjustment for multiple comparisons.*

Table S1. Clinical metadata for study cohorts and donors (Provided as a separate file)

Table S2. Gene sets and signatures used in this study (Provided as a separate file)

Table S3. Gene-set enrichment results for meta-programs (Provided as a separate file)

Table S4. Predicted transcription factor-binding sites in the *Klf4* promoter (Provided as a separate file)

Table S5. Patient characteristics of the IIT cohort (Provided as a separate file)

Table S6. Treatment-related adverse events in the IIT cohort (Provided as a separate file)

Table S7. Summary of treatment responses in the IIT cohort (Provided as a separate file)

Table S8. Changes in pain severity at follow-up according to baseline pain-severity level in the IIT cohort (Provided as a separate file)

Table S9. Information on public datasets used in this study (Provided as a separate file)

Table S10. Primer sequences used for qPCR and ChIP–qPCR (Provided as a separate file)

Table S11. Antibodies used for flow cytometry (Provided as a separate file)
